# The Nature of Centromeric Repeat Turnovers in the genus *Arabidopsis*

**DOI:** 10.64898/2026.09.10.749358

**Authors:** Anna Glushkevich, Robin Burns, Maria Vasilarou, Estela Perez-Roman, Uliana Kolesnikova, Akash Chandra Parida, Laura Avila Robledillo, Alexander Mackintosh, Ryo Suda, Laura Steinmann, Ursula Pfordt, Karl Schmid, Ute Kraemer, André Marques, Takashi Tsuchimatsu, Tiina M. Mattila, Alexandros Bousios, Ian Henderson, Alison Scott, Polina Novikova

**Affiliations:** Department of Chromosome Biology, Max Planck Institute for Plant Breeding Research, Carl-von-Linne-Weg 10, Cologne, 50829, Germany; Department of Plant Sciences, University of Cambridge, Cambridge, UK; School of Life Sciences, University of Sussex, UK; Department of Molecular Genetics and Physiology of Plants, Ruhr University Bochum, Universtitaetsstrasse 150 ND3/30 Box 44, D-44810 Bochum, Germany; Department of Bioinformatics and Genetics, Swedish Museum of Natural History, Stockholm, Sweden; Department of Ecology and Genetics, Evolutionary Biology Centre, Uppsala University, Uppsala, Sweden; Department of Biological Sciences, Graduate School of Science, The University of Tokyo, Tokyo, Japan; Institute of Plant Breeding, Seed Science and Population Genetics, University of Hohenheim, Stuttgart, Germany; Cluster of Excellence GreenRobust, University of Hohenheim, Stuttgart, Germany; Ecology and Genetics Research Unit, University of Oulu, PO Box 3000, 90014, Oulu, Finland; Department of Plant Biotechnology and Bioinformatics, Ghent University, Ghent, Belgium; VIB-UGent Center for Plant Systems Biology, Ghent, Belgium

## Abstract

Centromeres are critical for accurate segregation of chromosomes and are often composed of megabases of tandemly arranged satellite repeats. Yet, despite their conserved function, the DNA sequence of centromeres is, paradoxically, rapidly evolving. To understand the nature of centromeric sequence turnover, we assembled 417 centromeres from nine species representing the entire *Arabidopsis* genus. In the genus, centromeres are formed by four main satellite repeats with homologous sequences forming central arrays and minor repeat types. We identify the ancestral centromeric repeat type for the *Arabidopsis* genus and three independent turnovers to different repeats: (1) a complete turnover in *A. thaliana*, (2) a turnover of seven out of the eight centromeres in the ancestor of *A. cebennensis* and *A. pedemontana*, (3) turnovers of three to five centromeres in the genomes of *A. halleri* and *A. lyrata*. Most centromeric repeats are also present throughout the genome with shared syntenic locations between *Arabidopsis* species, and some being similar to parts of transposable elements and genes, suggesting that centromeric repeats originate outside of the centromeres. In allotetraploid *A. suecica* we find that the repeats from centromeric arrays on one subgenome can transpose and invade the other, likely via transposable element activity. All four main centromeric repeats can recruit the CENH3 (CENP-A) histone variant, however, when a new repeat successfully proliferates in an old array, CENH3 is primarily recruited to the new array marking functional take over. Complete centromeric turnovers occurred in species that experienced severe bottlenecks in their evolutionary history, where new centromeric alleles may have been fixed by genetic drift. Yet, we also observe segregation distortion between two different centromeric repeat types in *A. lyrata*, suggestive of centromere drive and we propose that both drift and drive contribute to centromeric repeat turnover. Together, our findings reveal the origin of centromeric repeats, mechanisms of repeat proliferation and spread, and the evolutionary dynamics of centromeric repeat turnovers in the *Arabidopsis* genus.

## Introduction

During cell division chromosomes segregate to opposite poles as spindle microtubules attach via kinetochores to the centromeric regions of the chromosomes and pull them apart. Centromeres are defined epigenetically by a special variant of histone 3, CENH3/CENP-A^1^, binding DNA underlying the centromere which can be made of transposable elements or tandem repeat arrays. While tandem repeats are highly prevalent at centromeres, suggesting their functional role, the homology of those repeats is generally only conserved between closely related species^2–6^. An exception to this trend is found in holocentric *Rhynchospora* species that show high conservation of centromeric satellite repeats despite 40 million years of evolution accompanied by extensive karyotypic changes^7^. Long-read genome sequencing technologies have enabled the complete assembly of the tandem repeat arrays of many eukaryote centromeres, including non-model systems. It remains unclear how centromeric function is maintained despite the rapid evolution of the underlying DNA, and what mechanisms generate extreme divergence of centromeric sequences that often exceed the genome-wide divergence.

The asymmetric cell division during female meiosis, when one large egg is produced instead of four equal-sized ones, creates a competitive environment for chromosomes to be inherited in the next generation^8,9^. The stronger the centromere binds to the kinetochore, the more chances it has to be transmitted to the egg^10^. A genetic conflict between the DNA with high propensity to be inherited, so-called “selfish elements” and the proteins binding centromeric regions, also evolving to restore the inheritance balance, has been suggested to drive the rapid evolution of centromeres^5,11^. The centromeric repeats usually are of length roughly corresponding to a nucleosome unit (ca. 150-200 bp) which probably allows to bind CENH3 in the most efficient way^5,7,12^, especially in homogenic repeat arrays^13^. CENH3 shows higher sequence variability when compared to other histone H3 proteins in eukaryotes^14,15^. The C-terminus of CENH3 contains the histone fold used in kinetochore localization and shares homology with other histone proteins^14,15^, however, the N-terminus does not share sequence homology to canonical histone proteins, and while the function remains to be resolved, through tail-swap experiments, is known to be dispensable during mitosis but indispensable for meiosis-specific centromere assembly^16^. Other selfish elements, like transposons (*e.g.*, ATHILA in *A. thaliana*^17^) that evolve to preferentially integrate into the centromere and therefore ensure their transmission, disrupt the order and probably lower the efficiency of CENH3 binding^18^. This process is counteracted by the recombination machinery associated with kinetochore promoting homogenisation of the centromeric repeat arrays^17–19^. Proliferation of repeats pushes the older repeat arrays to the sides of the centromere, where the arrays shrink and degrade^20^. Centromeric repeat arrays exhibit higher point mutation rates than the rest of the genome and undergo non-allelic gene conversion^21^, sufficient to generate substantial within-species diversity and result in the formation of higher-order structures^18^, repeats of large multimonomeric units. However, frequent repeat turnover of the actively CENH3 binding part of centromeres has been observed in many species^6^ and it is not known if the mutation and recombination processes would be sufficient to generate an array of different repeats or if a new substrate is required for turnovers.

Phylogenetically complete sampling of centromeric diversity from a genus with multiple repeat turnovers can help to increase understanding on the nature of centromere evolution. We use the *Arabidopsis* genus as a tractable system for studying genetic turnovers of repeats underlying centromeres due to its well studied evolutionary history of roughly six million years^22–25^. Europe is the center of genetic diversity for the genus from where it spread to eastern Eurasia, Africa and North America^26–29^. *A. thaliana*, *A. lyrata*, *A. arenosa*, and *A. halleri* are the most widespread^22,30^, while *A. cebennensis*, *A. pedemontana*, and *A. croatica* have restricted distributions and lower genetic diversity^31,32^. Moreover, two allopolyploid species in the genus, *A. suecica* (hybrid between *A. thaliana* and *A. arenosa*) and *A. kamchatica* (hybrid between *A. lyrata* and *A. halleri*) allow to trace the dynamics of the centromeric repeats after the hybridization and the merger of the subgenomes.

Previous studies in *Arabidopsis* showed no overlap in centromeric repeat types between *A. thaliana* and *A. lyrata*^17,33,34^, which diverged 5-10 million years ago^22,24,35^. Apart from *A. thaliana* centromeric repeats AthCEN178 (former naming pAL1) and AthCEN159, three types of repeats differing in sequence and ranging in length (159-179 bp) were found in the *Arabidopsis* genus by cloning and cytological studies: AlyCEN179 (pAa), AlyCEN168 (pAge1) and AlyCEN176 (pAge2). These three repeats form centromeric arrays of *A. lyrata* and *A. halleri*, while *A. arenosa* centromeres formed by AlyCEN179 only^33,36,37^. Not much is known about the centromeric composition of endemic *Arabidopsis* species (*A. cebennensis*, *A. pedemontana* and *A. croatica*) apart from that AlyCEN168 is present on the centromere of chromosome 5 of *A. cebennensis*^33,36^. To what extent the diversity of centromeric repeats is affecting CENH3 evolution within the Arabidopsis genus is also still unclear. Five distinct protein variants of CENH3 were observed within 66 *A. thaliana* genomes all composed of mainly centromeric repeats AthCEN178 repeat type^17^. Despite rapid evolution of centromeric arrays and CENH3 variants, CENH3 from distant species can bind *A. thaliana* centromeric arrays at the same location of the most homogeneous and therefore youngest parts of arrays^38^. The centromeres on both sub-genomes of allotetraploid *A. suecica*, which are *A. thaliana* with AthCEN178 centromeres and *A. arenosa* with AlyCEN179 centromeres, incorporate CENH3 encoded by both expressed homeologs^39^. While Arabidopsis centromeres are mostly satellite repeat based, the repeat array is often interrupted by transposons, which are not as effective at attracting CENH3 compared to satellite repeats^17^. *A. thaliana* has only one centrophylic TE type, Ty3 transposon Athila, especially Athila5^4,13^ while *A. lyrata* centromeres are also enriched in Ty1 elements in addition to Athila5, and have higher, more than doubled, overall retrotransposon content compared to *A. thaliana*^4^.

To understand the centromeric repeat turnover, we generated T2T assemblies of all species in the *Arabidopsis* genus and analysed the centromere diversity patterns from the *Arabidopsis* species and the outgroup, *Capsella orientalis*. We ask if there was a prevalent centromeric repeat in the ancestral lineage of the genus *Arabidopsis* and how many turnovers occurred during the evolutionary history of at five to ten million years. Do the four described centromeric repeat types in *Arabidopsis* share their origin or did they evolve from different founder events? How did the new centromeric repeats form and then spread from a short array to a complete centromere and then turn over all the centromeres into a new repeat type? Finally, we ask which evolutionary forces, neutral demographic changes or an active meiotic drive leaving selection signatures on the genomes, lead to the observed diversity of centromeres within and between *Arabidopsis* species.

## Results

We obtained 54 genomes representing all *Arabidopsis* species (*A. thaliana*, *A. arenosa*, *A. croatica*, *A. cebennensis*, *A. pedemontana*, *A. lyrata*, *A. halleri*, *A. suecica* and *A. kamchatica*) and the outgroup genome of *Capsella orientalis* (Supplementary Data 1). We used TRASH^40^ to identify individual tandem repeats in the centromeric regions using consensus sequences of known centromeric repeat types AthCEN178, AlyCEN179, AlyCEN186 and AlyCEN176 as a template. We identified centromeres by the presence of dense, long (hundreds kb) repeat arrays of centromeric repeat types known from the previous studies (AthCEN178, AlyCEN179, AlyCEN186, AlyCEN176, AthCEN159)^17,33^. More than 90% (378 out of 417) of centromeres were assembled as a single contig showing their completeness, including both the shortest and the longest centromeres (Fig. 1, contig breaks represented by black tick marks). Overall, we identified 5.3 million monomers with 1.5 million unique sequences. *Arabidopsis* centromeres range from 0.8 to 6.1 Mb in length and are shorter than those of *C. orientalis* ranging from 5 to 8.5 Mb (Supplementary Fig. S1a). Centromeric arrays contribute to total chromosomal length in *A. lyrata*, the species with most of the assembled genomes in our study (Supplementary Fig. S1b, correlation p-val = 6.4e-05), however when the length of a centromeric array is excluded from the corresponding chromosome length that weak correlation disappears, suggesting there is no functional dependence between the centromeric array length and the chromosome length (Supplementary Fig. S1c). Centromere locations are shared between *Capsella* and *Arabidopsis* (Supplementary Fig. S2), including syntenic centromeres of *A. thaliana*, which are centromeres on chromosomes 1 (syntenic to chromosome 1 in *A. lyrata*), 2 (syntenic to chromosome 3 in *A. lyrata*), 3 (syntenic to chromosome 5 in *A. lyrata*) 4 (syntenic to chromosome 6 in *A. lyrata* from one side) and 5 (syntenic to chromosome 7 in *A. lyrata* from one side)^33^.

**Figure 1.**
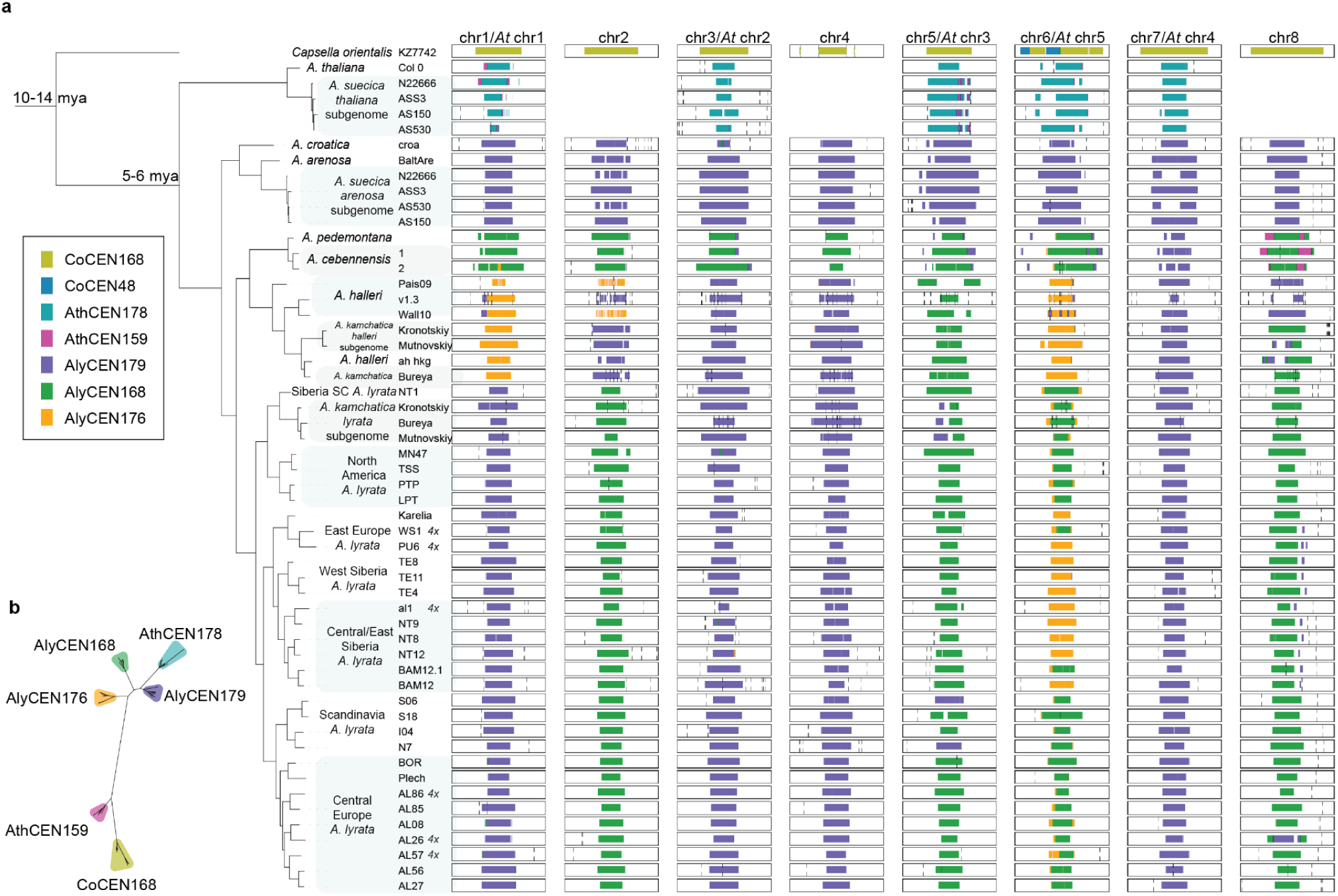
Centromere composition across the genus *Arabidopsis* with *Capsella orientalis* as outgroup. (a) Rectangular blocks represent a 10 Mb region, containing the centromere. Syntenic centromeres are shown between *A. thaliana* and other *Arabidopsis* based on the genomes synteny (Supplementary Fig. 2) and previous study^33^. Centromeric repeat arrays are colored by repeat type. Black vertical lines show breaks between the assembled contigs. The tree on the left shows the relatedness of the samples based on 36,000 orthologous protein sequences. (b) Phylogenetic tree based on monomer alignments of *Arabidopsis* and *Capsella* centromeric repeat types. Highlighting corresponds to repeat types in (a).

We classified the repeat types using PCA on 5-mer monomer composition (Supplementary Fig. S3, an example for *A. lyrata*). *Arabidopsis* has four major repeat types forming centromeric arrays: AlyCEN179, AlyCEN168, AlyCEN176, AthCEN178. The sequences of the repeats are alignable, suggesting their shared ancestry (Fig. 1b, Supplementary Fig. S4). A complete turnover of the repeat types occurred during the evolutionary history of *Capsella* and *Arabidopsis* as none of the major repeat types are shared between them. Genus-level resolution in *Arabidopsis* allows us to infer the following history of turnovers:

(1) AlyCEN179 repeat type is shared by all of the *Arabidopsis* species apart from *A. thaliana* and is probably the closest to the ancestral repeat type (Fig. 1a).
(2) The centromeres of *A. arenosa* and *A. croatica* remained AlyCEN179-type (Fig. 1, Supplementary Fig. S5), with minor presence of AlyCEN168 shared with *A. cebennensis*, *A. pedemontana*, *A. halleri* and *A. lyrata*.
(3) *A. thaliana* centromeres turned over into a AthCEN178 repeat type. AthCEN178 and AlyCEN179 repeats are genetically closest repeats with a relatively frequent intermediate class in *A. thaliana* (Fig. 1,2, Supplementary Fig. S6). A minor presence of AthCEN159 in *A. thaliana* centromeres is shared only with centromeres of *A. cebennensis* and *A. pedemontana*, two closely related, endemic and endangered species in France and Italy^24,31^. AthCEN159 is close to the *Capsella* centromeric repeat (Co_168) with the sequence identity between the types from 60 to 68% while having no identity to the rest of the *Arabidopsis* repeats^34^ (Fig. 1b, Supplementary Fig. S4).
(4) The centromeres of *A. cebennensis* and *A. pedemontana* are turning over into AlyCEN168 repeat type (Supplementary Fig. S7, repeats are chromosome-specific). This does not include the centromere on chromosome 7, which remains the ancestral AlyCEN179 type in all species, except for the syntenic centromere in *A. thaliana* (Supplementary Fig. S8).
(5) Centromeres of *A. halleri* and *A. lyrata* are the most diverse in repeat types. Apart from AlyCEN179 and AlyCEN168, a new repeat-type, AlyCEN176, can also take over an entire centromere.

**Figure 2.**
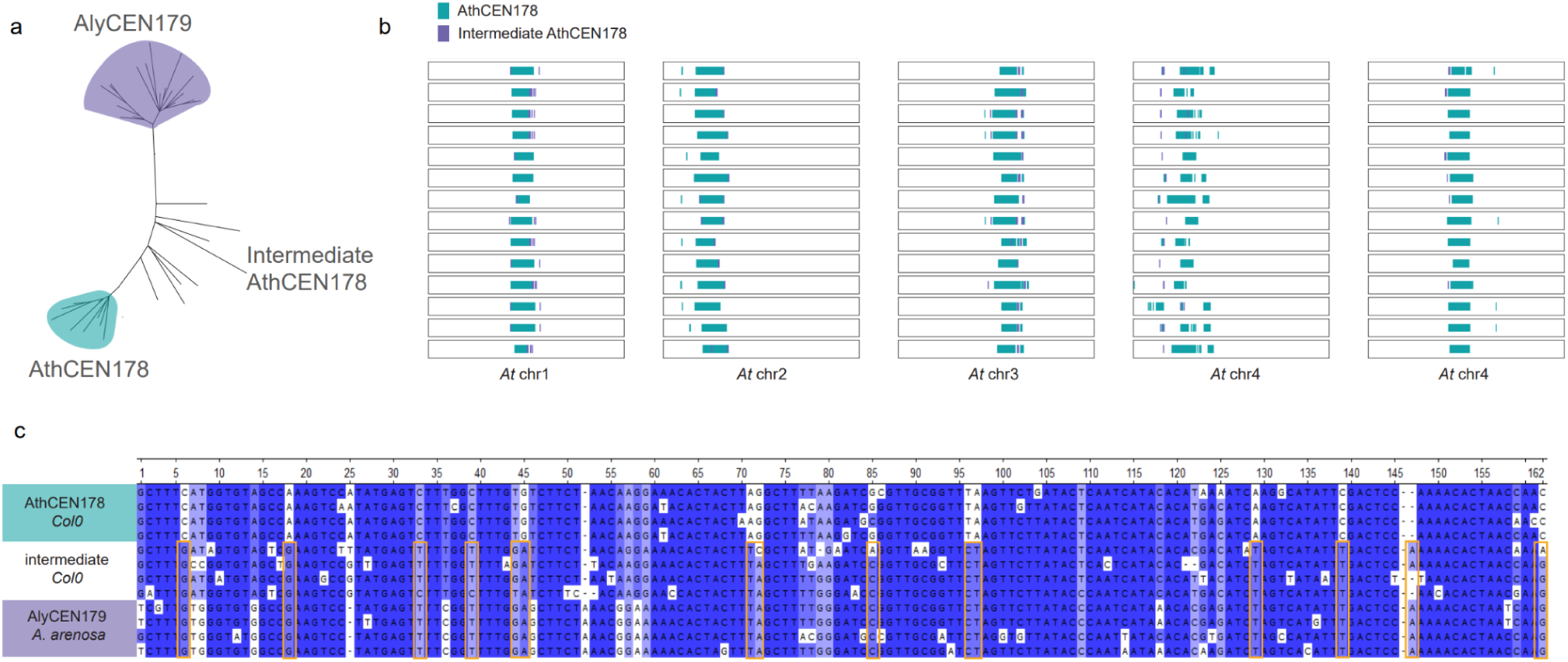
The *A. thaliana*-specific AthCEN178 evolved from the ancestral *Arabidopsis* repeat AlyCEN179 (pAa) repeat through intermediate types. a) Unrooted maximum-likelihood tree of sampled monomers from *A. thaliana* and *A. arenosa*. b) BLAST hits of main centromeric repeat types on 15 *A. thaliana* assemblies (all 66 used in the study in the Supplementary Fig. S6). Blue - BLAST hit of AthCEN178 repeat, purple - BLAST hit of AlyCEN179. Each block represents the whole chromosome. c) Multiple sequence alignment of sample monomers from Col-0 and Baltic *A. arenosa* assembly. Sample AthCEN178 repeat from Col-0 is used as a reference and mismatches with this repeat are highlighted. Some SNPs are private to AlyCEN179 from *A. arenosa* while others are shared with AlyCEN179-like repeats from *A. thaliana* (marked with orange boxes).

Different centromeric repeat arrays segregate within *A. lyrata* and *A. halleri*. Chromosome 2 of *A. halleri* can be predominantly AlyCEN176 (orange) or AlyCEN179 (purple) (Fig. 1). Chromosome 5 of *A. lyrata* segregates AlyCEN168 (green) and AlyCEN179 (purple), the latter likely ancestral (Supplementary Fig. S9, Supplementary Data 2), but only present at high frequencies in Scandinavian *A. lyrata* (Supplementary Fig. S11). Chromosome 6 of *A. lyrata* segregates AlyCEN168 (green) or AlyCEN176 (orange) (Fig. 1). The likely evolution of turnovers of chromosome 6 repeat types is from ancestral AlyCEN179 to AlyCEN168 (green) in the common ancestor of *A. cebennensis* and *A. pedemontana* and AlyCEN176 (orange) in common ancestor of *A. halleri* and *A. lyrata* (Supplementary Fig. S10, Supplementary Data 2) The AlyCEN168 (green) in *A. lyrata* are likely also derived as they are different from *A. cebennensis* and *A. pedemontana* (Supplementary Fig. S9, Supplementary Data 2).

Two *A. lyrata* plants sampled at the same location BAM12 and BAM12.1 share repeat types in all the centromeres, reaching 95% of repeat libraries sharing on the centromere of chromosome 1 (Supplementary Data 2), but not at the centromere of chromosome 6, where BAM12 has only AlyCEN176 repeats and BAM12.1 has AlyCEN168 centromeric repeat arrays (Fig. 1a). Similar patterns when centromere haplotypes on some chromosomes can be shared while others are different were observed in closely related *A. thaliana* accessions^17^. Higher-order patterns of centromeric structure can be shared even between species, for example, in between *A. cebennensis* and *A. pedemontana* (Supplementary Fig. S12) diverged about 170 kya^31^.

### Ancestral repeat-type

The AlyCEN179 repeat type centromeres are deeply shared by all the *Arabidopsis* species except *A. thaliana* (Fig. 1). Moreover, *A. thaliana* has centromeric repeat monomers that are intermediate between the most common type of AthCEN178 and AlyCEN179 (Fig. 2a,b). These intermediate repeats are located in the end of centromeric arrays and shared in the same locations between diverse accessions of *A. thaliana* which also suggests they are old (Fig. 2b). The AlyCEN179 repeats are sometimes found at the ends of the centromeres formed by other repeat arrays in *A. cebennensis*, *A. pedemontana*, *A. lyrata* and *A. halleri* (Fig. 1). All four major centromeric repeat types (AlyCEN179, AlyCEN168, AlyCEN176, AthCEN178) are present in the genomes outside of the centromeres in shorter arrays. The repeats are more frequent on arms of the chromosome with the same repeat type forming the main centromeric array. However, the AlyCEN179 is more frequent even on non-AlyCEN179-type chromosomes throughout the genus (Supplementary Fig. S13a, Supplementary Table 1, p-value=0).

The PCA of five-mers from AlyCEN179 sequences across all species (Supplementary Fig. S8) shows that clustering is chromosome-specific rather than species-specific, similar to the patterns observed within *A. thaliana*, where centromeric haplotypes are more commonly shared between the syntenic centromeres from different accessions than between all centromeres within the same accession^17^. Sharing of 60bp-long k-mers between all the centromeres also shows high similarity between syntenic chromosomes of closely related species and lower sharing between different chromosomes of the same sample (Supplementary Data 2). Full repeat sequences are also shared between syntenic chromosomes of closely related species more so than between different chromosomes within the same species (Supplementary Fig. S5, S7). However, repeats from some AlyCEN179-type centromeres (e.g., *A. halleri* chromosome 2 and *A. lyrata* chromosomes 4 and 8) are not clustering with syntenic AlyCEN179 centromeres from *A. arenosa* and *A. croatica* (Supplementary Fig. S8), suggesting independent subsequent turnovers in centromeres of these species by the same AlyCEN179 repeat type.

### Origin of repeats

Repeat arrays forming a core centromeric region are also present outside of centromeres (Supplementary Table 1, Supplementary Fig. S13), mostly occurring in pericentromeric regions. Repeats located closer to a centromere tend to have the most similar sequence in that centromere compared to centromeres in other chromosomes. Repeats located far from the centromeres can have similar repeats in any nonsyntenic centromere (Supplementary Fig. S14). This suggests that repeats move in and out of the primary centromere-forming array.

Three (AlyCEN179, AlyCEN168, AlyCEN176) out of four major centromeric repeat types (excluding AthCEN178, which is present only in *A. thaliana*) are also present outside of the main centromeric arrays and sometimes even share the outside locations between *Arabidopsis* species (Supplementary Fig. S13). This suggests their non-centromeric origin. For example, *A. arenosa* centromeric repeats remain the ancestral AlyCEN179 type, while AlyCEN176 (orange, Fig. 1, Supplementary Fig. S13c) present outside of the *A. arenosa* centromeric array, in the right pericentromeric region on chromosome 6. AlyCEN176 took over the 1st, 2nd and 6th centromeres in *A. halleri* and the 6th centromere in *A. lyrata*. Apart from the pericentromeric region of chromosome 6 in *A. arenosa*, AlyCEN176 is also present in the same location in *A. cebennensis*, *A. pedemontana, A. lyrata* and *A. halleri*. This suggests that AlyCEN176 was present in the ancestral pericentromeres and only succeeded to take over in *A. halleri* and *A. lyrata*.

New repeats are continuously recruited to centromeres, as we detect several unrelated tandem repeats within centromeric arrays with no sequence similarity to the four major *Arabidopsis* centromeric repeat types (AlyCEN179, AlyCEN168, AlyCEN176, AthCEN178). We call such repeats within the centromeric arrays non-canonical (Supplementary Fig. S15):

(1) A repeat of length about 1.5 kb (Supplementary Fig. S15, Aa1500, red) forms several arrays with total length over 760 kb on the centromere of chromosome 2 of *A. arenosa.* Same repeat forms arrays outside of centromeres in *A. cebennensis*, *A. pedemontana* and some accessions of North American *A. lyrata*. The repeat sequence of Aa1500-type resembles a part of SAP2 (aspartyl protease) gene (a blast bit score=700, e-value=0), which is expressed in *A. lyrata*^25^. The blast hits span across the coding region of the gene (Supplementary Fig. S16a). This gene has a Helitron transposon located near it (Supplementary Fig. S16a), which could explain multiplication of this region as Helitrons can capture gene fragments during transposition^41^.
(2) Another repeat of length 365 bp (Supplementary Fig. S15, Ah365, pink) is present on the centromere of chromosome 2 and 6 in *A. halleri* and chromosomes 2 and 8 of *A. halleri* subgenome of *A. kamchatica* as large arrays from 550 to 2700 monomers long. The sequence of this repeat resembles a part of LTR retrotransposon (annotated by EDTA as ATHILA8A) spanning less than a half of the annotated retrotransposon model (Supplementary Fig. S16b).
(3) Most of *A. lyrata* accessions have arrays of a 696 bp-long repeat (Supplementary Fig. S15, Al696, green) present close to the centromere on the chromosome 2 and on pericentromeres of other chromosomes in some accessions. Two copies of the same repeat are present in the centromere on chromosome 2 in *A. croatica*.
(4) *A. cebennensis* and *A. pedemontana* have arrays of 335 bp-long repeat (Supplementary Fig. S15, Al335, blue) located at the right end of the centromeric array of chromosome 2 and in the centromeric array of chromosome 8, which is otherwise often present in pericentromeres of chromosome 2 in *A. lyrata*. Unlike Ah365 and Aa1500, the other two repeats Al335 and Al696 are present in the genomes almost exclusively in a form of arrays and do not have high identity with known TEs or genes. Therefore, Al335 and Al696 repeats are probably older than Ah365 and Aa1500 and already lost their original identity.

Thus, the new centromeric repeats likely originate from scrambled sequences of transposons, supporting the KARMA^17^ model, where uneven recombination leads to satellite homogenisation and centromeric transposon deterioration which in turn can lead to formation of new centromeric repeats. However, we observe that new centromeric repeats first form arrays in the pericentromeres, before invading the centromeres. It is possible that a single monomer or a short repeat array originating within a centromere will have less chance to spread and will be quickly lost compared to a longer array. We simulated the emergence and the spread of a new centromeric array with a different number of monomers allowing for gene conversion^21^. This simulation takes an array of 15,000 identical repeats with an insertion of 1, 20 or 100 identical repeats of a different type and allows the array to evolve for a million generations in 100 independent simulations. The changes at every generation include non-allelic gene conversions, point mutations and indels in frequencies observed in natural centromeres of *A. thaliana*^21^. Longer arrays had a higher chance of being maintained (Fig. 3, Supplementary Table 2). A zone of lower identity is formed around the inserted repeat array on both sides of array borders in the simulations due to conversion events between different types leading to degraded repetitive structure (Fig. 3, lower panels). However, in natural centromeres, repeats of two distinct types can be located next to each other and form complex high-order-repeats without losing the original identity of individual repeats (Supplementary Fig. S17). This suggests that a model based on gene conversion does not completely explain the evolution of centromeres within the species with different types of repeats.

**Figure 3.**
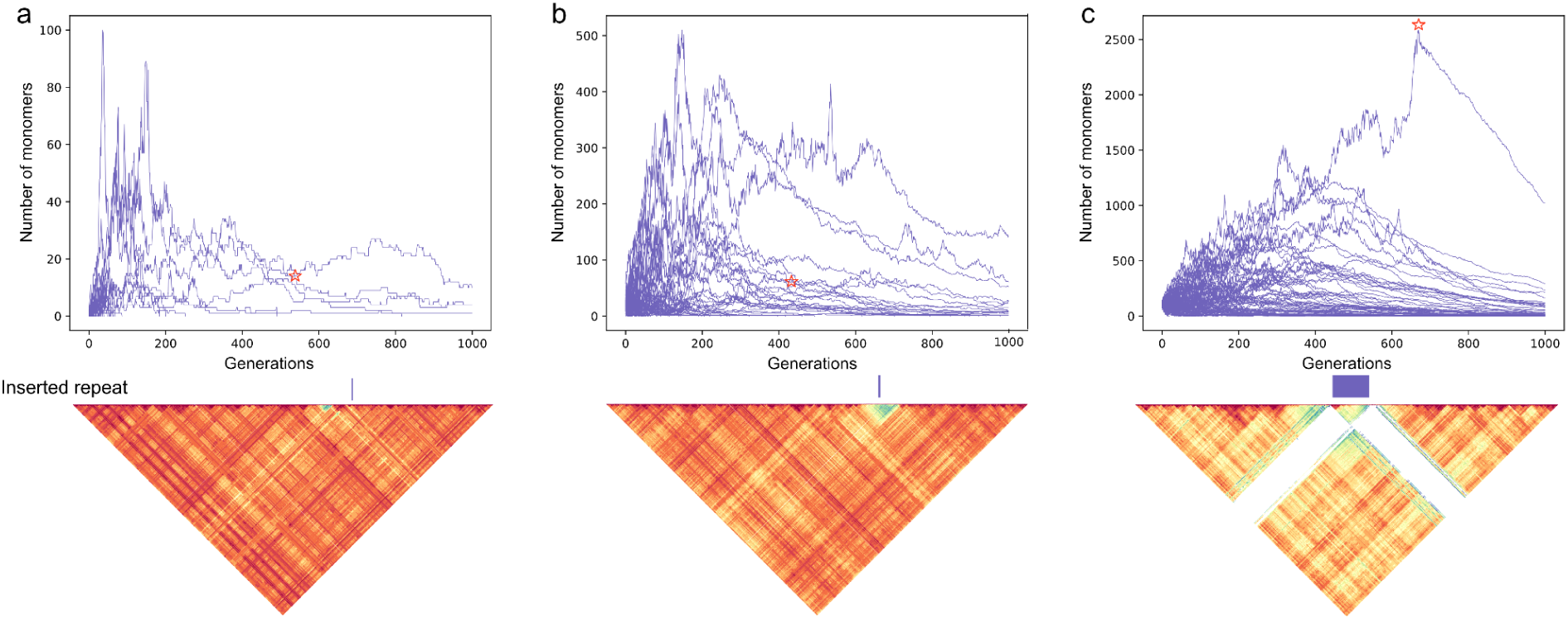
Simulation of monomer frequencies in centromeres with two repeat types. The model simulates point mutations, non-allelic gene conversion events and indels in frequencies observed in natural *A. thaliana* centromeres^21^. The longer new centromeric arrays have higher chances to take over the old centromeres. Simulations for one million generations with 15000 repeats of type AthCEN178 and (a) 1, (b) 20, (c) 100 AlyCEN179 repeats. Stained glass plots underneath show examples of simulated centromeres (marked with stars on the lineplots).

### Functional turnover of a centromere

To understand when the new repeats acquire their functionality and when the old repeats lose it, we performed ChIP-sequencing on NT1 *A. lyrata* accession and Kronotskiy *A. kamchatica* accession with an antibody designed for the protein encoded by the expressed copy of the *CENH3* gene of *A. lyrata* (see Methods, Supplementary Fig. S18, S19). The two accessions provide two different examples of a spread. The centromere on chromosome 6 of *A. lyrata* represents the classic scenario^18^, when a new repeat (AlyCEN168) array spreads inside a centromere and pushes the old (AlyCEN176) repeats towards the ends of the centromere in a form of previously described layered expansion^20,42^. We infer AlyCEN179 repeat type to be ancestral for all the centromeres, which was later replaced by AlyCEN176 in all the species, but *A. thaliana* and *A. arenosa*. This first turnover of the centromere on chromosome 6 into AlyCEN176 is evident from deep sharing of the AlyCEN176 repeats across *A. lyrata*, *A. halleri*, *A. cebennensis* and *A. pedemontana*, conserving the dimeric structure of this repeat which forms two distinct PCA clusters (Supplementary Fig. S10). Subsequently, the centromere arrays on chromosome 6 of *A. cebennensis* and *A. pedemontana* turned-over to AlyCEN168, pushing AlyCEN176 at the left edge of the new array (Fig. 1). The turnover of the same centromere on chromosome 6 in *A. lyrata* and the *A. lyrata* subgenome of *A. kamchatica* by AlyCEN168 is likely independent from *A. cebennensis* and *A. pedemontana*, as the k-mer composition on the AlyCEN168 repeats do not overlap (Supplementary Fig. S9, Supplementary Data 2). Therefore, the centromere on chromosome 6 of *A. lyrata* and the *A. lyrata* subgenome of *A. kamchatica* contains a new repeat array of AlyCEN168 in the middle and AlyCEN176 at the edges (Fig. 4 a,c). The CENH3 protein only binds to the new array (Fig. 4 a,c).

**Figure 4.**
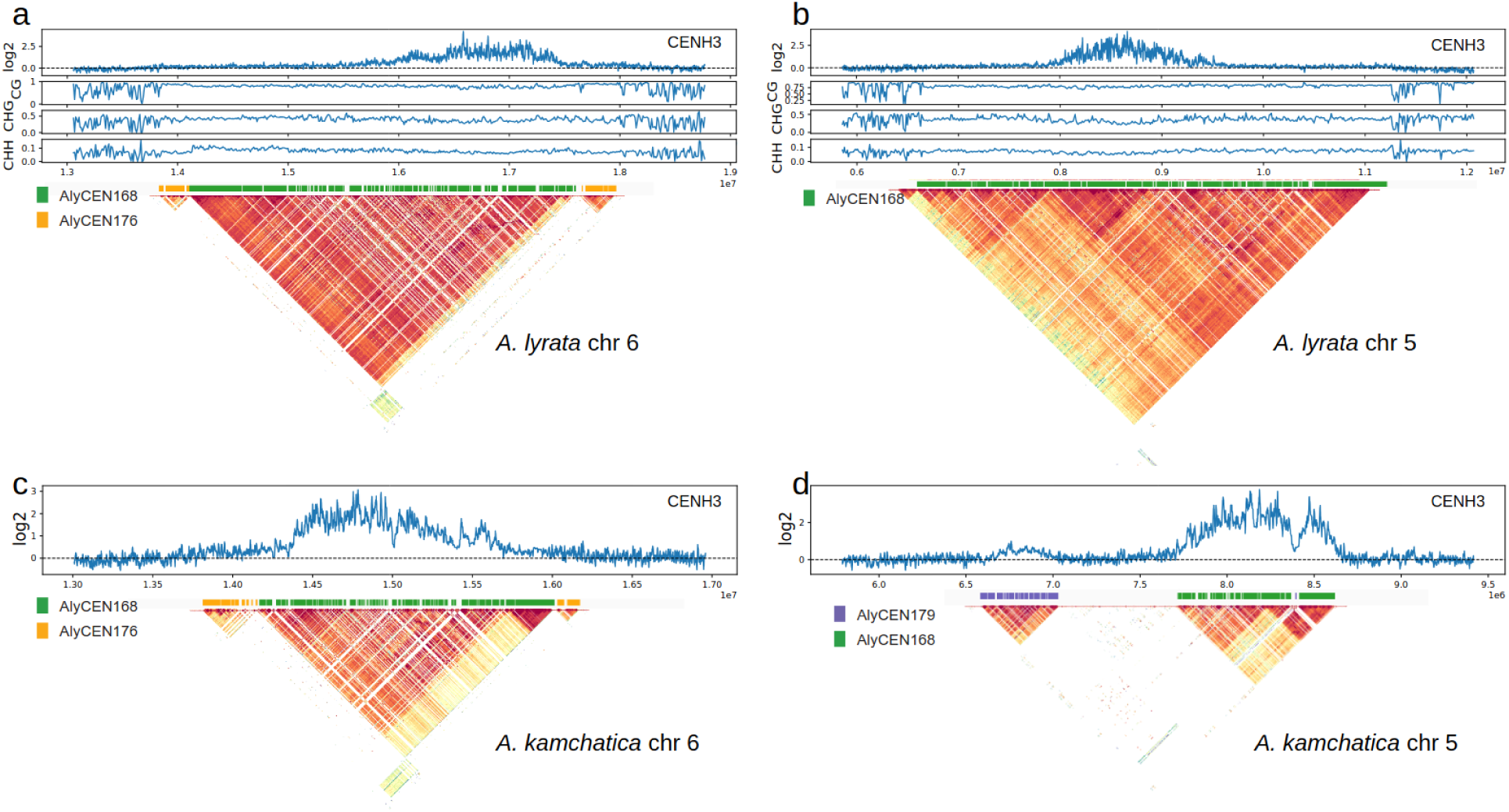
(a,b) Similarity heatmap for *A. lyrata* NT1 centromeres on chromosomes 5 and 6, DNA methylation and CENH3 binding, (c,d) similarity heatmap for *A. lyrata* subgenome of *A. kamchatica* centromeres on chromosomes 5 and 6 and CENH3 Chip-seq.

The centromere on chromosome 5 on the A. lyrata subgenome of Kronotskiy *A. kamchatica* represents a different scenario, where a new repeat array (AlyCEN168) occurs next to an old array (AlyCEN179). The older origin of the AlyCEN179 array is shown by sharing of frequent kmers with chr5 of *A. cebennensis* and *A. pedemontana* (Supplementary Fig. S20). Again, only the new array attracts CENH3, suggesting completed functional turnover (Fig. 4d). In both cases the total length of a new array binding CENH3 is longer compared to the old array.

The centromeres on all chromosomes attract CENH3, regardless of the major repeat-type. The methylation levels, especially in the CHG context, along the centromere are showing a slight decrease at the locations where CENH3 shows stronger binding (Fig. 4a,b), however, the decrease is not as pronounced compared to the observed levels in *A. thaliana*^13,43,44^ (Supplementary Fig. S21). CENH3 can bind throughout the entire centromere (chr2 of NT1 *A. lyrata*), only towards one end (chr5, chr7 of NT1 *A. lyrata*) where the similarity of arrays seems higher, or without a particular preference (chr7) (Fig. 4, Supplementary Fig. S22-S24). This raises a question if there is a concerted evolution of CENH3 proteins associated with centromeric repeat turnovers in *Arabidopsis.* Most *Arabidopsis* species have two copies of *CENH3* gene, apart from *A. thaliana* which shares the ancestral single copy with the outgroup species *Capsella* (Supplementary Fig. S18). The amino acid sequences of CENH3 vary between the species (Supplementary Fig. S25), however without a signature of positive selection, as they do not show the excess of nonsynonymous divergence on the entire gene or on the N-tail only (Supplementary Table 3).

### Spread of centromeric repeats between centromeres on different chromosomes

A complete turnover of centromeres in all the chromosomes to the same repeat type requires either multiple independent successful invasions by the same repeat of every centromere, which is unlikely, or an exchange of repeats between the centromeres. A recently formed allopolyploid species *A. suecica* provides the cleanest natural experiment for such exchanges, as it combines two subgenomes with complete sets of centromeres of different repeat arrays: *A. thaliana* (AthCEN178) and *A. arenosa* (AlyCEN179). We found multiple arrays of the *A. thaliana*-specific AthCEN178 repeat on the *A. arenosa* subgenome of *A. suecica*. One of the repeats was located next to an LTR retrotransposon (Supplementary Fig. S26), suggesting that the exchange likely occurred due to transposable element activity. We checked an additional 10 draft *A. arenosa* assemblies^45^ with BLAST for evidence of *A. thaliana* repeats segregating in *A. arenosa* centromeres, but did not find such evidence, strongly supporting that centromere repeat exchange occurred only in *A. suecica* following hybridization of *A. thaliana* and *A. arenosa*. Similarly, we found a small array of 10 AlyCEN179 monomers in the centromere of chr2 of the *A. thaliana* subgenome of *A. suecica* (Supplementary Fig. S26), but no such repeats in 66 representative *A. thaliana* assemblies^17^.

*A. lyrata* species with three types of repeats forming centromeres provides another opportunity to detect exchanges of repeat arrays between the centromeres. For example, an array of seven AlyCEN176 monomers on chromosome 3 of the Siberian *A. lyrata* NT12 accession is probably a result of such an exchange, as it is absent in other accessions (Supplementary Fig. S27b). Similarly, 32 AlyCEN168 monomers on chromosome 3 are private to the North American *A. lyrata* MN47 (Supplementary Fig. S27a). In both cases, the exchanged repeats are located close to an LTR retrotransposon.

### Transposable element content of satellite-based *Arabidopsis* centromeres

Repeat-based centromeres are invaded by centrophilic transposons across eukaryotes^46^. We therefore characterised the non-satellite portion of the centromeric arrays in the *Arabidopsis* genus for its TE content. Overall, on average 23.6% of the centromere space is not occupied by satellites across the genus, of which the majority (20.7%) is annotated as LTR retrotransposons, with the remaining allocated to other types of TEs or uncharacterised DNA (Fig. 5). We observed varying degrees of TE invasion based on the host species and centromeric repeat type, consistent with specific host-TE evolutionary dynamics but also different affinities of TEs for invading the four major repeat types. For example, it was previously shown that *A. thaliana* centromeres are depleted of TE content compared to *A. lyrata* centromeres^17^, which remains true with our increased sample size of *A. lyrata* and also in comparison to all other *Arabidopsis* species (Fig. 5). This is also consistent with the overall low genome-wide TE content in *A. thaliana* compared to other species resulting in smaller genome size overall^47^. In addition, *A. arenosa* and *A. croatica* centromeres have comparatively low TE occupancy, which is consistent with the lower TE abundance in their chromosome arms (Supplementary Fig. S28b). Finally, both *A. lyrata* and *A. halleri* show in general higher levels of TE invasion in the centromeres of all repeat types, which follows their high TE load across the genome (Supplementary Fig. S28b). We note, however, the substantial variation in *A. lyrata*, where individuals from Central Europe and Scandinavia have lower levels of TE invasion compared to Siberia and North America (Fig. 5, Supplementary Fig. S29). The types of TEs enriched in the centromeres also differ between the species (Supplementary Fig. S28a).

**Figure 5.**
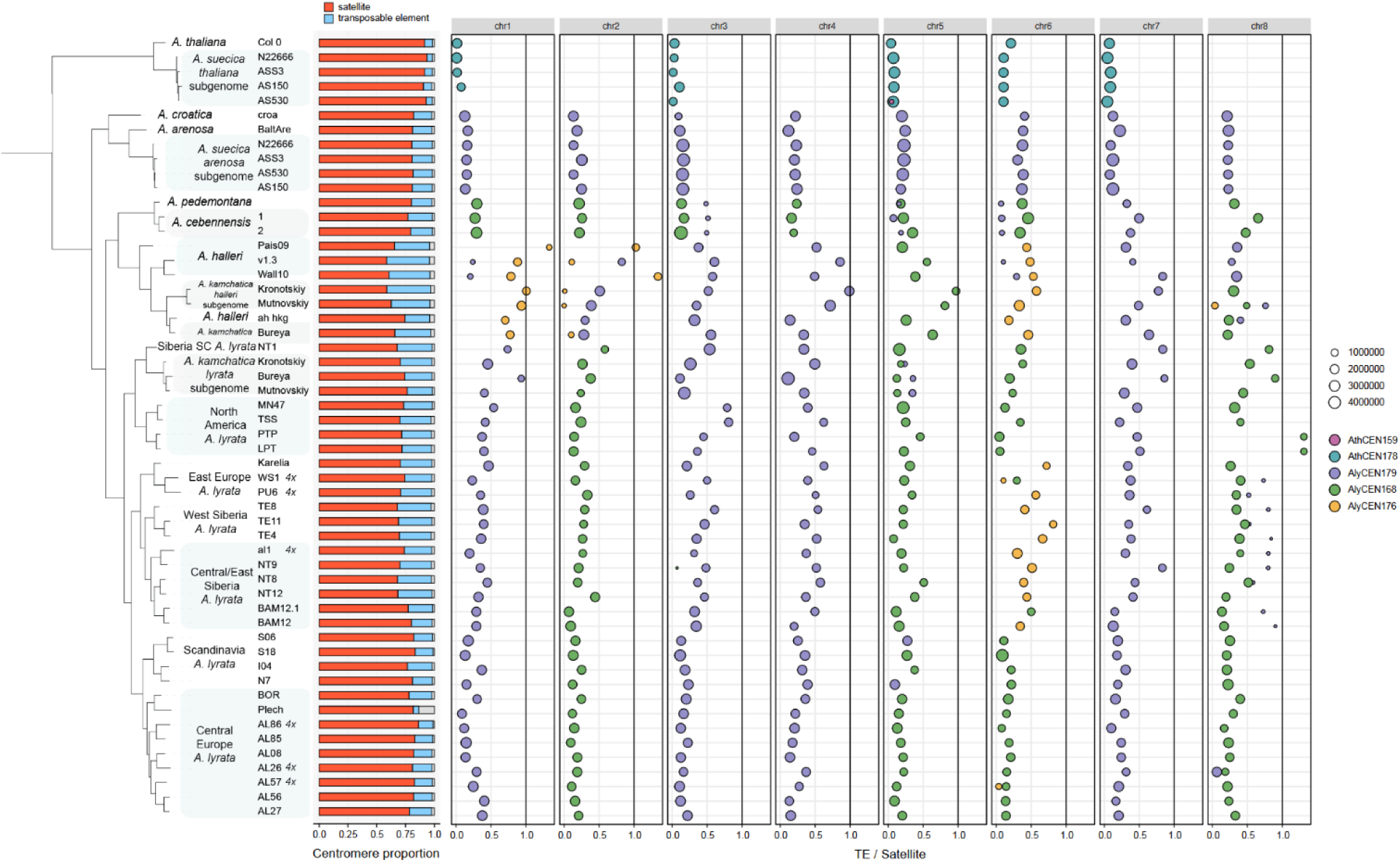
The ratio between TE and satellite DNA within every centromere of every chromosome, split and coloured by the main repeat type of the centromeric array. The size of the bubble corresponds to the size of the array.

### Evolutionary forces acting on centromeres

We notice that both major turnovers of centromeric repeat arrays from ancestral (AlyCEN179) to derived states in *A. thaliana* (AthCEN178) and in *A. cebennensis* and *A. pedemontana* (AlyCEN168) occurred in the species with evolutionary history of strong bottlenecks. While *A. thaliana* currently spread over the Northern hemisphere, earlier it experienced a strong demographic decline due to the karyotypic change from eight to five chromosomes^47^ and mating system shift from obligate outcrossing to self-compatibility^33,48^. The populations of *A. cebennensis* and *A. pedemontana* are currently geographically very restricted and depleted of genetic diversity^24,31,32^. At the same time, in *A. arenosa*, which has the largest effective population size among *Arabidopsis* species^24,25^, all centromeres remain of the ancestral AlyCEN179 repeat type. It is possible that fixation of one centromeric type can happen during a bottleneck and, therefore, turnovers may not necessarily require selection.

To test this hypothesis further, we investigate segregating centromeric alleles on chromosome 6 within *A. lyrata* species (Fig. 1). We genotyped two different alleles segregating on centromere 6, AlyCEN176 (orange on Fig. 1) and AlyCEN168 (green on Fig. 1), in the larger dataset of 600 *A. lyrata* samples^24,28,49–55^ across the species range by overall proportions of repeat types (see Methods) in the whole-genome short-read sequences. The older centromeric repeat (AlyCEN176, Supplementary Fig. S10) is maintained at high frequencies in Siberia: it is prevalent over a wide territory from Karelia to Chukotka, while North America, Scandinavia and Central Europe are fixed for the newer array (AlyCEN168) (Fig. 6a). Only 0.3% of all SNPs segregating in *A. lyrata* and only 0.08% of 4-fold degenerate sites follow a similar geographical distribution pattern of allele frequencies, suggesting that this scenario is possible but not the most likely under neutral expectations of random fixation (see Methods).

**Figure 6.**
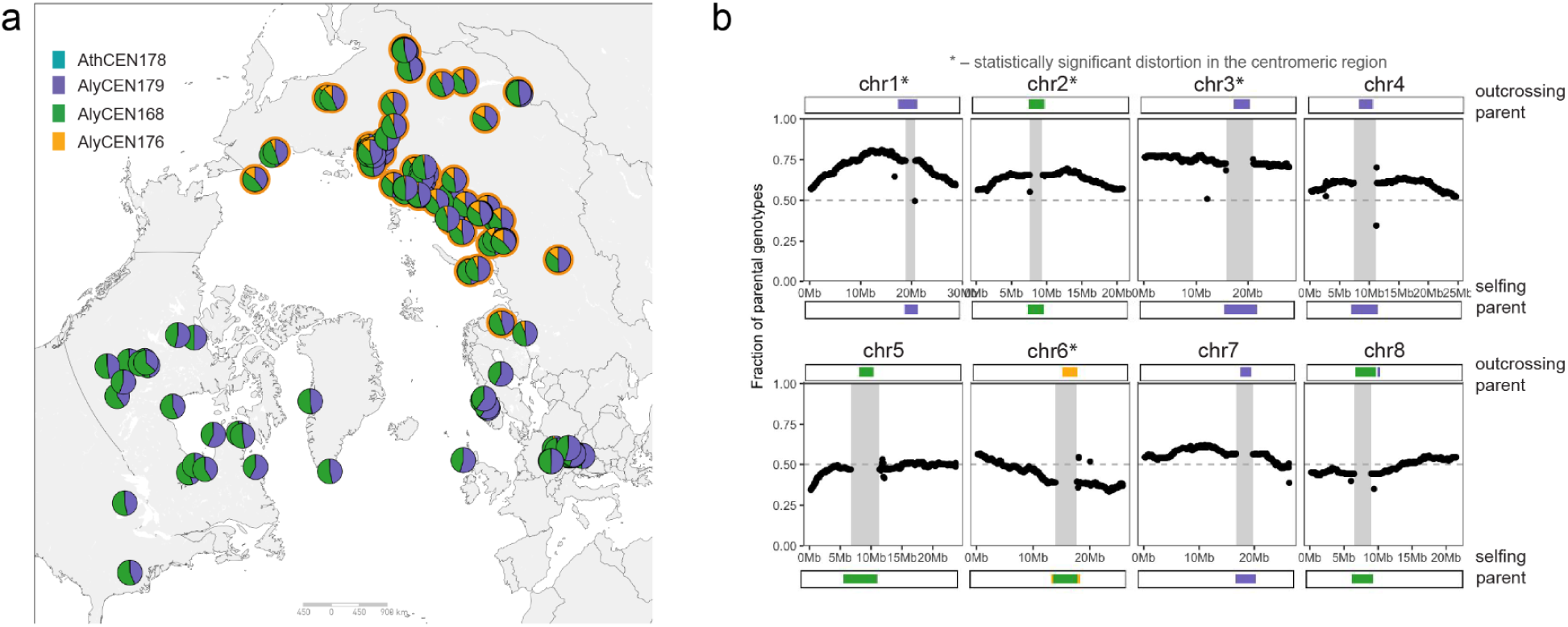
(a) Geographic distribution of total centromeric repeat type composition in *A. lyrata* calculated from whole genome short-read genotyping. The differences in total centromeric repeat compositions are mainly driven by centromeres on chromosome 6 segregating AlyCEN176 (orange, frequent in Siberia) and AlyCEN178 (green, frequent in Europe and North America) of *A. lyrata*. Higher proportion of AlyCEN179 in Scandinavian *A. lyrata* accessions is likely driven by centromere on chromosome 5. (b) Segregation distortion towards AlyCEN168 type centromere on chr6 in cross of Siberian selfing and Siberian outcrossing lineage. The grey bar in the center is representing the masked centromeric array region of the reference genome.

While the demographic history can play a role in the centromeric turnovers, fixing a new centromeric allele, it is also possible that a new allele gets an advantage through, for example, meiotic drive. In order to check whether the two centromeric alleles on chromosome 6 in *A. lyrata* are segregating normally or their inheritance is distorted, we genotyped 111 F_2_ offspring self-fertilized individuals from a F_1_ cross between maternal TE10 (AlyCEN176) and paternal NT1 (AlyCEN168) *A. lyrata* accessions. F_1_ plants are self-compatible, as NT1 has a loss-of-function *S*-allele (responsible for self-incompatibility in outcrossing *Arabidopsis*) that is dominant over both functional *S*-alleles in TE10^49^. We observed a significant segregation distortion on chromosome 6 covering a wide region including the centromere (Fig. 6b), however, we also observed distortion on another three chromosomes, spanning over centromeric regions. While segregation distortion is common throughout the genome in the outcrossing *A. lyrata* and *A. halleri*^56–60^, the distortion we observe on the centromere 6 is towards the newer (AlyCEN168) centromeric array and is also the only distortion towards the selfing parent. To test if such segregation distortion on centromere 6 is due to pre-pollination gametic selection, we performed bulk pollen sequencing^57,58^ of F_1_ plants from two crosses: (1) between Siberian selfing NT1 (AlyCEN168) and Siberian outcrossing lineage NT8 (AlyCEN176) and (2) between Siberian outcrossing TE11 (AlyCEN176) and Central European outcrossing lineage AL62 (AlyCEN168) (two plants). Pollen from these F_1_ plants did not show segregation distortion (Supplementary Fig. S30), indicating that distortion in the F_2_ plants is likely caused by other mechanisms than pre-pollination male gametic selection such as, for example, female meiotic drive (Fig. 6).

An active drive can leave a similar signature on allele frequencies as a selective sweep^61^. To test for sweeps we run SweeD^62^ on a subset of European, Scandinavian and Siberian *A. lyrata*. However, no clear peaks associated with new centromeric arrays were found on the composite likelihood ratio close to the centromeres (Supplementary Fig. S31). It is possible that we find no linked sweeps to centromeres in *A. lyrata* because of low linkage disequilibrium in the outcrossing plants and because crossovers can occur close to centromeres in Arabidopsis^63,64^. To check how far the linkage extends at the adjusted regions to the two centromeric alleles of different arrays AlyCEN176 and AlyCEN168 in *A. lyrata*, we focus on the two genetically closest accessions as they are sampled in the same location, but different in centromeric alleles on chromosome 6 (BAM12 - AlyCEN176, BAM12.1 - AlyCEN168, Fig. 1a). While centromeric regions are clearly distinct, the pericentromers cluster together up to 50 kb close to the centromere (Supplementary Fig. S32). This distance is shorter compared to the smallest reported non-recombining region in *A. thaliana* (90 kb)^43^. Therefore, it is possible that we could not identify signatures of selective sweeps which should accompany a meiotic centromeric drive because the crossovers occur very close to the centromere and unlink the adjusted regions. Overall, we suggest that both mechanisms can play a role: the neutral fixation of new centromeric alleles due to drift and the active meiotic drive on one of the centromeric alleles, as we could not reject either of these hypotheses.

## Discussion

This study reveals the complex nature of centromeric repeat turnovers in the *Arabidopsis* genus. Using complete genomes of every species within the *Arabidopsis* genus, we track the centromeric repeats from their origin, their transposition into an existing centromeric array, their spread within a centromeric array and their spread from one chromosome’s centromere to another, their functional take over of an old centromeric array, to a complete turnover of all centromeres by a new centromeric type in a species.

All centromeres of the *Arabidopsis* genus are repeat-based with four repeat types (AthCEN178, AlyCEN179, AlyCEN168, AlyCEN176, see Fig. 1, Supplementary Fig. S3) forming main centromeric arrays. We infer that AlyCEN179 is an ancestral repeat type, as it is the most prevalent across the genus and present in 8 out of 9 *Arabidopsis* species. In *A. thaliana*, where the AlyCEN179 is absent, we find monomers which are intermediate by sequence homology between AlyCEN179 and the main centromeric repeat type in *A. thaliana* AthCEN178 (Fig. 2). The intermediate repeats are located on the periphery of *A. thaliana* centromeric arrays, suggesting their older state compared to AthCEN178. All centromeres in *A. thaliana* species turned over from the ancestral AlyCEN179 to AthCEN178, with some minor presence of AthCEN159, also in the peripheral zones of the centromeric arrays. This centromeric repeat type is probably also relatively ancient as it is close to *C. orientalis* centromeric repeat CoCEN168 by sequence homology and is also shared with centromeres of *A. cebennensis* and *A. pedemontana* and pericentromeres of *A. halleri*.

An almost complete turnover (all centromeres but chromosome 7) into a AlyCEN168 centromeric repeat type occurred in a shared ancestor of *A. cebennensis* and *A. pedemontana*. Turnovers of some centromeric arrays into the same AlyCEN168 repeat type occurred in *A. lyrata* and *A. halleri*, were likely independent from the turnover in *A. cebennensis* and *A. pedemontana,* which is evident from their different k-mer composition of the repeats (Supplementary Fig. S9, Supplementary Data 2). Some centromeres of *A. lyrata* and *A. halleri* turned over to AlyCEN176. Repeat types AthCEN178, AlyCEN179, AlyCEN168 and AlyCEN176 share high identity values^33,34^, which points at their common origin.

We found several new repeat types in the centromeres, not related to any previously described centromeric satellites of *Arabidopsis* (Al335, Al696, Aa1500, Ah365, see Supplementary Fig. S15). Interestingly, these repeats are present outside of the centromeres too and share a high identity with non-centromeric TEs and even a coding gene (Supplementary Fig. S16). This supports the KARMA model^17^ with the difference that the source of new centromeric satellites is not centromeric but rather pericentromeric TEs that already formed short arrays outside of the centromeres. The origin of repeat arrays from TEs and their higher affinity to CENH3 compared to TEs themselves have been recently described in *R. canina*^65^. Our simulations of an insertion of a new repeat type into the centromere suggests that a single monomer can not outcompete the existing array, while a longer newly inserted array gives substantial chances for a new repeat to proliferate. Overall, in *Arabidopsis* data we observe that different repeats can proliferate in centromeres for a long time without taking over a full centromeric array. The extreme version of ancestral repeat conservation in holocentric *Rhynchospora*^7^ across 20 species and more than 40 Myr, where short interspersed centromeric arrays are gained, lost and rearranged rather replaced, suggests that centromeric repeat turnover requires a permissive architecture. We suggest that turnovers in monocentric *Arabidopsis* occur in a single locus per chromosome, where and when one array can capture the whole kinetochore, as our CENH3 binding data shows (Fig. 4).

The spread of the centromeric repeats from one chromosome to another is likely TE-driven. We find evidence for this in allotetraploid *A. suecica*, a clean case of hybridisation between two species with different repeat types fixed in all chromosomes: *A. arenosa* (AlyCEN179) and *A. thaliana* (AlyCEN178). We tracked the appearance of *A. arenosa* type of repeats in *A. thaliana* subgenome centromeres and the other way around, and showed that the moved repeats are flanked by TEs (Supplementary Fig. S26).

We found cases of recent functional turnover of the new centromeric repeat type over the old one. In the centromere of chromosome 6 in *A. lyrata* the old repeat type (AlyCEN176) is pushed towards periphery by a new repeat type for this centromere (AlyCEN168), and only the new repeat attract CENH3. This scenario is consistent with previously described dynamics of centromeric repeats through layered expansions^20^. In the case of *A. kamchatica*, a centromere of chromosome 5 of the *A. lyrata* subgenome consists of two long arrays separated by a gap. Only the new array (AlyCEN168) attracts CENH3, while the old one (AlyCEN179) is rendered non-functional. In both cases the new arrays are longer compared to the old ones, which may indicate that the functional turnover depends on the relative abundance. When centromeres on different chromosomes formed by these types of repeats, all repeat-type centromeres attract CENH3, however, when two different repeat types are abundantly present in one centromere, one of the repeat types takes over the function of CENH3 binding. The *CENH3* duplication is shared between *Arabidopsis* species with the exception of *A. thaliana*, however, there is no relationship between CENH3 protein sequence and diversity of repeat types in the species (Supplementary Fig. S25).

The species which show turnovers of centromeric arrays into derived types, *A. thaliana*, *A. cebennensis* and *A. pedemontana*, also experienced severe demographic bottlenecks in their evolutionary history^33,47,48,66,67^, while the only species retaining all the centromeres in the ancestral repeat type, *A. arenosa*, has the largest effective population size among *Arabidopsis* species^24,25,68^. It is possible that a fixation of new centromeric arrays occurred in a process of random genetic drift. However, we also observe that two centromeric alleles on chromosome 6 in *A. lyrata* violate Mendelian distribution and segregate unevenly (Fig. 5b), favoring a newer centromeric array in the F_2_ generation. Together with the absence of segregation distortion in pollen of chromosome 6 centromere heterozygous F_1_, one possible scenario is that female centromeric drive^11^ takes place in *A. lyrata* and promotes centromeric array turnovers. However, we can not rule out that segregation distortion in F_2_ generation at the centromeric regions is due to non-meiosis related mechanisms acting via gametic or zygotic stages such as egg or pollen competition, female gametic selection, zygotic selection or allele incompatibilities affecting for instance seed germination^69–75^. We also observe segregation distortion in other chromosomes, which have the same centromeric arrays in both alleles, as distortion is common in the outcrossing *A. lyrata* and *A. halleri*^56–60^. Interestingly, in the hybrids between outcrossing and selfing *Mimulus* species, a centromere drive favored the outcrossing parent^76^, while we observe favoring of selfing parent on the chromosome with segregating centromeric alleles, and the rest of the chromosomes with segregation distortion favor the outcrossing parent instead. The geographical distribution of the centromeric alleles shows that the older repeat type allele (AlyCEN176, orange) remains frequent in Siberian populations of *A. lyrata*, while the new repeat type allele (AlyCEN168, green) takes over the European and North American *A. lyrata* (Fig. 5). Although rare (0.08% of the 4-fold degenerate sites), some neutral SNPs show similar geographical distribution, however, it is also possible that environmental conditions can favour one of the alleles, for example, if temperature impacts the efficiency of CENH3 binding to one of the centromeric repeat types^77^. Overall, we can not confidently reject either of the mechanisms acting on evolution of centromeric arrays, the turnovers can potentially be promoted by neutral fixation of one of the centromeric alleles due to a bottleneck or can be favored due to a centromeric drive in meiosis or due to selection acting on a linked trait.

Here be describe the turnovers between different centromeric satellite repeats within evolutionary history of one genus. On a bigger evolutionary scale, the turnovers occur between satellite repeats and transposable elements forming functional centromeric arrays^46^. Both types of centromeric architectures can even co-exist within one genome^65,78^. Satellite arrays emerge, maintained via homogenization by gene conversion or replaced with new satellite arrays, all while being invaded by centrophilic transposons. Inserted transposons disrupt the function of satellite repeat binding CENH3 until TEs themselves can reach a certain copy number and functionally replace the satellite arrays^46,65^. More focal studies within species and genera with contrasting centromeric contents, demography and mating systems are required in the future to understand the evolutionary forces behind one of the most rapidly evolving regions of eukaryote genomes.

## Methods

### Plant material and sequencing

*A. lyrata*, *A. arenosa*, *A. cebennensis* and *A. pedemontana*, *A. croatica* and *A. kamchatica* plants were grown in the greenhouse under 21 °C, 16 h day. Approximately 2 g of leaf material was collected for sequencing. High molecular weight DNA was extracted with a NucleoBond HMW DNA kit (Macherey Nagel). HiFi libraries were then prepared according to the manual “Procedure & Checklist — Preparing HiFi SMRTbell® Libraries using SMRTbell Express Template Prep Kit 2.0” with an initial DNA fragmentation by g-Tubes (Covaris) and final library size selection on BluePippin (Sage Science). Size distribution was again controlled by FEMTOpulse (Agilent).

The DNA was sequenced with PacBio Sequel II or PacBio Revio device (Supplementary Data 1). *A. halleri* plants Pais09 and Wall10 were grown and maintained in a greenhouse through vegetative propagation as described in ^79^ and crossed (Pais_o9 as mother) to obtain an F1 hybrid individual. High-molecular-weight DNA (NucleoBond HMW kit) of the F1 individual was depleted of fragments < 25 kb (Short Read Eliminator) and sequenced on a PacBio Sequel II. Additionally, parental DNA was extracted from snap-frozen young leaf tissue (NucleoMag Plant kit), and PCR-free TruSeq libraries were generated according to the manufacturer’s instructions (Illumina) and sequenced on an Illumina NovaSeq 6000 (150 bp paired-end, ≥ 100x coverage per parent; Pais_09, Wall_10; Novogene, Munich DE).

### ONT sequencing of NT1 *A. lyrata* accession

Three week old seedlings were grown on half-strength Murashige and Skoog (½ MS) medium supplemented with 1% (w/v) sucrose at 21°C, 16 h day. Approximately 4 grams of fresh tissue was harvested, flash-frozen in liquid nitrogen, and ground into a powder using a mortar and pestle. HMW genomic DNA was extracted using the NucleoBond HMW DNA kit (Macherey-Nagel), following the manufacturer’s protocol optimized for plant material. Barcoded adapters were ligated to the DNA using the Oxford Nanopore Native Barcoding Kit, compatible with the v14 sequencing chemistry, and R10.4 flow cells. DNA libraries were sequenced on a PromethION platform using R10.4 flow cells. Base-calling was performed using Dorado (version 0.9.5). Genomes were assembled using Hifiasm (version 0.25.0) with the option ‘--ont’.

### Genome assembly

The genomes of *A. lyrata*, *A. arenosa*, *A. cebennensis* and *A. pedemontana*, *A. croatica* were assembled with Mabs-hifiasm^80,81^ and scaffolded using RagTag^82^ against NT1 assembly of self-compatible *A. lyrata* published earlier^49^. Hifiasm 0.25.0-r726 with --dual-scaf option was used for *Capsella orientalis* assembly. *A. halleri* sample ah_hkg was assembled with hifiasm and scaffolded with European *A. halleri* genome as a reference. Scaffolds shorter than 100 kb and sequences with >99% coverage by organelle assemblies were removed.

*A. suecica* genomes were assembled with hifiasm (version 0.25) with a genome size set to 320Mb. For accessions ASS3 and AS530, published Hi-C paired-end short reads were also included in the hifiasm assembly using the options -h1 and -h2^83^. Contigs were scaffolded to a published *A. suecica* reference^83^ using the software RagTag (version 2.1.0). Reads were mapped using Winnowmap (version 2.03) and inspected manually for inconsistencies with assemblies, such as significantly increased or decreased coverage.

Parental short reads of Pais09 and Wall10 *A. halleri* samples were trimmed with Cutadapt v3.7 (TruSeq adapters, Q30 ends, poly-G removal, pairs ≥ 120 bp retained)^84^. F1 HiFi reads were cleaned with HiFiAdapterFilt v2.0.0^85^ using defaults. Genome size and heterozygosity were estimated in Meryl v1.4.1^86^ 31-mer histograms fitted with GenomeScope2 using diploid model^87^. Parent-specific 31-mer databases were built with yak v0.1-r66^88^, and haplotype-resolved contig level assemblies were generated with hifiasm v0.19.9-r616^89^ in trio mode^90^; of two conFigurations (default trio; --trio-dual), the -trio-dual assembly was carried forward. Haplotype 1 is the paternal (Wall_10), haplotype 2 the maternal (Pais_09) genome.

Genomic DNA of the three *A. kamchatica* accessions was sequenced using a combination of PacBio HiFi and and Hi-C. For each accession, *de-novo* assembly was performed with hifiasm (version v0.19.3)^89^, utilizing both the HiFi long reads and the Hi-C data. The parameter ‘nhap=2’ was specified to guide the software to phase the assembly into two distinct groups, which correspond to the two subgenomes. The contigs were scaffolded into chromosome-level sequences with HapHiC (version 1.0.2)^91^. The contact maps were visualized and corrected in Juicebox^92^ while the curated assembly was regenerated with the HapHiC ‘juicer post’ utility. Homeologous chromosome pairs were identified with GENESPACE^93^ by sequence synteny to the diploid progenitors, *A. lyrata* NT1 and Japanese *A. halleri* and assigned to separate subgenomes.

Then scaffolding was manually curated based on evidence from GENESPACE^93^ for gene synteny, TRASH^40^ for centromere locations and tidk toolkit^94^ for telomere locations (short contigs that were scaffolded by ragtag but occurred after the telomere repeat locations, were trimmed from the ends of scaffolds). Long reads were mapped back to the assemblies using Winnowmap^95^ to find regions with zero coverage. OrthoFinder^96^ tree from the GENESPACE analysis was also used to show the phylogenetic relation between the genomes on the left panel of Fig. 1.

### Centromeric region annotation

Centromeric repeat arrays coordinates were annotated using TRASH^40^ using *A. lyrata* NT1 lineage consensus repeat as a template. For *A. thaliana* genomes, Col-0 consensus repeats were used as a template instead. In addition, to confirm that the repeats have similarity with known centromeric repeats, we ran EDTA v2.0.1 with the *A. thaliana* TAIR10 TE database. Centromeric repeats of all known types are annotated as “AT1TE49780_ATREP18” by EDTA. Centromeric boundaries were identified from the first to the last centromeric repeat with gaps no longer than 100 kb between the repeats. Repeat regions for this annotation were taken by the widest coordinate between TRASH and EDTA annotations.

### Repeat library construction

All TRASH^40^ annotated repeats overlapping centromere coordinates were filtered by repeat length to remove the outliers that were much longer or much shorter than expected centromeric repeats. We kept all the repeats if they had high frequencies to allow for repeats with strong higher-order structure.

For the principal component analysis, the sequences of individual repeats were broken into k-mers of length 5, taking into account both forward and reverse strands. The frequencies of each k-mer were used for the PCA. A similar approach was used in other centromere-focused studies previously^97^. The first two PCs were used to cluster the samples by repeat type in each sample. If a sample had one repeat type prevalent (for example, in *A. arenosa*), we additionally projected k-mer matrix from this sample on the principal components of Siberian *A. lyrata* TE8 which had enough repeats of three repeat types. Repeat with annotated types were plotted in the coordinates of the corresponding genome or visualised using IGV^98^.

To find non-canonical repeats and canonical repeats outside the centromeres we used nucleotide BLAST^99^ of the consensus sequences.

To show overall repeat sharing between the centromeres we found all 60-mers present in the centromeres using jellyfish^100^, filtered out k-mers with less than 50 occurrences across the whole dataset and calculated weighted Jaccard index between the centromeres.

### ChIP-Seq

Anti-CENH3 rabbit antibody was produced and purified by LifeTien, USA. The peptide sequence used for immunisation was RTKHFATRTGSGNRTDA-C.

The suitability of the anti-CENH3 antibody for ChIP-seq was first assessed by immunostaining chromosome spreads prepared from flower buds (Supplementary Fig. S19). Samples were fixed in 3% formaldehyde, permeabilized, and blocked in PBS containing BSA, followed by incubation with the primary anti-CENH3 antibody overnight at 4℃. After washing, samples were incubated with a fluorophore-conjugated secondary antibody, counterstained with DAPI, and examined by fluorescence microscopy. The antibody was considered suitable for further analyses if it produced a clear centromeric signal with low background.

ChIP-seq was performed according to^101^, with minor modifications. Briefly, chromatin was isolated from 10 g of plant tissue and cross-linked with formaldehyde. The chromatin was then fragmented by sonication to an average size of approximately 300 bp and immunoprecipitated using the anti-CENH3 antibody, while a portion was retained as input control. Three to six technical replicates were used. Chromatin–antibody complexes were recovered using anti-rabbit Protein A beads. After reversal of cross-links, the enriched DNA was purified by phenol:chloroform, used for library preparation, and sequenced on an Illumina platform.

The ChIP-seq analysis workflow was adapted from^17^, with minor modifications. Briefly, the snakemake workflow includes read deduplication, trimming and genome alignment using Bowtie2^102^, with the modification of allowing up to 200 alignments per read (-k 200) to retain multimapping information. Aligned reads were subsequently filtered using SAMtools to retail properly paired reads with MAPQ ≥ 2 and allowing for no more than two mismatches. Multimapping reads were retained, to capture the signal in repetitive genomic regions. ChIP enrichment profiles were calculated as the log2(ChIP/input) signal, after CPM normalization, using deepTools bamCompare^103^.

### TE annotation and analysis

Transposons were annotated using EDTA v2.0.1^104^ and the library was further classified with TESorter^105^.

### Segregation Distortion

F_2_ seeds were obtained after self-pollination of the F_1_ plant from a cross between TE10.3-2 (mother) and NT1.3-1 (father). To collect leaf material we grew F2 plants in common conditions for one month (22 °C day, 18°C night, 14 day light, 80% humidity).

We used approximately 1 cm^2^ of leaf tissue for the library preparation of each individual F_2_. Genomic DNA was isolated with the “NucleoMag© Plant” kit from Macherey and Nagel (Düren, Germany) on the KingFisher 96Plex device (Thermo) with programs provided by Macherey and Nagel. Libraries were pooled and sequenced on the NovaSeq 6000 S4 flow cell Illumina system.^104^

Short reads were mapped to the NT1 reference^49^ using bwa mem v0.7.17^106^ and samtools v1.15.1^107^. A vcf file with F2 plants and parental NT1 and TE10 was made with GATK version 3.8-1-0 using HaplotypeCaller, CombineGVCFs and GenotypeGVCFs^108^. The filtering was done according to the GATK Best Practices recommendations^108,109^. We removed sites heterozygous in all samples with bcftools^107^. We calculated the fraction of parental genotypes along the genome in 200 kb windows overlapping by 50 kb and plotted the results with the R ggplot2 package^107,110^.

### Pollen sequencing

Roughly 50 flowers from three F_1_ plants (two plants from cross AL62xTE11 and one NT8xNT1 plant) were collected for the bulk pollen sequencing. DNA extraction and data analysis followed the protocol from ^57,58^. Briefly, flowers were collected into ethanol, pollen was spinned down and DNA extracted with the Macherey Nagel Nucleospin food kit. Reads, sequenced with NovaSeq 6000 were mapped, genotyped with GATK v4.6.0.0, phased with WhatsHap^111^ and filtered for sufficient coverage and phasing quality with the previously tested pipeline^57,58^. Additionally, only forward reads were filtered using the same pipeline, mapped and genotyped with bcftools^107^ to get unbiased coverage estimations. The coverages were plotted following the pipeline.

### Genotyping of total centromeric composition using short-read WGS data

Whole-genome sequencing short reads from previous studies^24,28,49–55^ were mapped to an artificial array made from several monomers for each repeat type. Reads were mapped using BWA v0.7.17^106^, removal of duplicates and sorting was performed using samtools v1.15.1^107^. Then we counted the number of reads mapped to each repeat type and calculated the proportion of each type from the overall number of mapped reads. Geographical map (Fig. 5a) of centromere repeat proportions was visualized using QGIS.

### Phylogenetic tree construction

For CENH3 gene tree construction, genomes were annotated using Helixer^112^. Then, CENH3 genes were found using protein BLAST of *A. thaliana* gene sequence. Assemblies from ^28,45^ were added to the analysis. Both for CENH3 and repeat trees the following methods were used. DNA regions were aligned using L-INS-i algorithm of MAFFT^113^, then manually curated in UGENE^114^ and the tree was constructed with IQTree with fast bootstrap = 1000^115,116^ for DNA as well as protein sequences. The translated protein sequence was extracted with UGENE based on *A. thaliana* exon-intron boundaries. The MK test on CENH3 sequence was performed with mkado^117^.

### Centromeric arrays simulations

Forward time simulations for 1 million generations were performed following the protocol and parameters from ^21^ using the same input data but with additional 1, 20 or 100 copies of AlyCEN179 added as a single array and simulated independently 100 times.

### Selection scans and the SNP analysis

For the selection scans with SweeD^62^ and SNP frequency analysis we used SNP data from arabidopsislyrata.org^25^. SweeD was run with grid parameter = 500 and *A. cebennensis* as an outgroup. Centromeric arrays on the reference genome were masked for the analysis.

## Data Availability

The raw sequencing data and assemblies are accessible on ENA under project number PRJEB61074. The full list of accessions can be found in the Supplementary Data 1. The code used in this study can be found at https://github.com/novikovalab/Arabidopsis_pancentromere.

## Acknowledgements

We thank the MPIPZ Genome Center for performing HMW DNA extractions, library preparations and PacBio HiFi sequencing, as well as Soile Alatalo and Jana Flury for DNA extraction. We thank Marc Stift and Roswitha Schmickl for providing plant material for American and European *A. lyrata* samples and advice during the TAC meetings, and Mohsen Falahati Anbaran for *A. lyrata* sample collection from Norway. Jon Ågren and Pawel Wasowicz are acknowledged for sharing information on the Swedish and Icelandic *A. lyrata* populations. PYN acknowledges the European Research Council (ERC, HOW2DOUBLE, 101041354), the joint Deutsche Forschungsgemeinschaft (DFG) and GACR program—project number 490698526, and the exchange program by DAAD fund 57601834 for promoting scientific exchange. A.Mac. acknowledges funding from the Birgitta Sintring Foundation (S2024-0025). P.Y.N. and A.D.S., thank the DFG-funded research consortium TRR341 for support and scientific discussion. EMBO long-term postdoctoral fellowship ALTF224-2022 and Broodbank Research Fellowship from the University of Cambridge to R.B. L.A.R. and A.Mar. acknowledges funding from the European Research Council (ERC, HoloRECOMB, 101114879) and the DFG (grant MA 9363/6-1). U.K. acknowledges funding through TRR 341/1-456082119 (Deutsche Forschungsgemeinschaft, A.P.) and ERC-AdG LEAP-EXTREME (788380). T.M.M. was supported by the Research Council of Finland grant no. 360310. The views and opinions expressed are however those of the author(s) only and do not necessarily reflect those of the European Union or the European Research Council Executive Agency. Neither the European Union nor the granting authority can be held responsible for them.

## Notes

### Competing Interest Statement

The authors have declared no competing interest.

