## Supplementary Figures 1-32 for "The Nature of Centromeric Repeat Turnovers in the genus *Arabidopsis*"

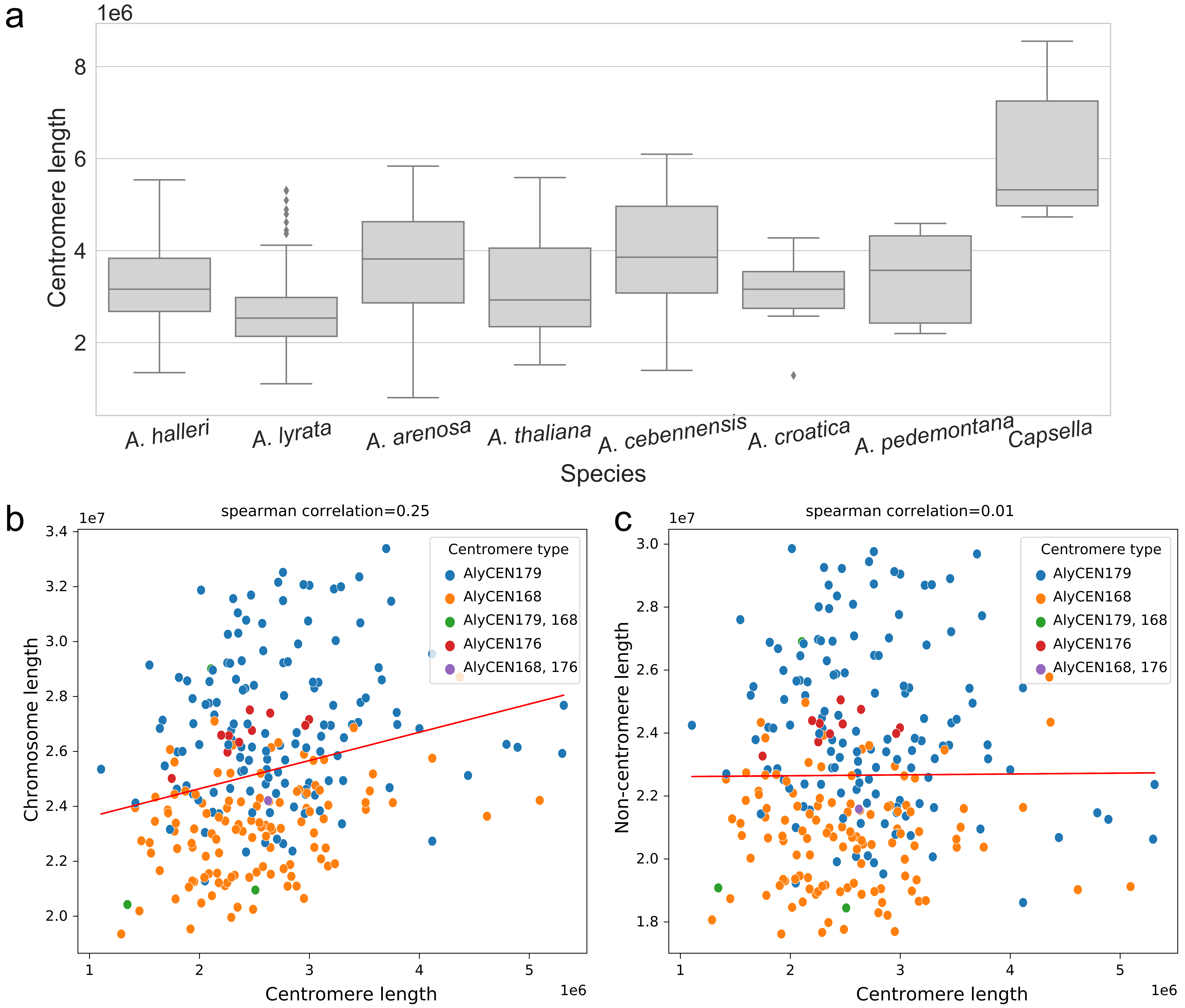


Figure S1. a) Centromere lengths (Mb) per species. *Capsella orientalis* has longer centromeres than any *Arabidopsis* species (p-val < 0.01). b,c) Correlation between *A. lyrata* centromeres and overall chromosome length. There is a significant correlation (b) (p-val = 6.4e-05), however, if we exclude the length of the centromere itself, the correlation is gone (c).

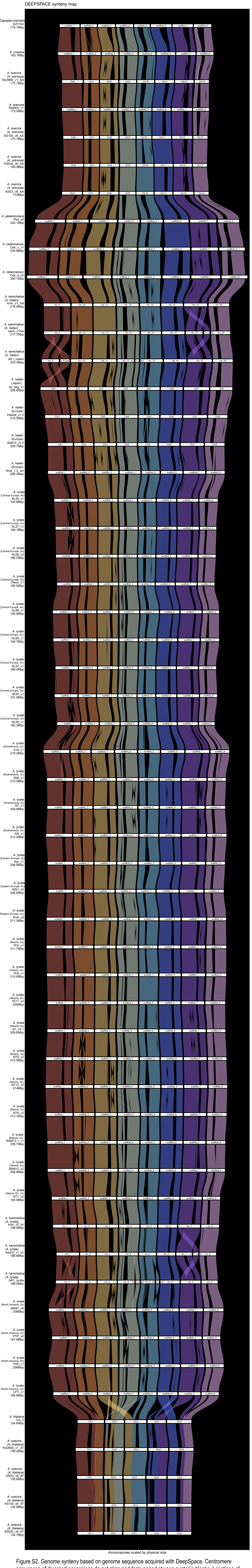


Figure S2. Genome synteny based on genome sequence acquired with DeepSpace. Centromere sequences of diverged accessions do not align and form gaps between syntenic blocks. Locations of these gaps are syntenic between the species.

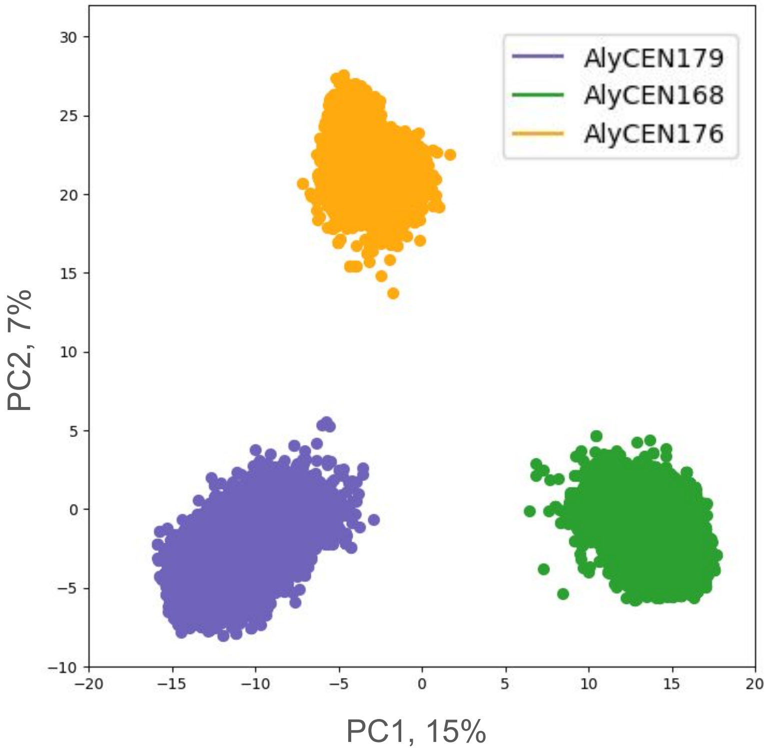


Figure S3. An example PCA of repeat monomers based on 5-mer composition for TE8 sample of *A. lyrata* (Siberian lineage). Repeat type of each cluster was confirmed with BLAST.

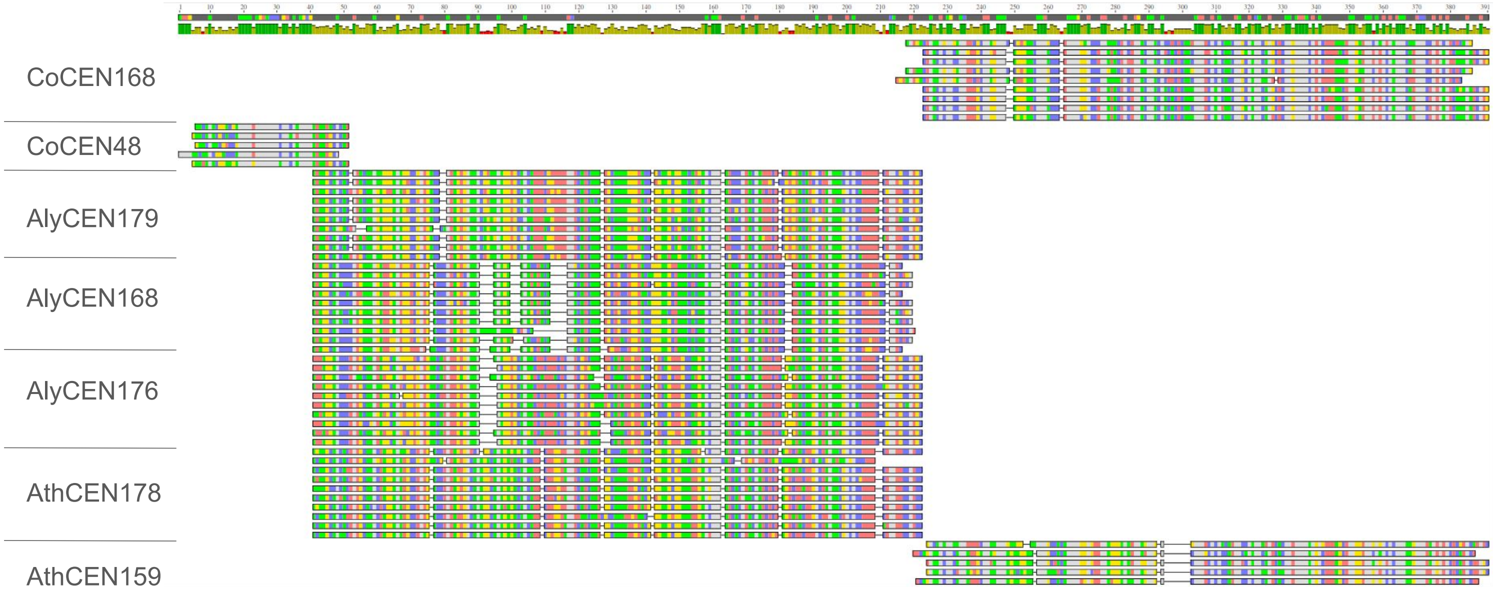


Figure S4. Multiple sequence alignment of randomly sampled repeat monomers from 7 repeat types. The alignment was performed and visualized with Geneious. All main *Arabidopsis* repeat types align together, while AthCEN159 aligns with *Capsella* monomers.

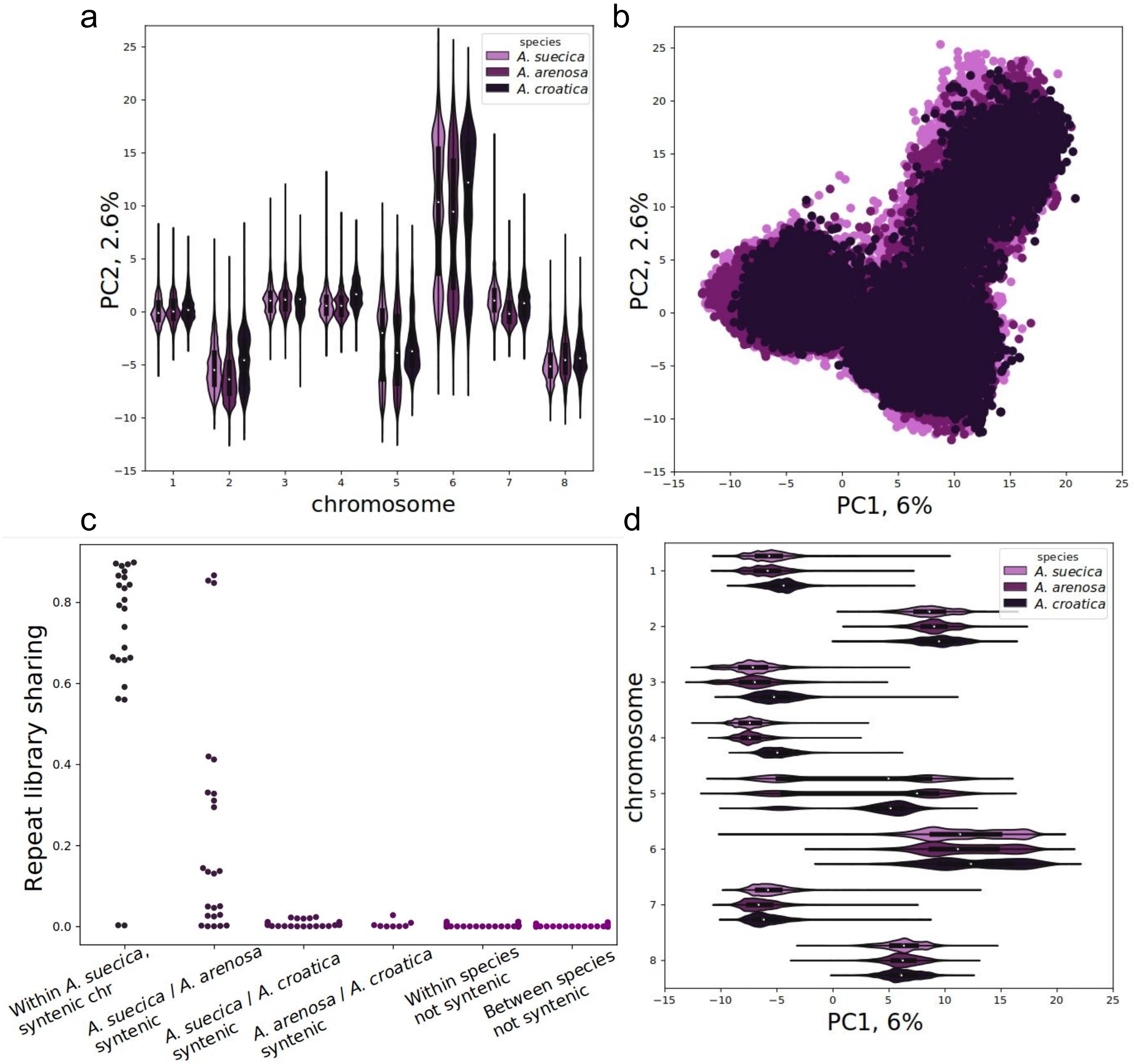


Figure S5. a, b, d) PCA of AlyCEN179 monomers from *A. arenosa* (purple), *A. arenosa* subgenome of *A. suecica* (light purple) and *A. croatica* (dark purple). (a) and (d) represent violin plots of repeat PC values split by the chromosome. The distributions are more similar on the same chromosome of the species than between the chromosomes. c) Proportion of shared repeats between syntenic and non-syntenic centromeres of *A. arenosa*, *A. arenosa* subgenome of *A. suecica* and *A. croatica*. Sharing between syntenic centromeres of any pair of species is significantly higher than sharing between non-syntenic centromeres (p-val < 0.003, MW test).

a

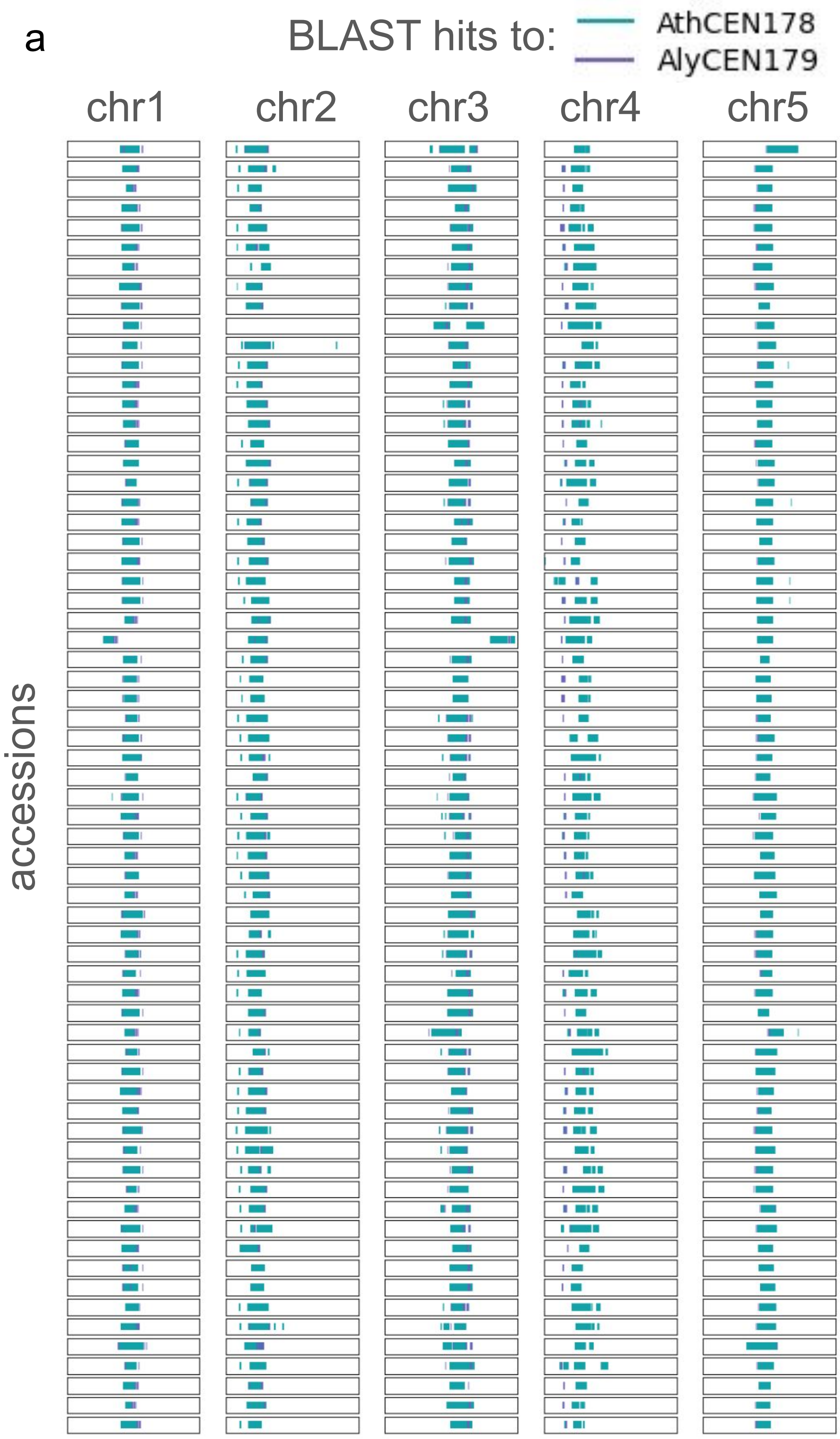


b

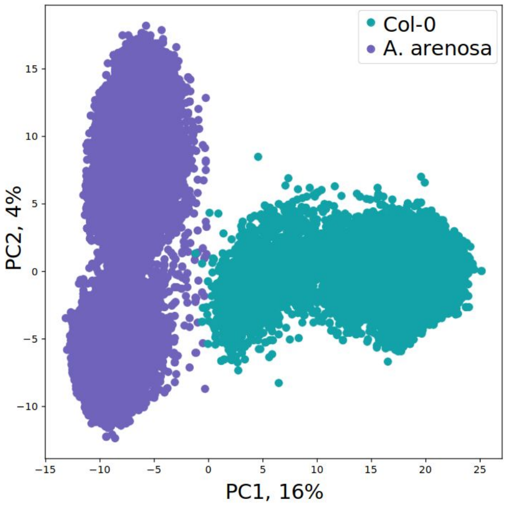


Figure S6. a) BLAST hits of main centromeric repeat types on 66 *A. thaliana* assemblies. Blue - BLAST hit on AthCEN178 repeat, purple - BLAST hit on AlyCEN179 (pAa), representing the intermediate repeat type. Each block represents the whole chromosome, intermediate repeat type occurs on the periphery of the centromeric arrays and pericentromeres. b) PCA of monomers from one *A. arenosa* and one *A. thaliana* genome combined. The majority of repeats are separated, however, some monomers overlap in the intermediate cluster.

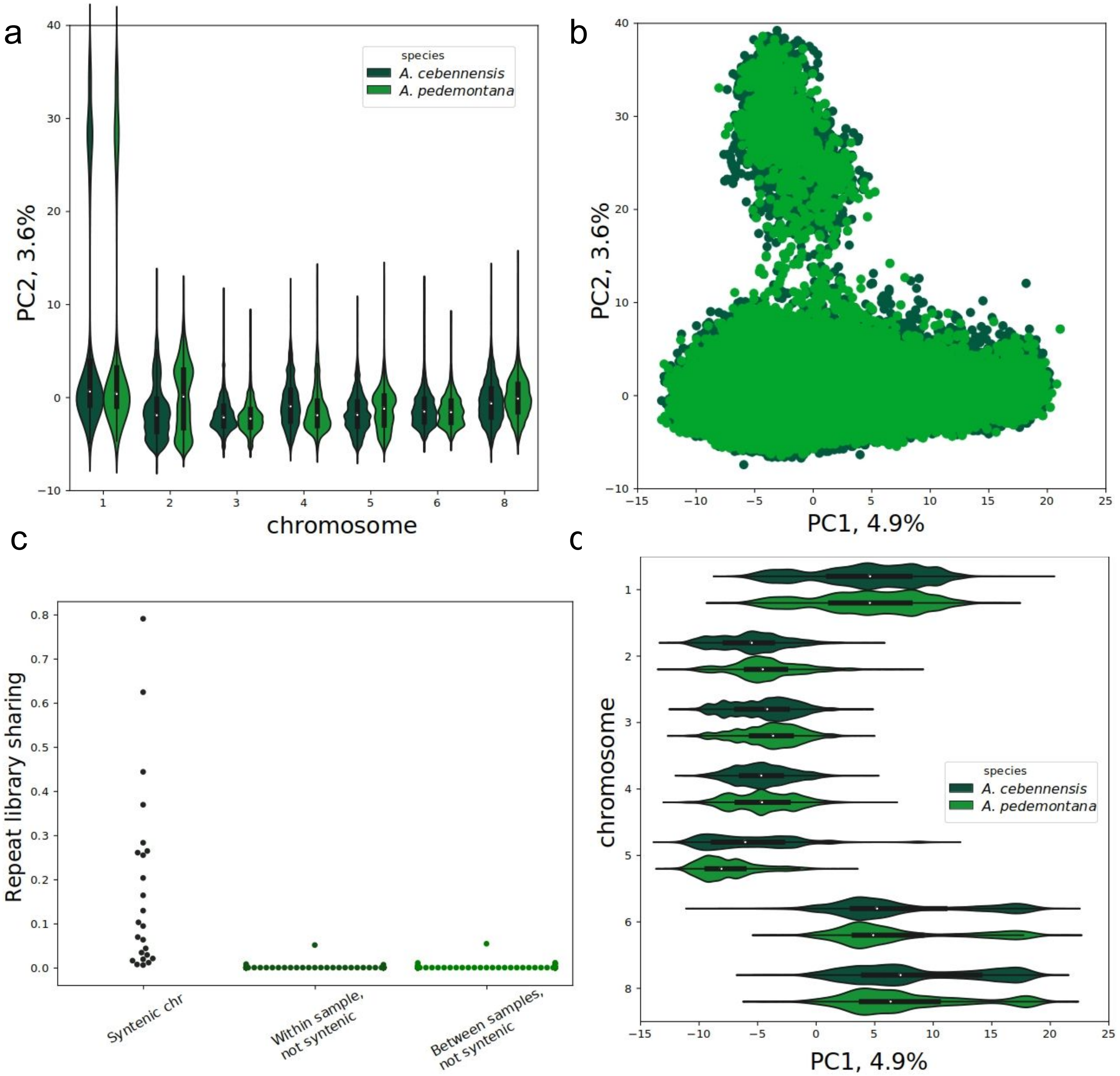


Figure S7. a, b, d) PCA of AlyCEN168 monomers from *A. cebennensis* and *A. pedemontana*. (a) and (d) represent violin plots of repeat PC values split by the chromosome. The distributions are more similar on the same chromosome of the two species than between the chromosomes.

c) Proportion of shared repeats between syntenic and non-syntenic centromeres of *A. cebennensis* and *A. pedemontana*. Sharing between syntenic chromosomes is significantly higher (MW test).

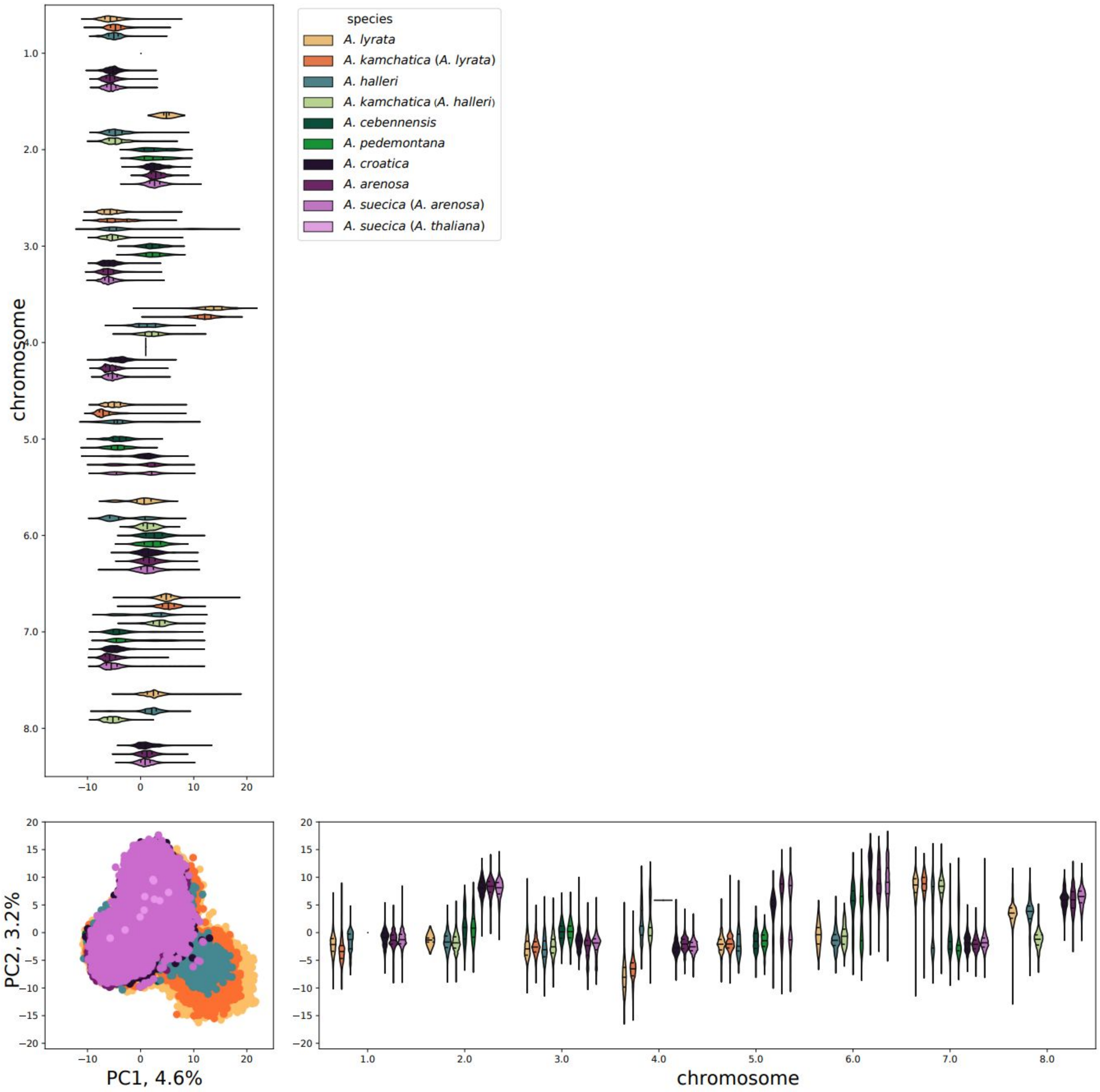


Figure S8. PCA of AlyCEN179 repeats colored by species. The violin plots of repeat PC values split by the chromosome. Even if the centromere has the same repeat type, the k-mer composition can be different. For example, on chr7 *A. lyrata* and *A. halleri* have the majority of repeats with higher PC1 and PC2 values than repeats of *A. arenosa* and *A. croatica*.

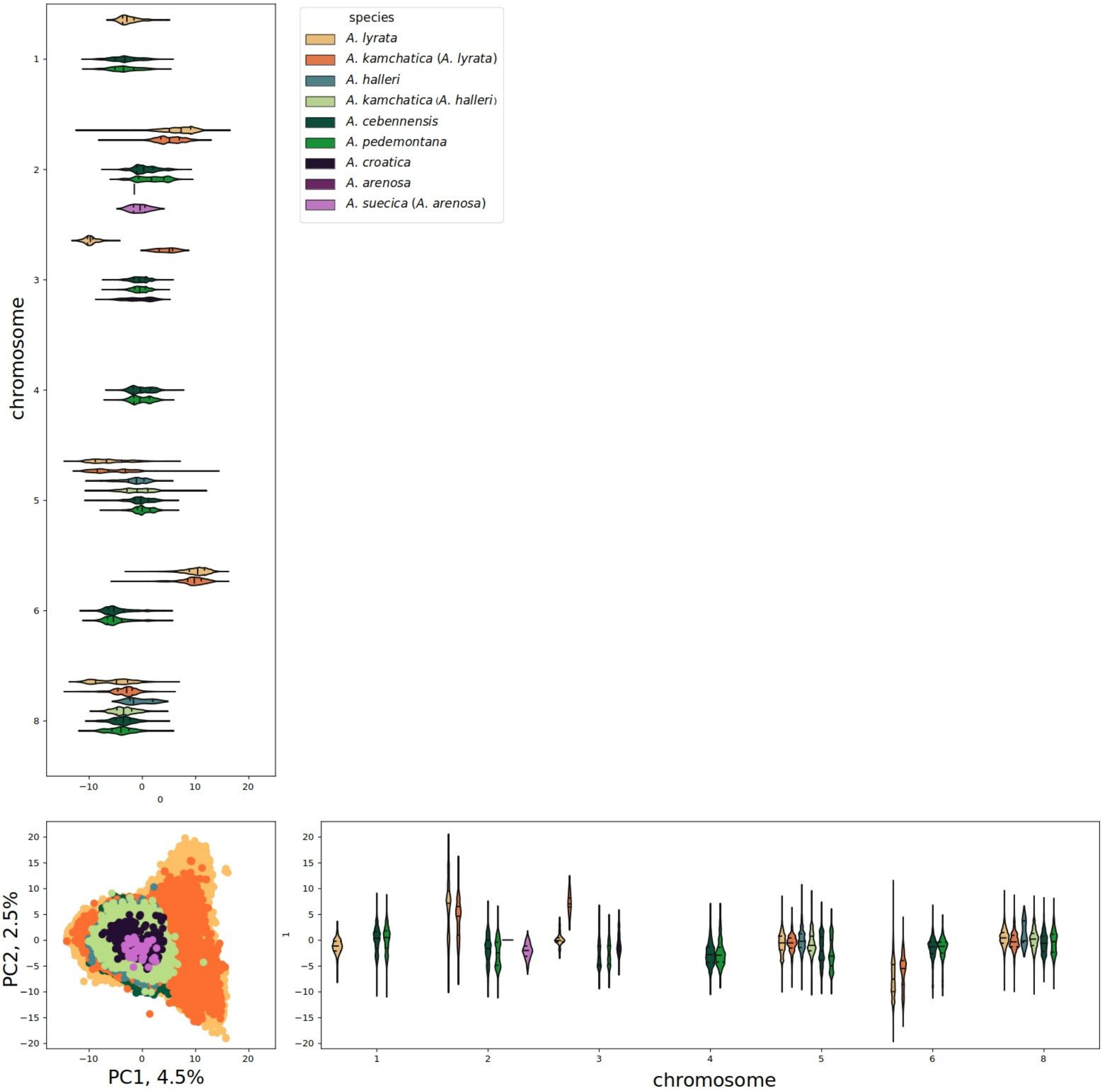


Figure S9. PCA of AlyCEN168 repeats colored by species. The violin plots of repeat PC values split by the chromosome. Even if the centromere has the same repeat type, the k-mer composition can be different. For example, on chromosome 6 *A. cebennensis* and *A. pedemontana* have clearly different PC1 values than *A. lyrata*.

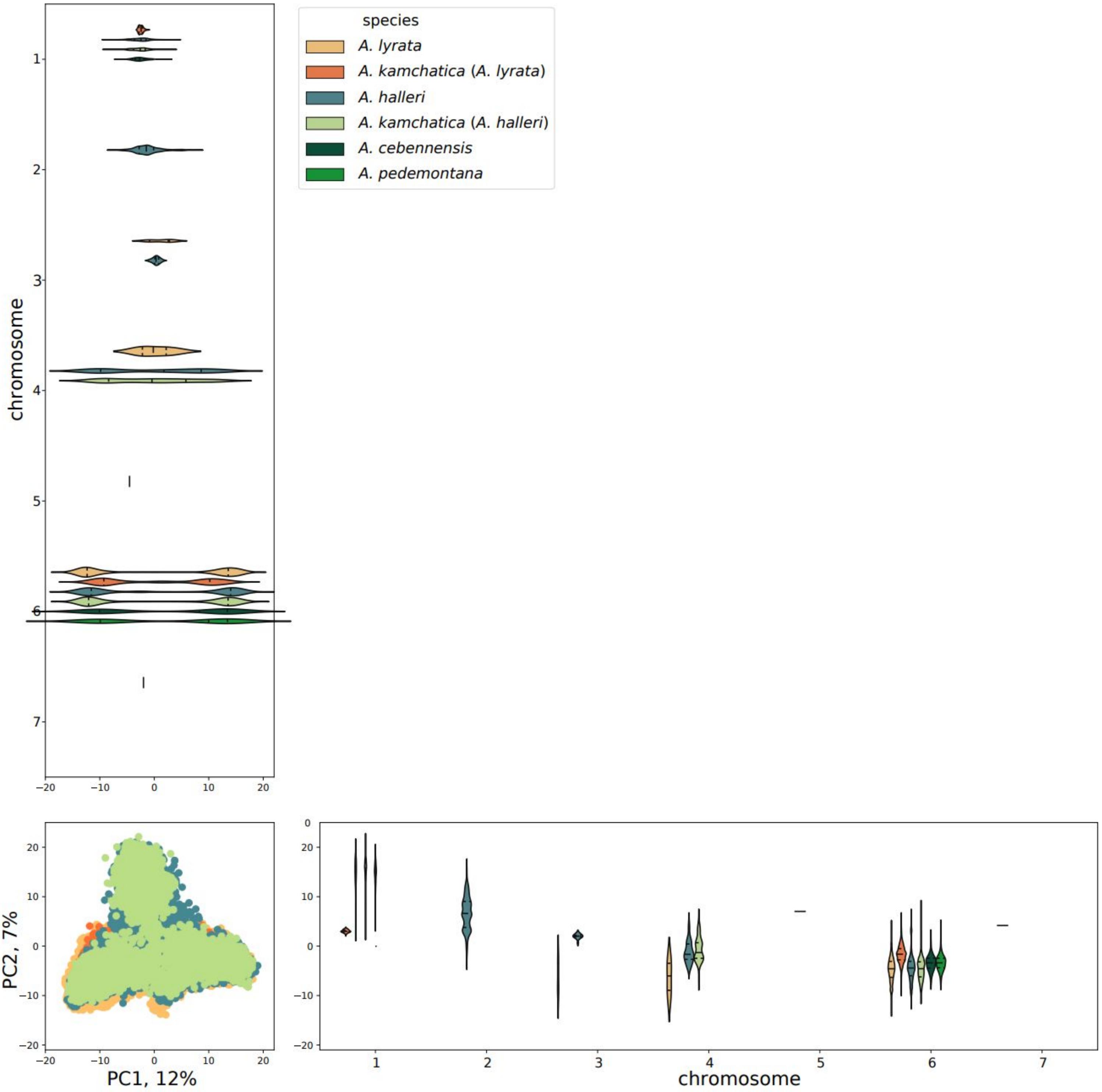


Figure S10. PCA of AlyCEN176 repeats colored by species. The violin plots of repeat PC values split by the chromosome. Chromosome 6 of all species shows highly pronounced dimeric composition with two repeat subclasses (2 clusters by PC1) while this is absent on the other chromosomes.

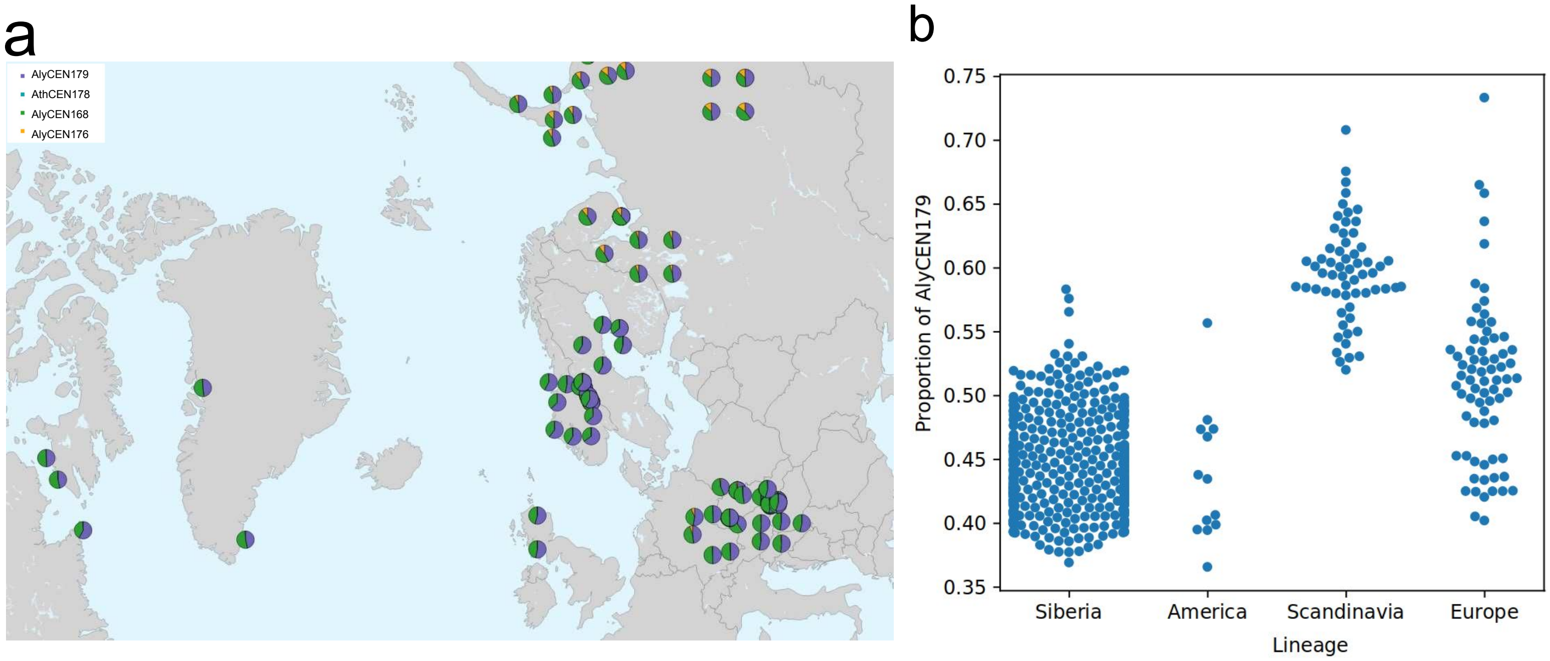


Figure S11. Proportions of short reads mapped on different repeat types. Scandinavian *A. lyrata* shows higher proportions of AlyCEN179 (b) in population in Sweden and Norway (a). This corresponds to the AlyCEN179 on centromere 5 in N7 and S06 genomes. Samples with high AlyCEN179 proportion come from LOI tetraploid population and could acquire additional AlyCEN179 through introgression with *A. arenosa*.

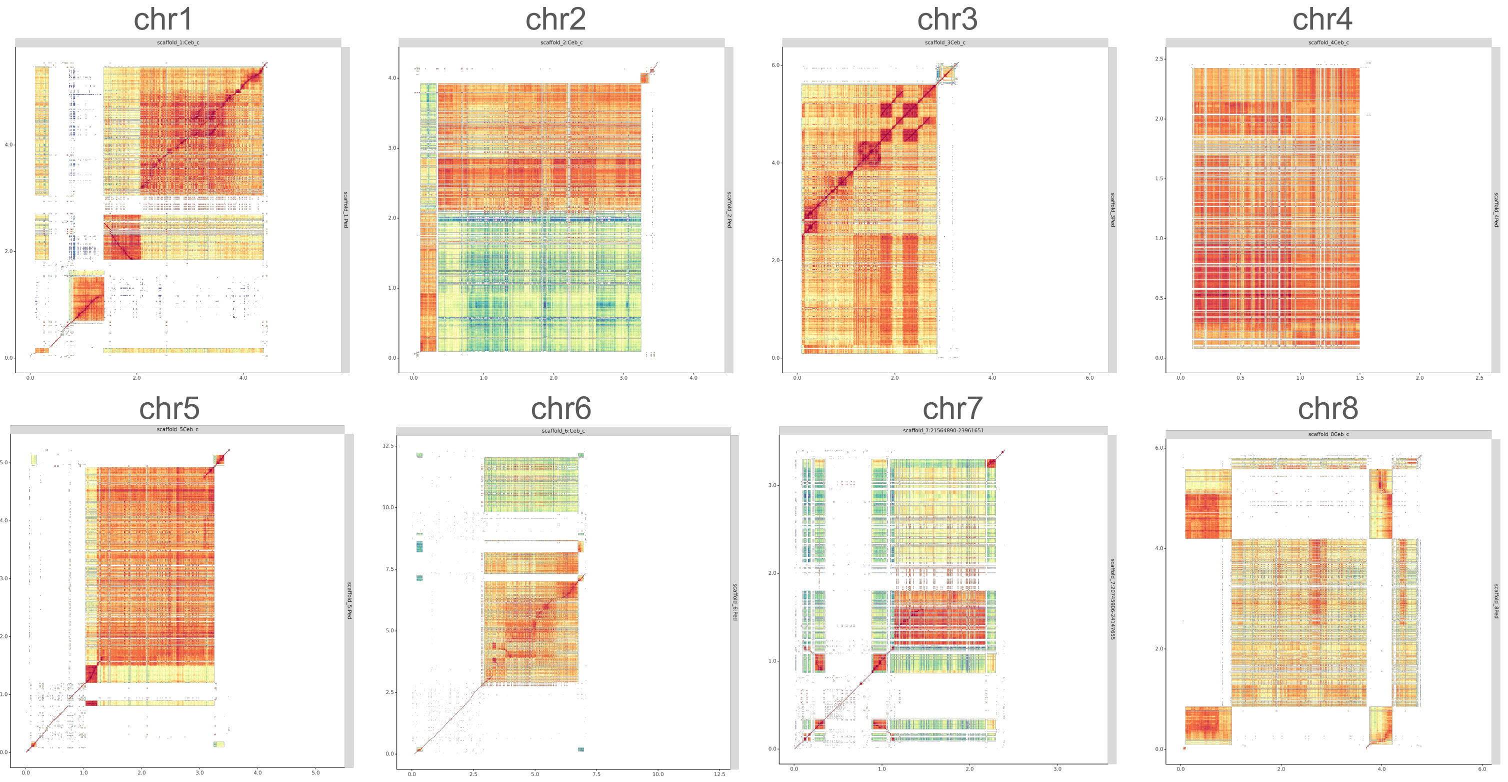


Figure S12. Dotplot of centromeres of *A. pedemontana* versus *A. cebennensis* (sample Ceb_c). Some centromeric regions have highly similar structures, for example, chr3.

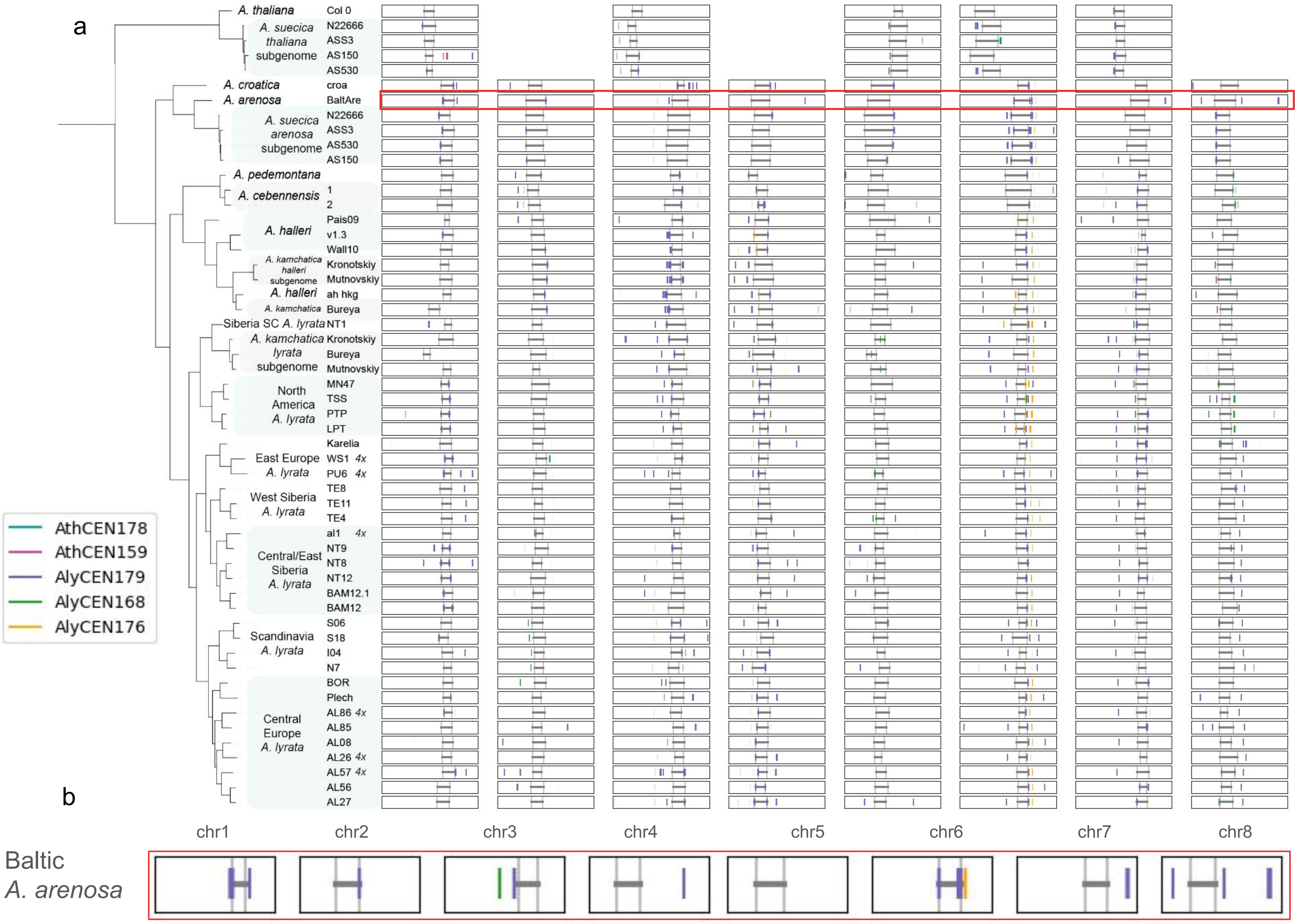

a
A. thaliana
Col 0
A. suecica thaliana subgenome
N22666
ASS3
AS150
AS530
A. croatica
croa
A. arenosa
BaltAre
A. suecica arenosa subgenome
N22666
ASS3
AS530
AS150
A. pedemontana
A. cebennensis
1
2
A. halleri
Pais09
v1.3
Wall10
A. kamchatica halleri subgenome
Kronotskiy
Mutnovskiy
A. halleri
ah hkg
A. kamchatica
Bureya
Siberia SC A. lyrata
NT1
A. kamchatica lyrata subgenome
Kronotskiy
Bureya
Mutnovskiy
North America A. lyrata
MN47
TSS
PTP
LPT
East Europe A. lyrata
Karelia
WS1 4x
PU6 4x
West Siberia A. lyrata
TE8
TE11
TE4
Central/East Siberia A. lyrata
al1 4x
NT9
NT8
NT12
BAM12.1
BAM12
Scandinavia A. lyrata
S06
S18
I04
N7
Central Europe A. lyrata
BOR
Plech
AL86 4x
AL85
AL08
AL26 4x
AL57 4x
AL56
AL27
AthCEN178
AthCEN159
AlyCEN179
AlyCEN168
AlyCEN176
b
chr1
chr2
chr3
chr4
chr5
chr6
chr7
chr8
Baltic A. arenosa

c

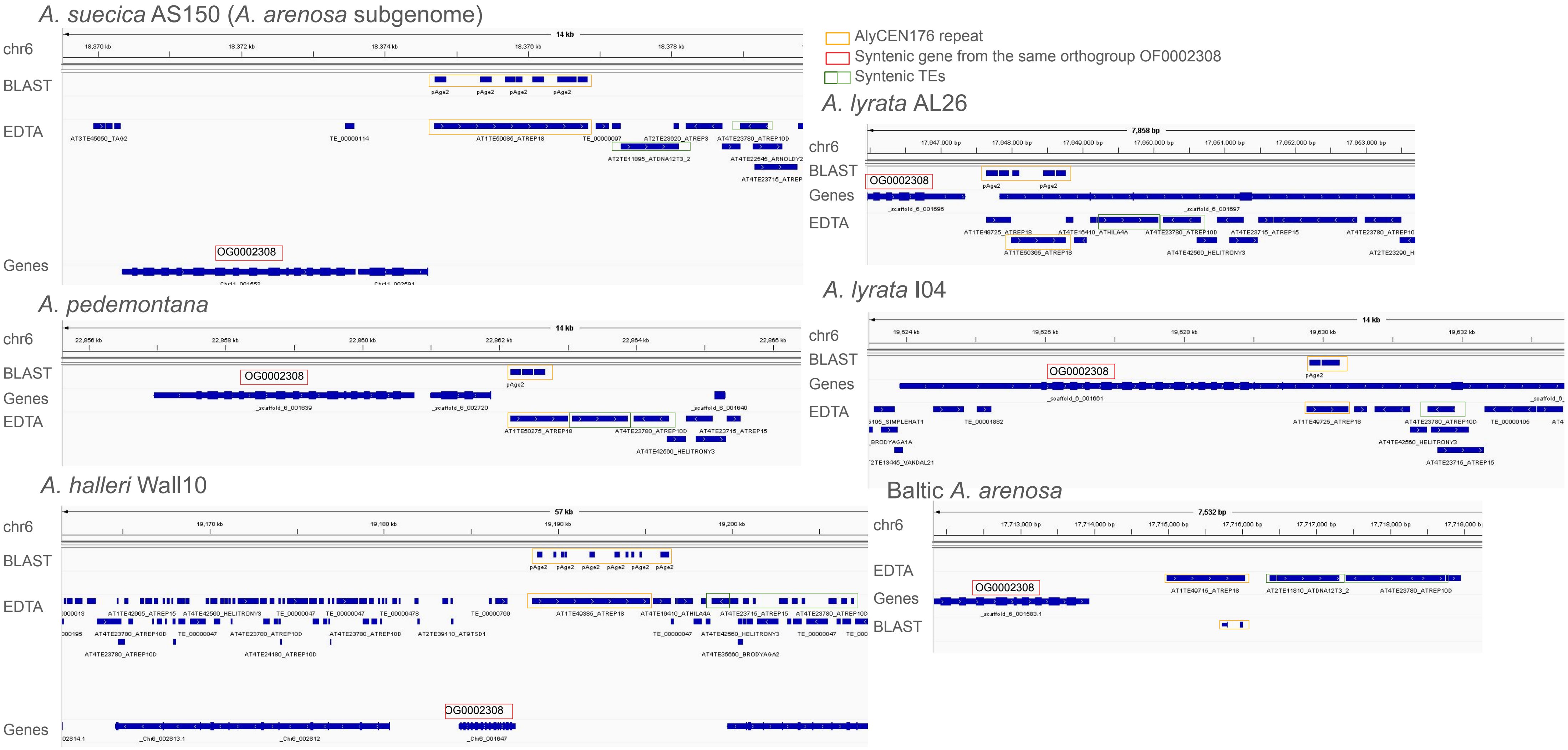


Figure S13. BLAST hits of canonical centromeric repeats outside centromeres. a) Locations of BLAST hits of the centromeric repeats on the chromosomes, each block represents the whole chromosome. Centromere boundaries are shown in gray. b) Bigger representation of BLAST hits in *A. arenosa* genomes. Three repeat types AlyCEN179, AlyCEN168 and AlyCEN176 are present. c) Microsynteny around non-centromeric AlyCEN176 array on chr6. In all species the array is located near a gene from one common orthogroup and two TEs of the same families.

a

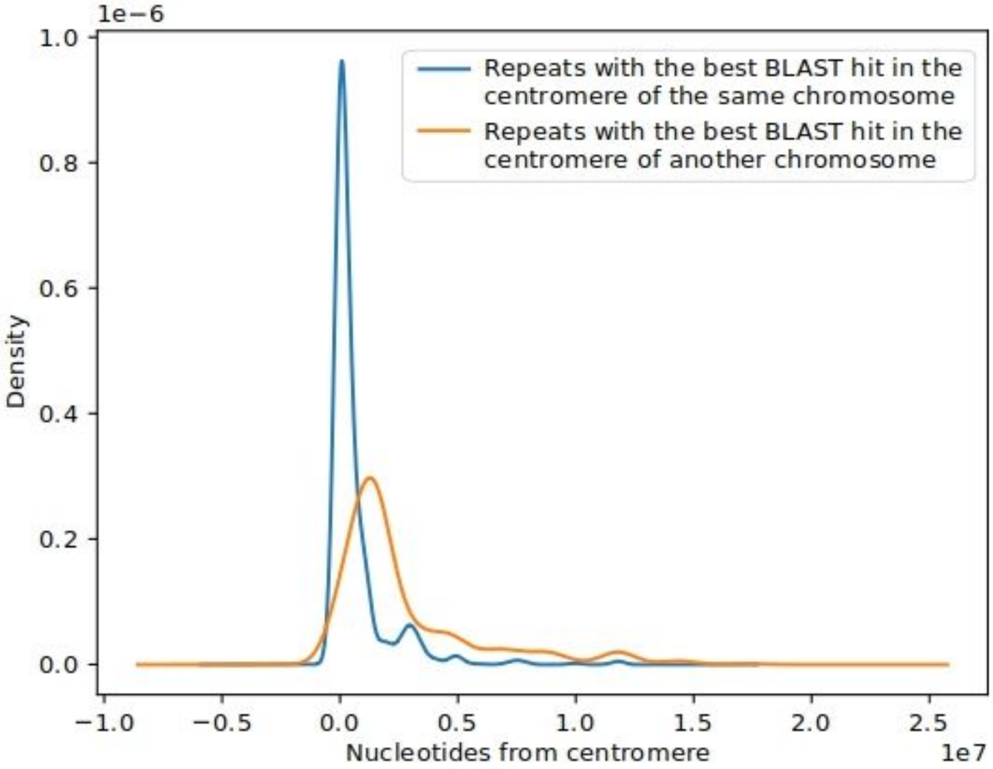


b

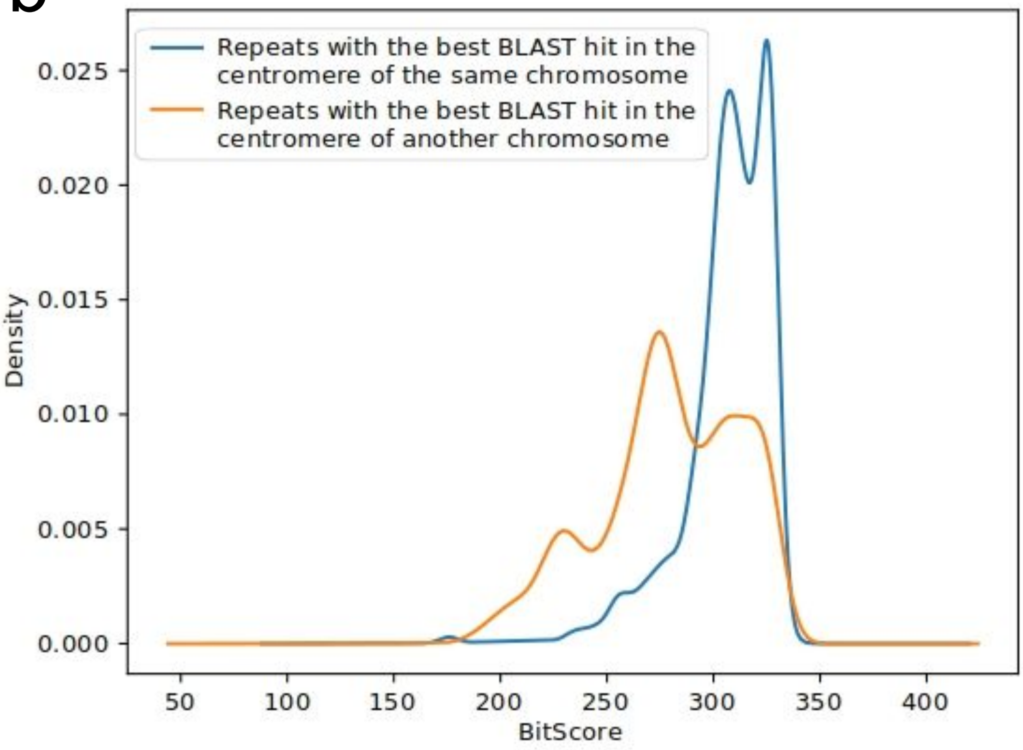


Figure S14. Canonical repeat arrays out of centromeres. Repeats from outside centromeres with the best BLAST hit in the centromere of the same chromosome (a) are located closer to the centromere (MW test, p-value=0.0) and (b) have higher BitScores (MW test, p-value=0).

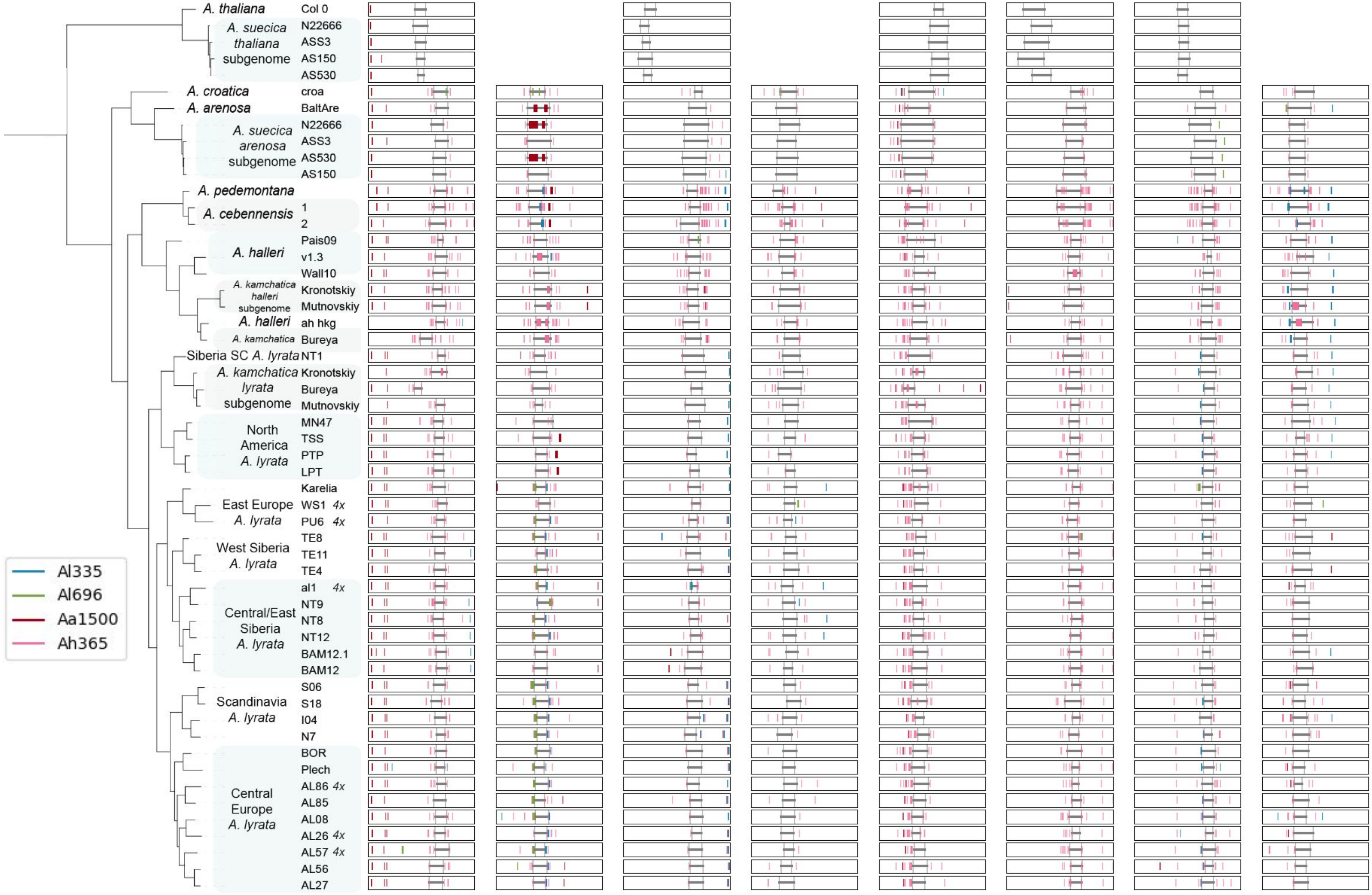


Figure S15. Noncanonical centromeric repeats. Each vertical line represents a BLAST hit. Each block is a whole chromosome. Centromere boundaries are shown in gray.

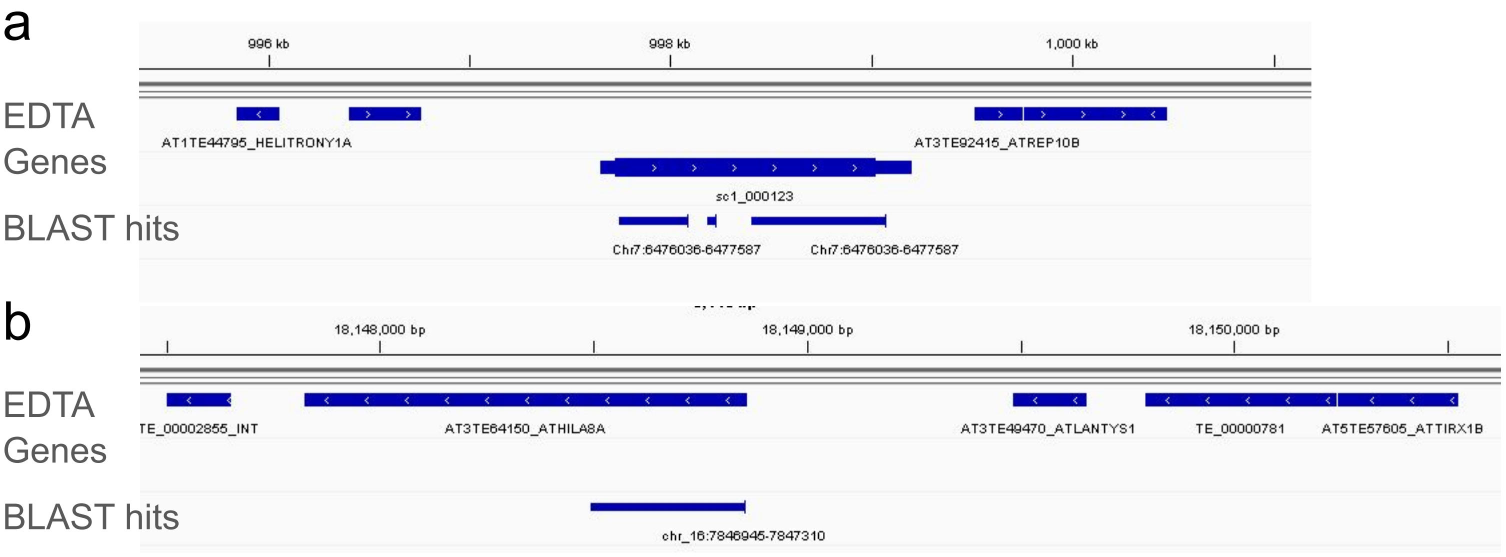


Figure S16. BLAST hits of noncanonical centromeric repeats on chromosome arms. a) Best hit of Aa1500 in NT1 genome, chr1, BitScore=760. b) Best hit of Ah365 in NT1 genome, chr6, BitScore=484

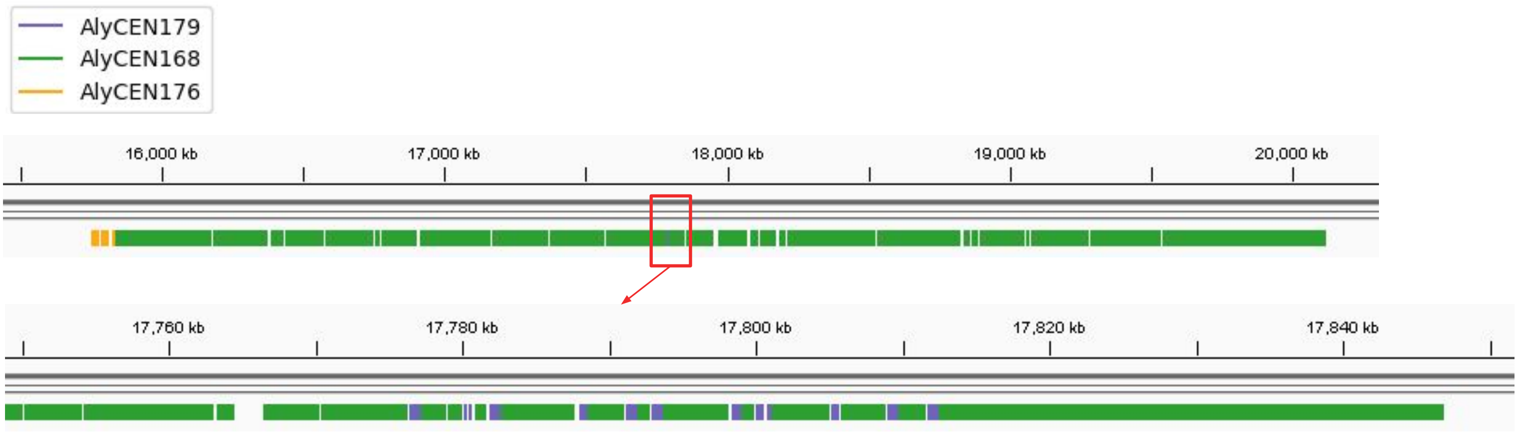


Figure S17. Centromere of chr6 of S18 Scandinavian *A. lyrata*. There are 11 of 2-4 repeat long arrays of AlyCEN179 intermingled with larger arrays of AlyCEN168 which is the main repeat type of the centromere.

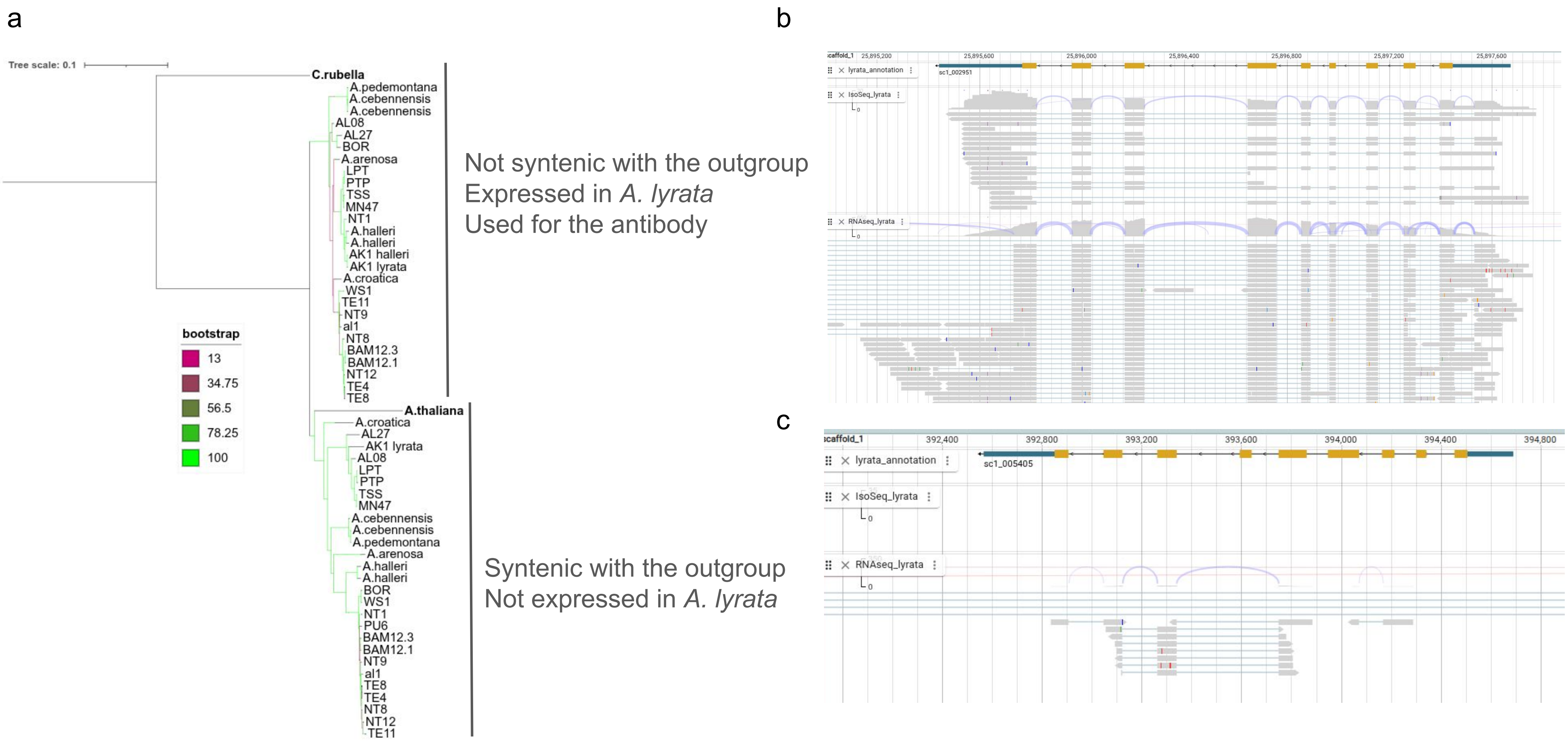


Figure S18. a) Maximum-likelihood tree constructed from DNA sequences of CENH3. Node color represents bootstrap values. *A. lyrata* samples are labeled by their sample ID. b), c) expression in *A. lyrata* NT1 selfing lineage from arabidopsislyrata.org, (b) - AL1G59290 copy, (c) - AL1G11082, a copy syntenic with *A. thaliana* CENH3.

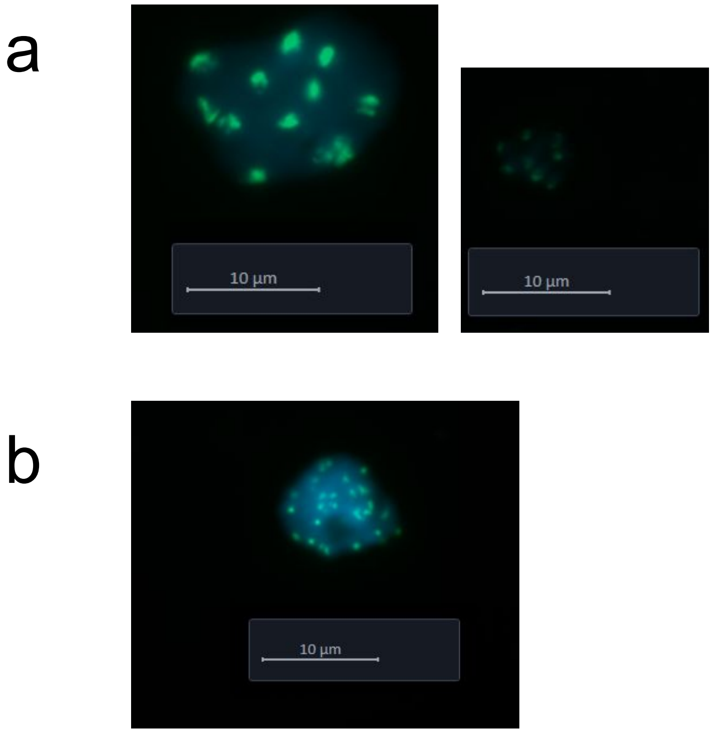


Figure S19. Immunostraining shows CENH3 antibody binding to centromeres. a) Selfing *A. lyrata* NT1, (b) *A. kamchatica*.

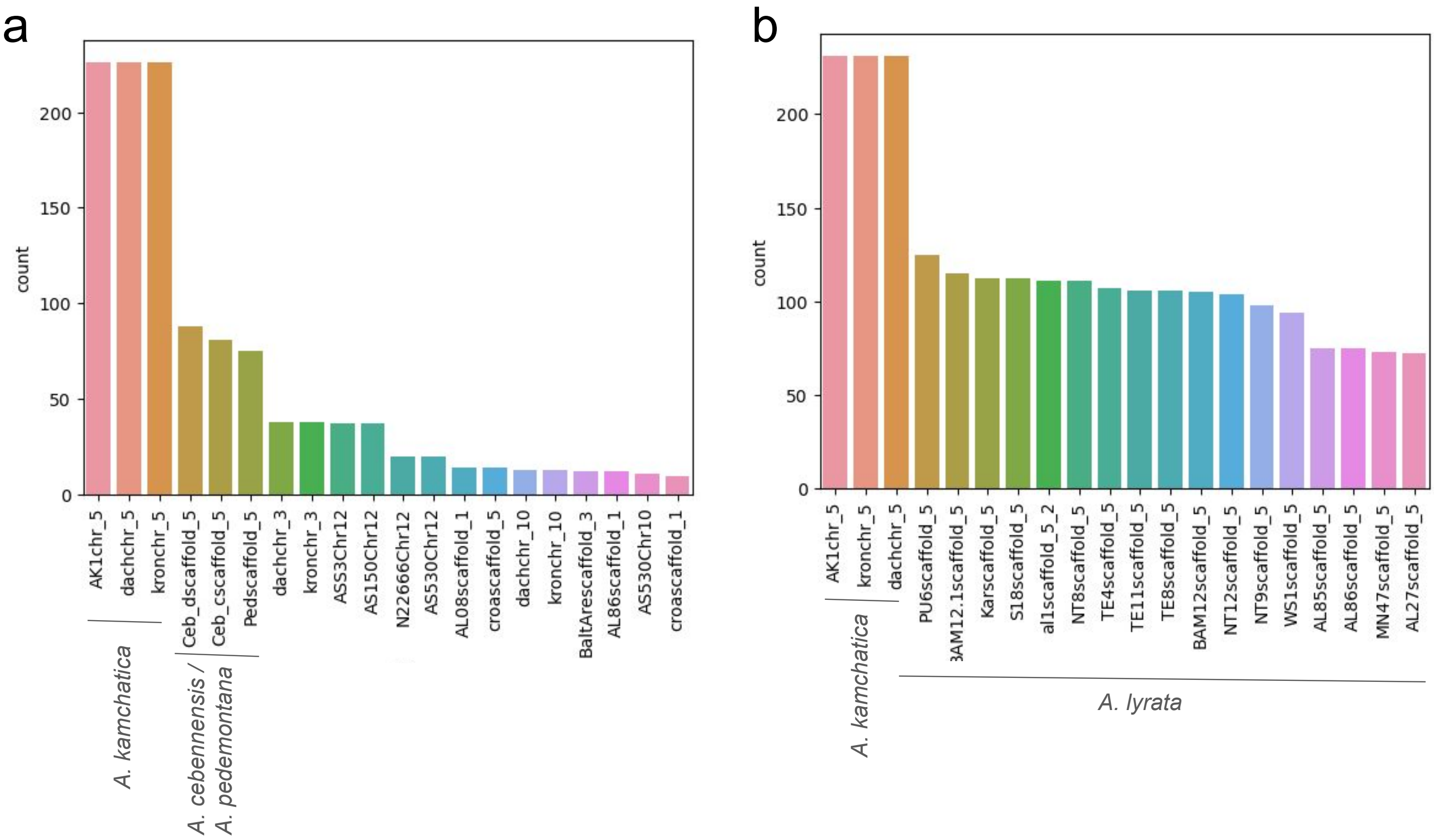


Figure S20. Number of 60-mers from chr5 centromere of Kronotskiy *A. kamchatica* sample, AlyCEN179 array (a) and AlyCEN168 array (b). Samples with the highest number of exact matches of the kmers plotted. Most kmers from the AlyCEN179 array are shared with chr5 of *A. cebennensis* and *A. pedemontana*. Kmers from the AlyCEN168 are highly shared only with *A. lyrata*.

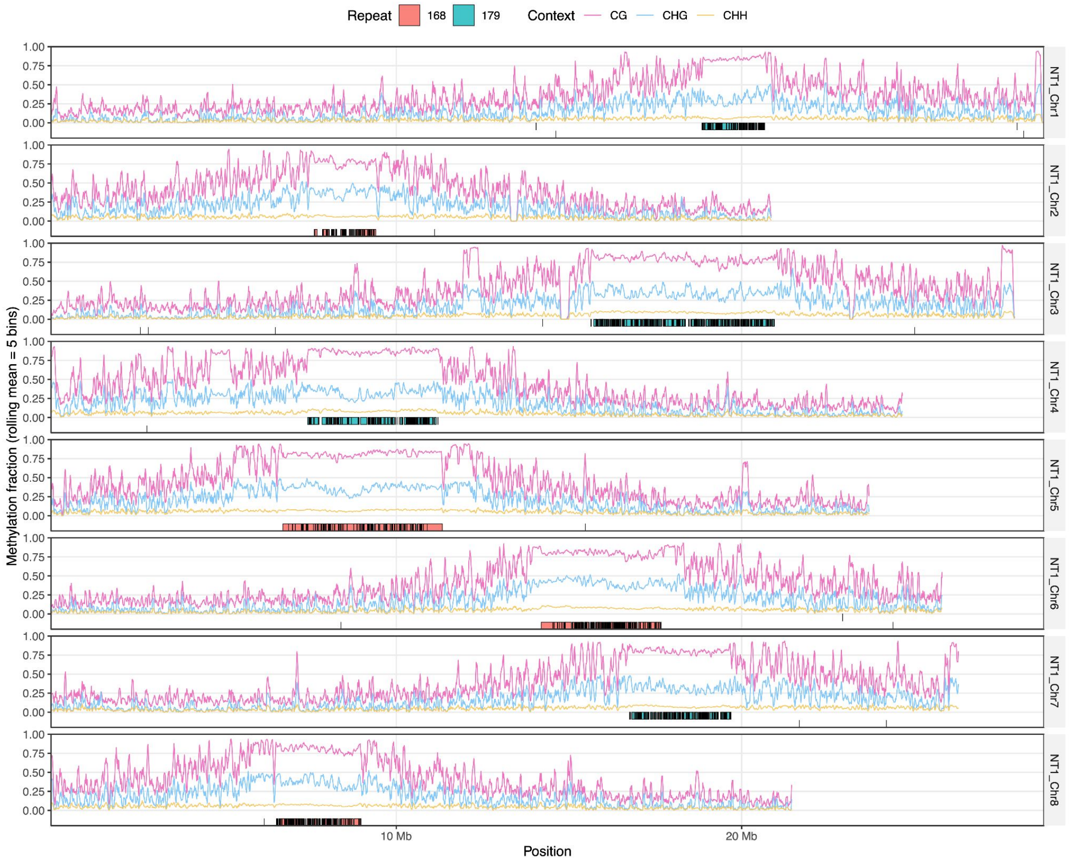


*A. lyrata*

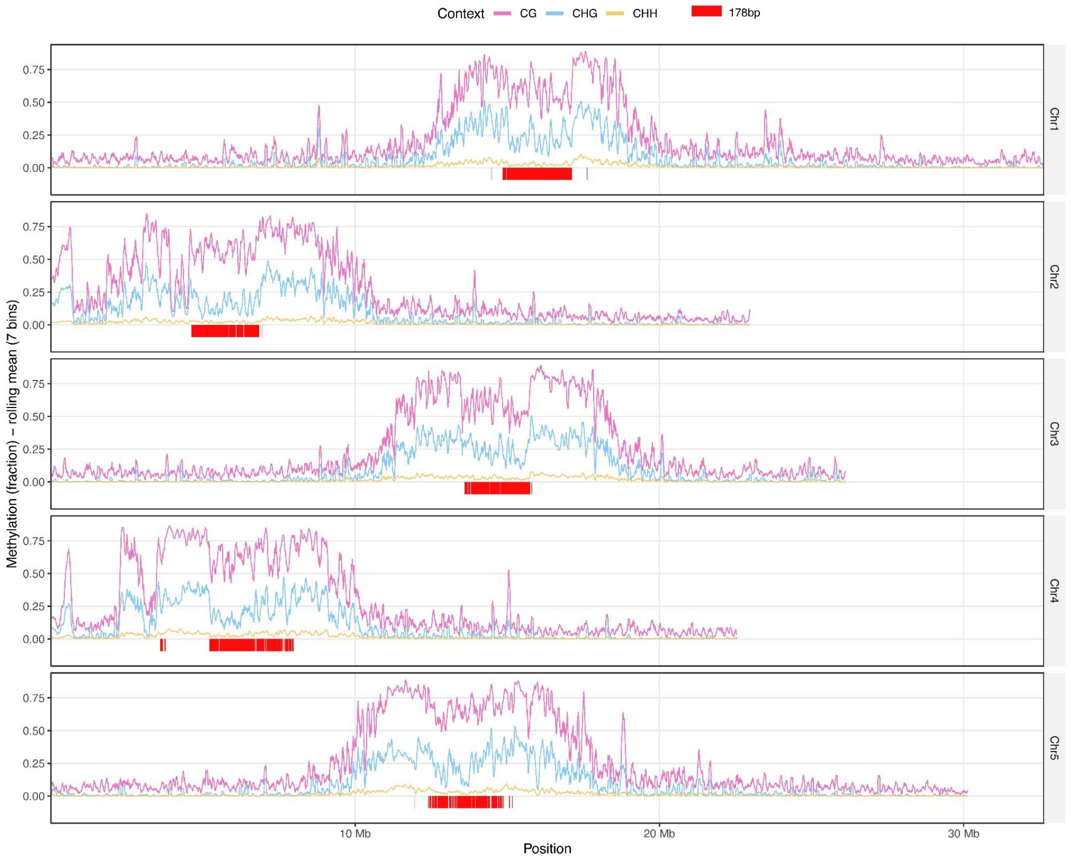


*A. thaliana*

Figure S21. Methylation of NT1 *A. lyrata* chromosomes compared with *A. thaliana* (data from (Burns et al. 2026)). Centromeres of *A. lyrata* are highly methylated, higher than in *A. thaliana*, regions of CENH3 binding with decreased methylation are not pronounced.

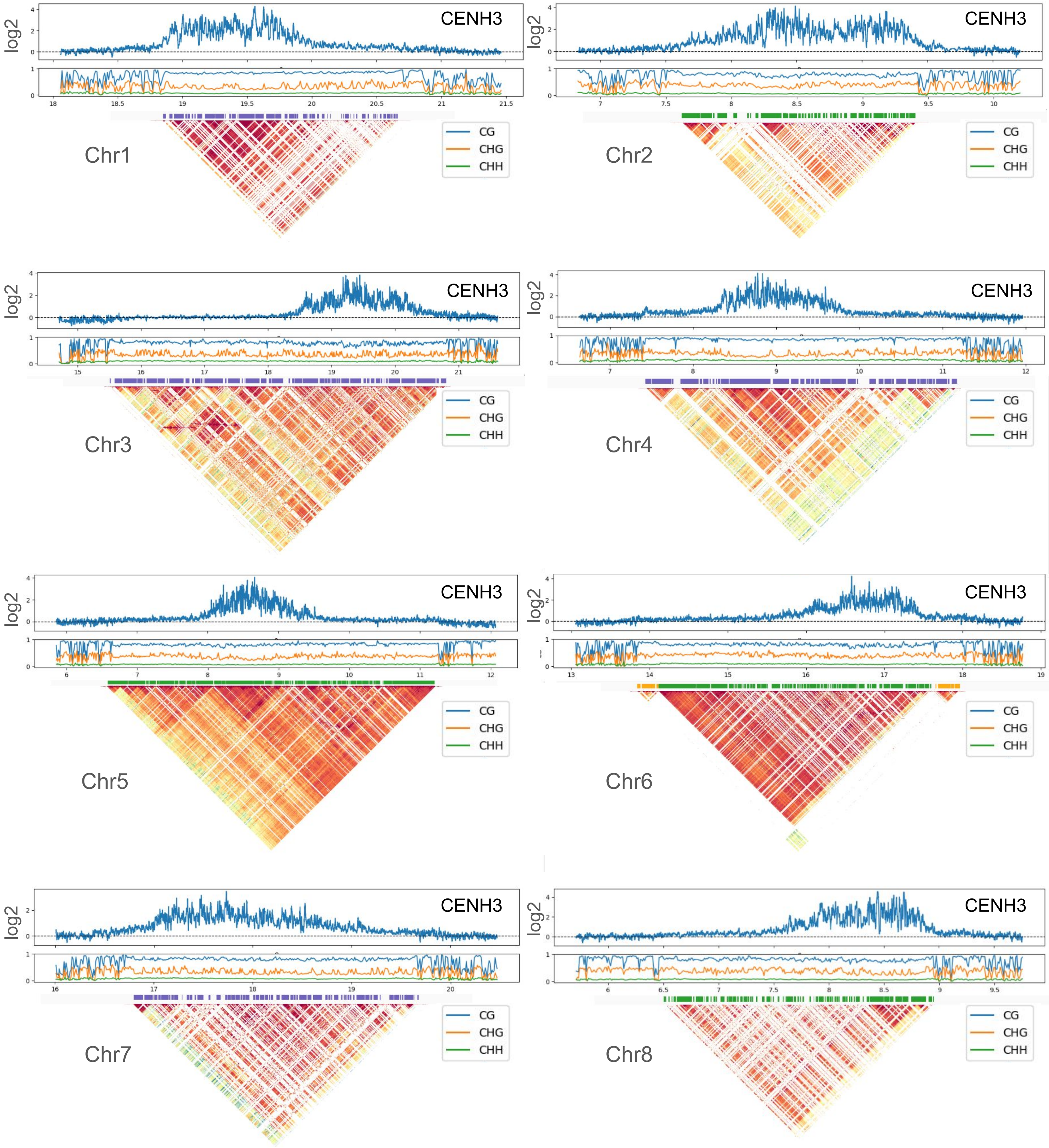


Figure S22. ChIP-seq for CENH3 binding and DNA methylation rate in *A. lyrata* NT1. ChIP-seq log2 ratio and DNA methylation rate are shown in 5 kb windows.

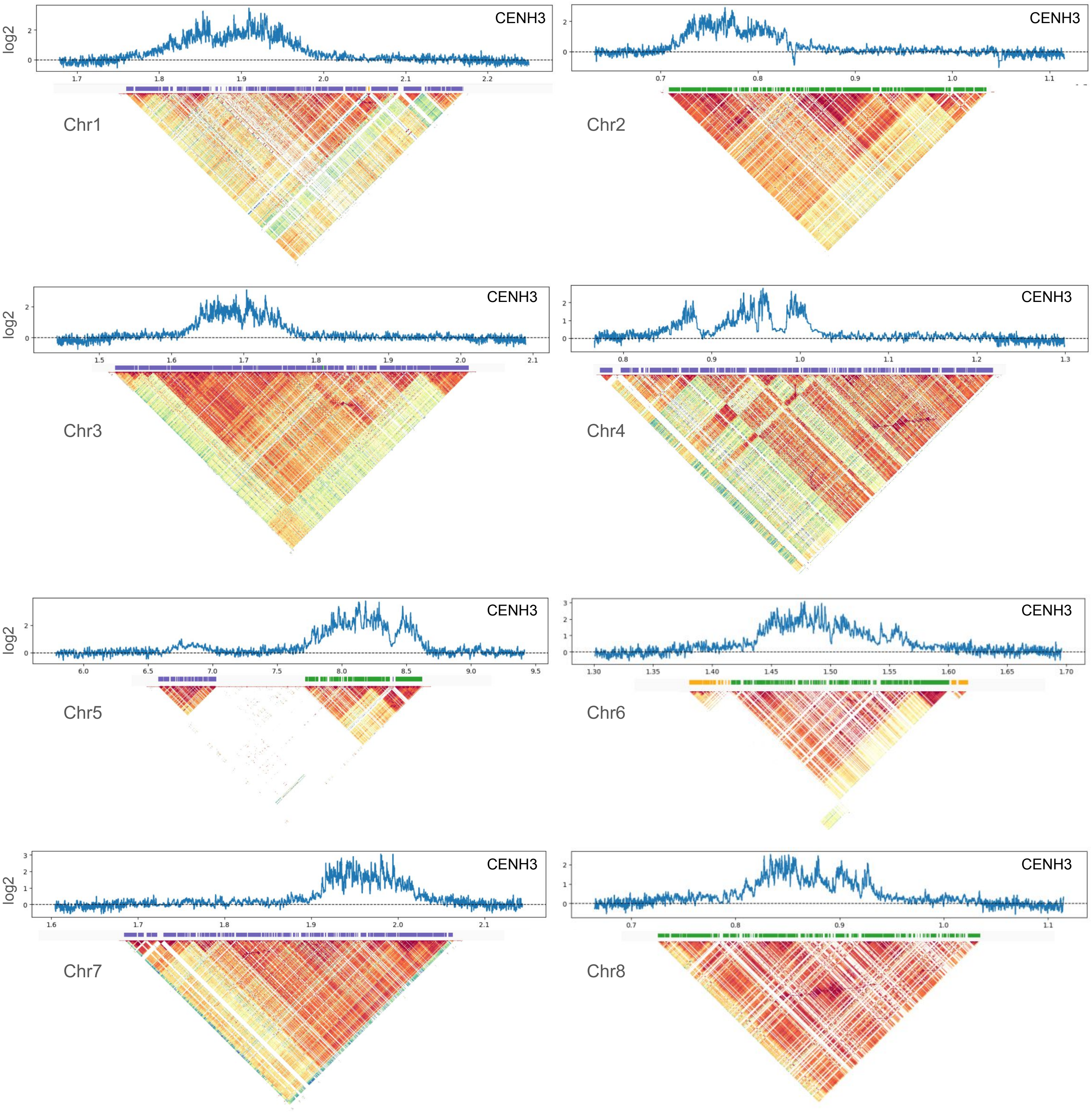


Figure S23. ChIP-seq for CENH3 binding in *A. lyrata* subgenome of Kronotskiy *A. kamchatica*. ChIP-seq log2 ratio is shown in 5 kb windows.

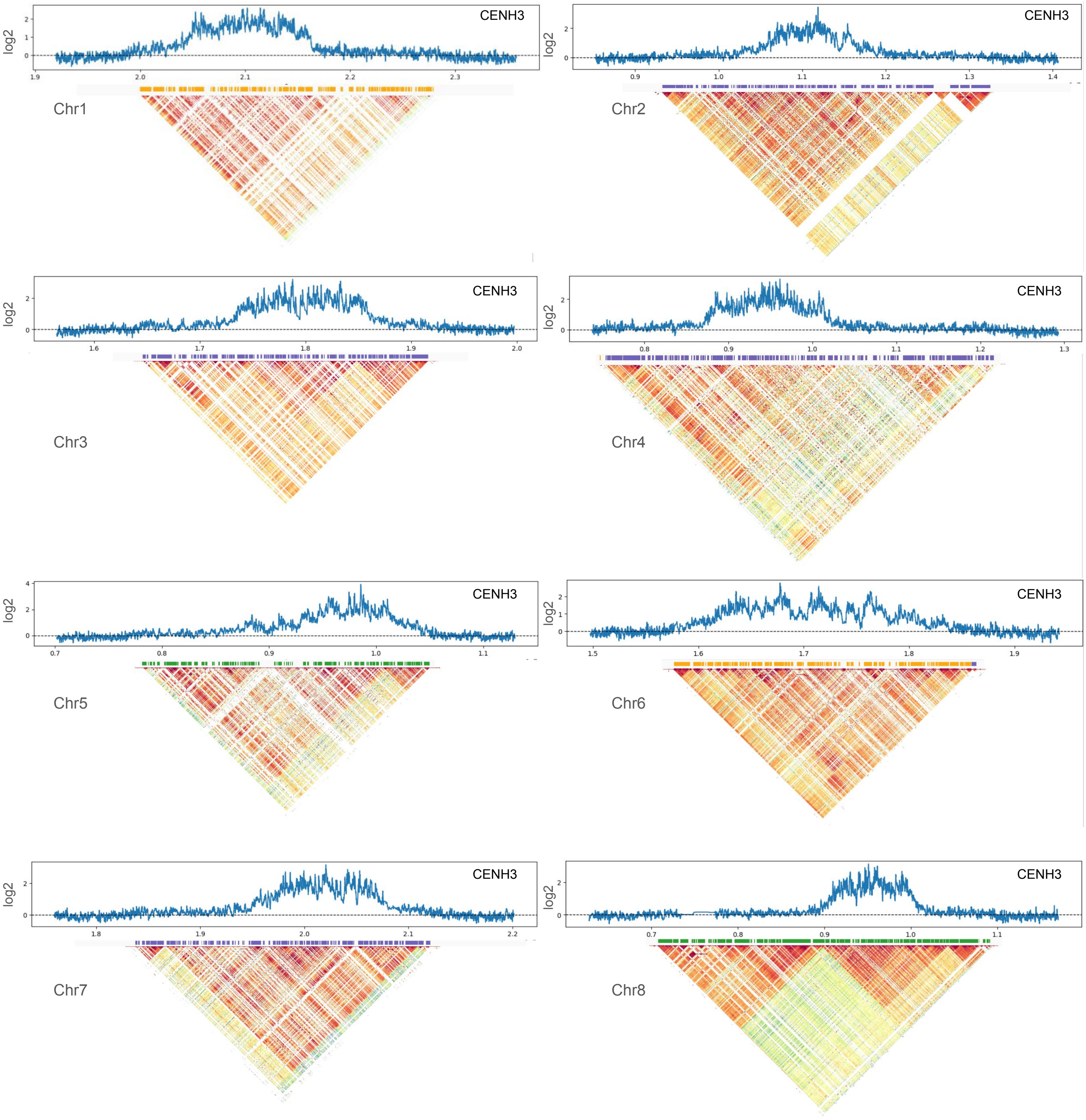


Figure S24. ChIP-seq for CENH3 binding in *A. halleri* subgenome of Kronotskiy *A. kamchatica*. ChIP-seq log2 ratio is shown in 5 kb windows.

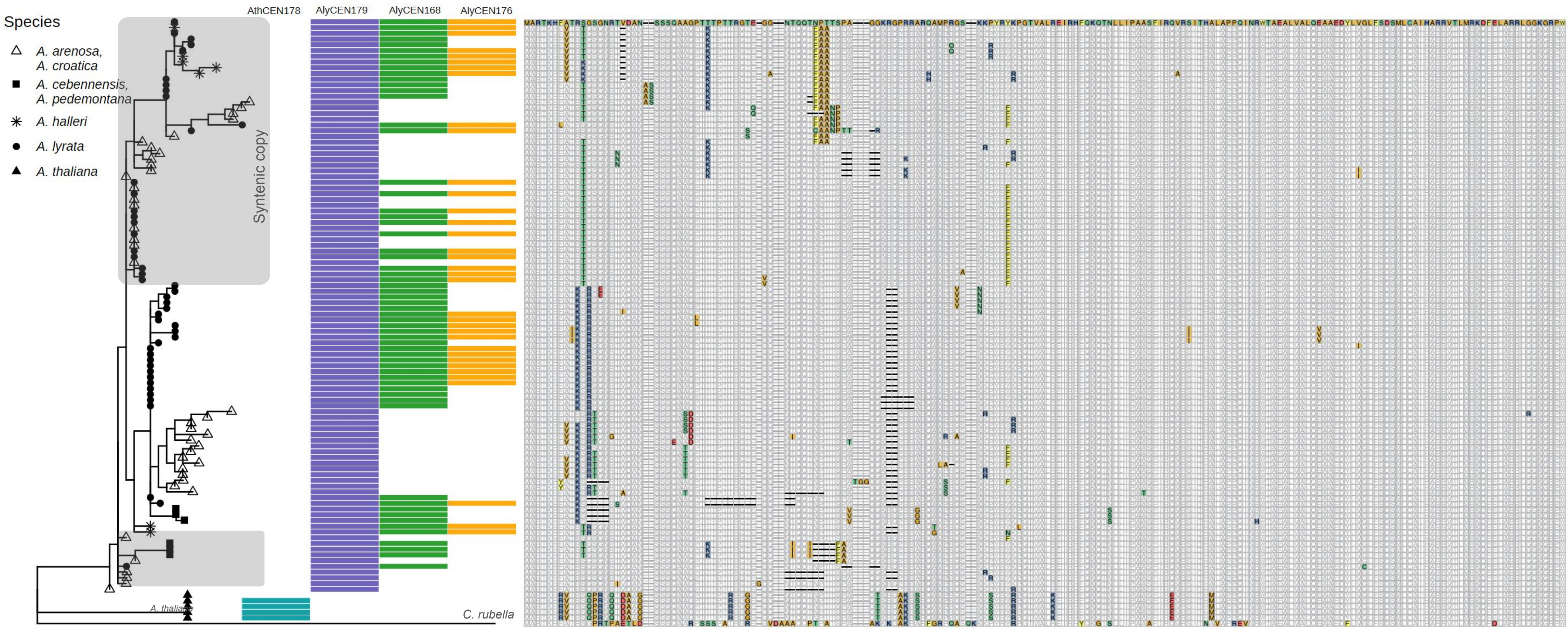


Figure S25. Protein alignment of CENH3 N-tail and a tree based on this alignment. The two copies are different by their protein sequence. The protein sequences of *Capsella* and *A. thaliana* CENH3 have many differences with the rest of the *Arabidopsis* genus, while between other species the variation is less.

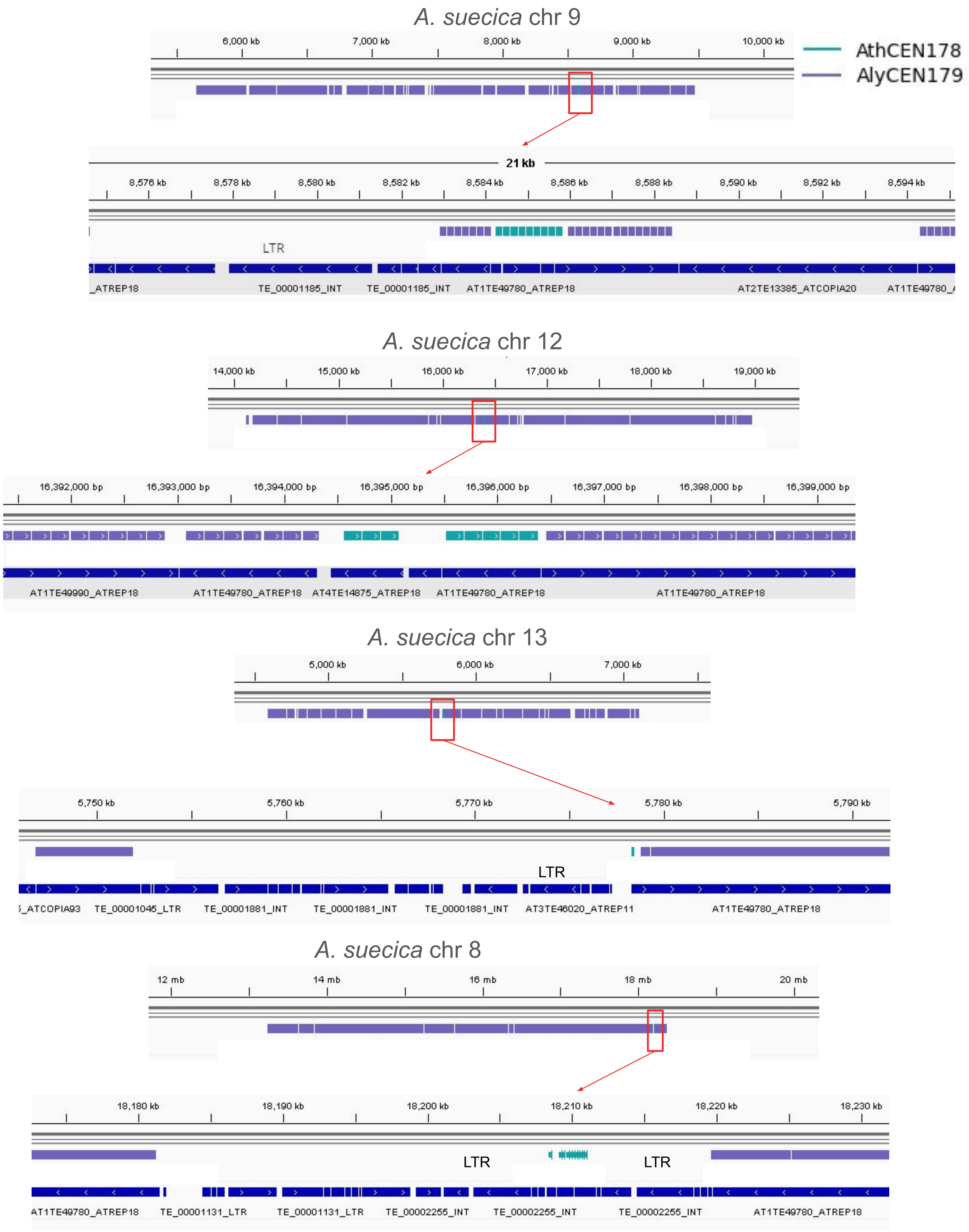

A. suecica chr 9
6,000 kb
7,000 kb
8,000 kb
9,000 kb
10,000 kb
AthCEN178
AlyCEN179
21 kb
8,576 kb
8,578 kb
8,580 kb
8,582 kb
8,584 kb
8,586 kb
8,588 kb
8,590 kb
8,592 kb
8,594 kb
LTR
_ATREP18
TE_00001185_INT
TE_00001185_INT
AT1TE49780_ATREP18
AT2TE13385_ATCOPIA20
A. suecica chr 12
14,000 kb
15,000 kb
16,000 kb
17,000 kb
18,000 kb
19,000 kb
16,392,000 bp
16,393,000 bp
16,394,000 bp
16,395,000 bp
16,396,000 bp
16,397,000 bp
16,398,000 bp
16,399,000 bp
AT1TE49990_ATREP18
AT1TE49780_ATREP18
AT4TE14875_ATREP18
AT1TE49780_ATREP18
AT1TE49780_ATREP18
A. suecica chr 13
5,000 kb
6,000 kb
7,000 kb
5,750 kb
5,760 kb
5,770 kb
5,780 kb
5,790 kb
LTR
_ATCOPIA93
TE_00001045_LTR
TE_00001881_INT
TE_00001881_INT
TE_00001881_INT
AT3TE46020_ATREP11
AT1TE49780_ATREP18
A. suecica chr 8
12 mb
14 mb
16 mb
18 mb
20 mb
18,180 kb
18,190 kb
18,200 kb
18,210 kb
18,220 kb
18,230 kb
LTR
LTR
AT1TE49780_ATREP18
TE_00001131_LTR
TE_00001131_LTR
TE_00002255_INT
TE_00002255_INT
TE_00002255_INT
AT1TE49780_ATREP18

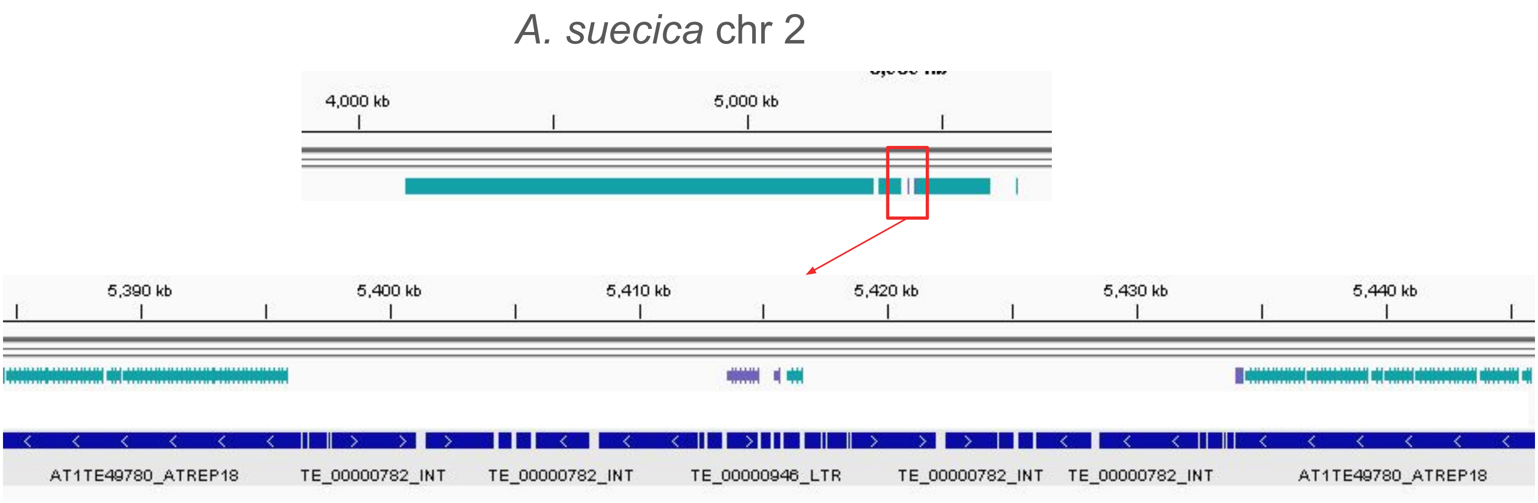


Figure S26. AthCEN178 arrays in the *A. arenosa* subgenome of *A. suecica* (chr 8, 9, 12, 13) and AlyCEN179 array on chr 2 of *A. thaliana* subgenome of *A. suecica*. The small arrays of repeat type from the other subgenome represent jumps that happened after *A. suecica* origin.

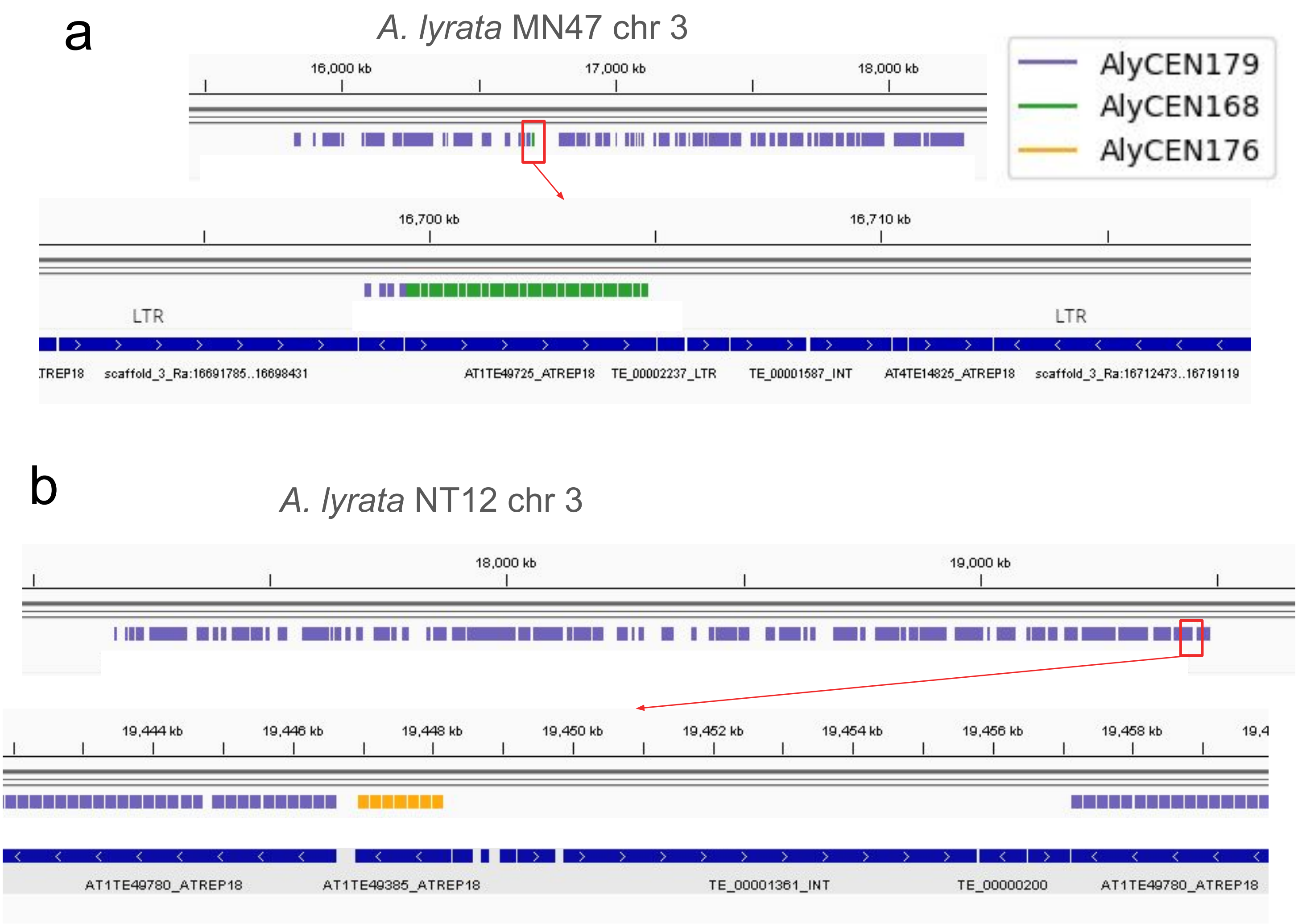


Figure S27. Examples of arrays of atypical repeat types in *A. lyrata* centromeres.

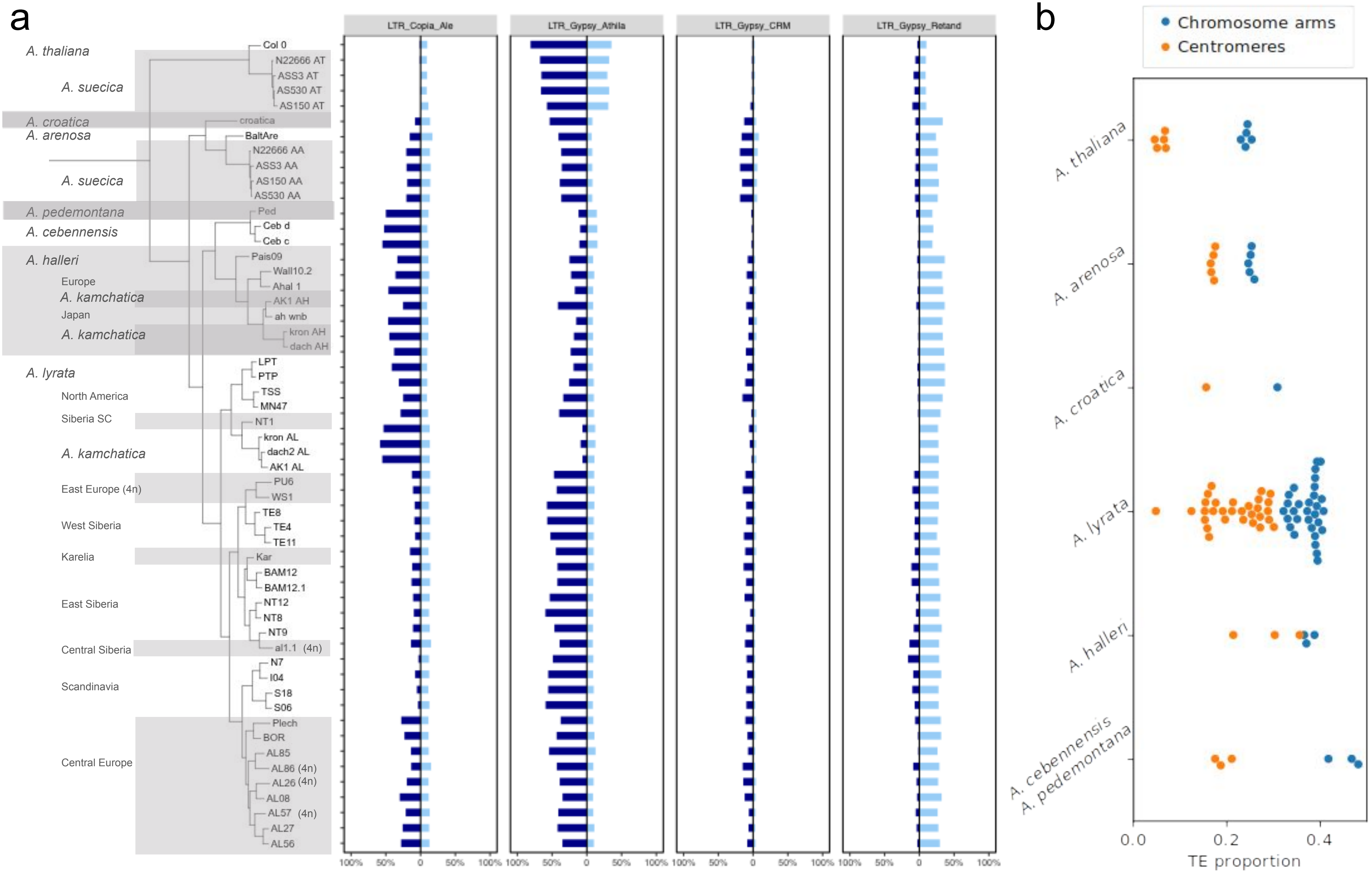


Figure S28. a) Proportion of LTRs of different subfamilies in the centromeric arrays (dark blue) and in the chromosome arms (light blue). b) Total portion of TEs in the genome excluding the centromeres and in the centromeres. *A thaliana* and *A. arenosa* have the least TEs in the chromosome arms as well as in the centromeres.

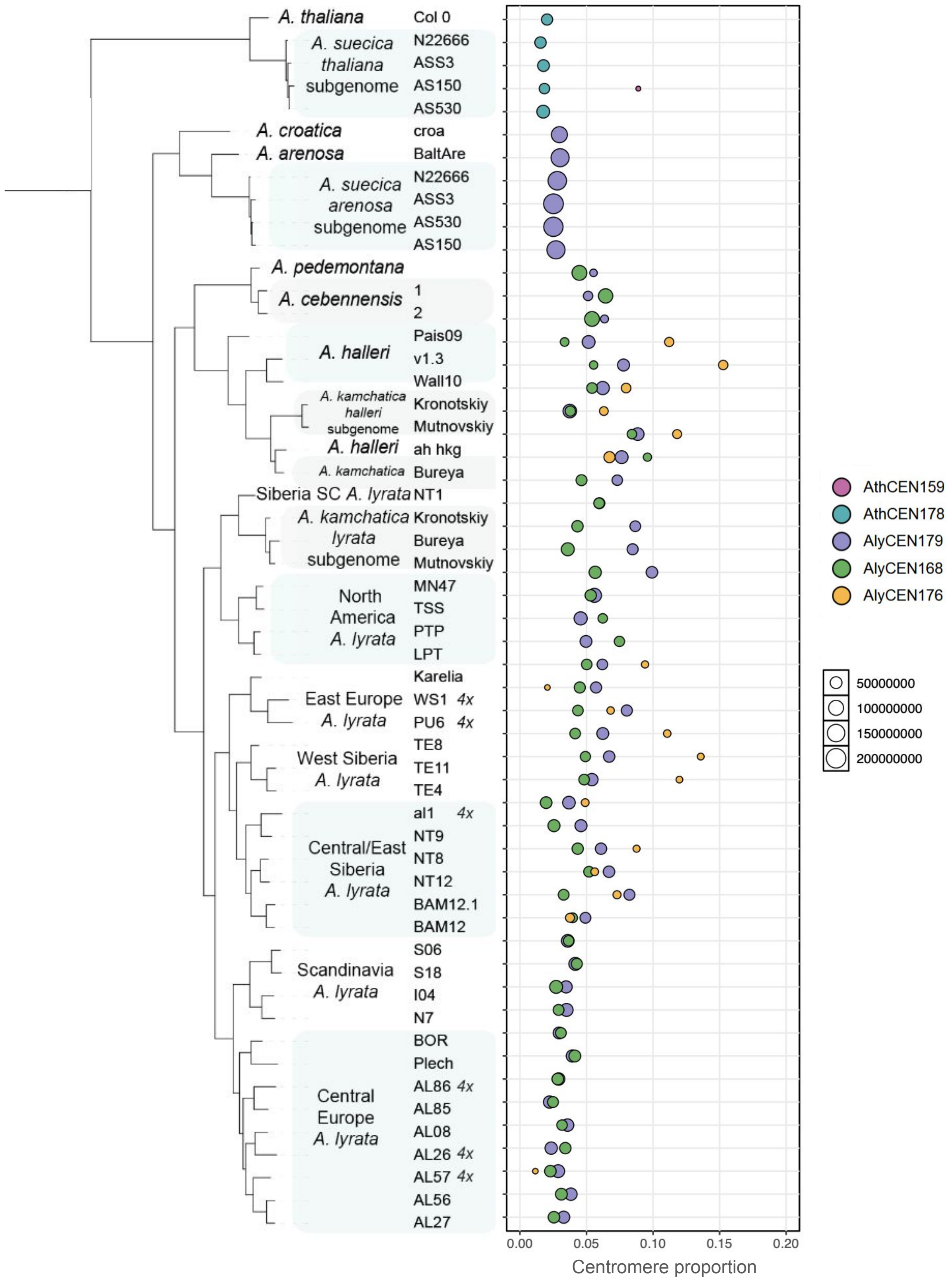


Figure S29. Summarized TE proportion in the centromeres of each sample, split and coloured by the repeat type of the centromeric arrays. The size of the bubble corresponds to the summarized size of the arrays.

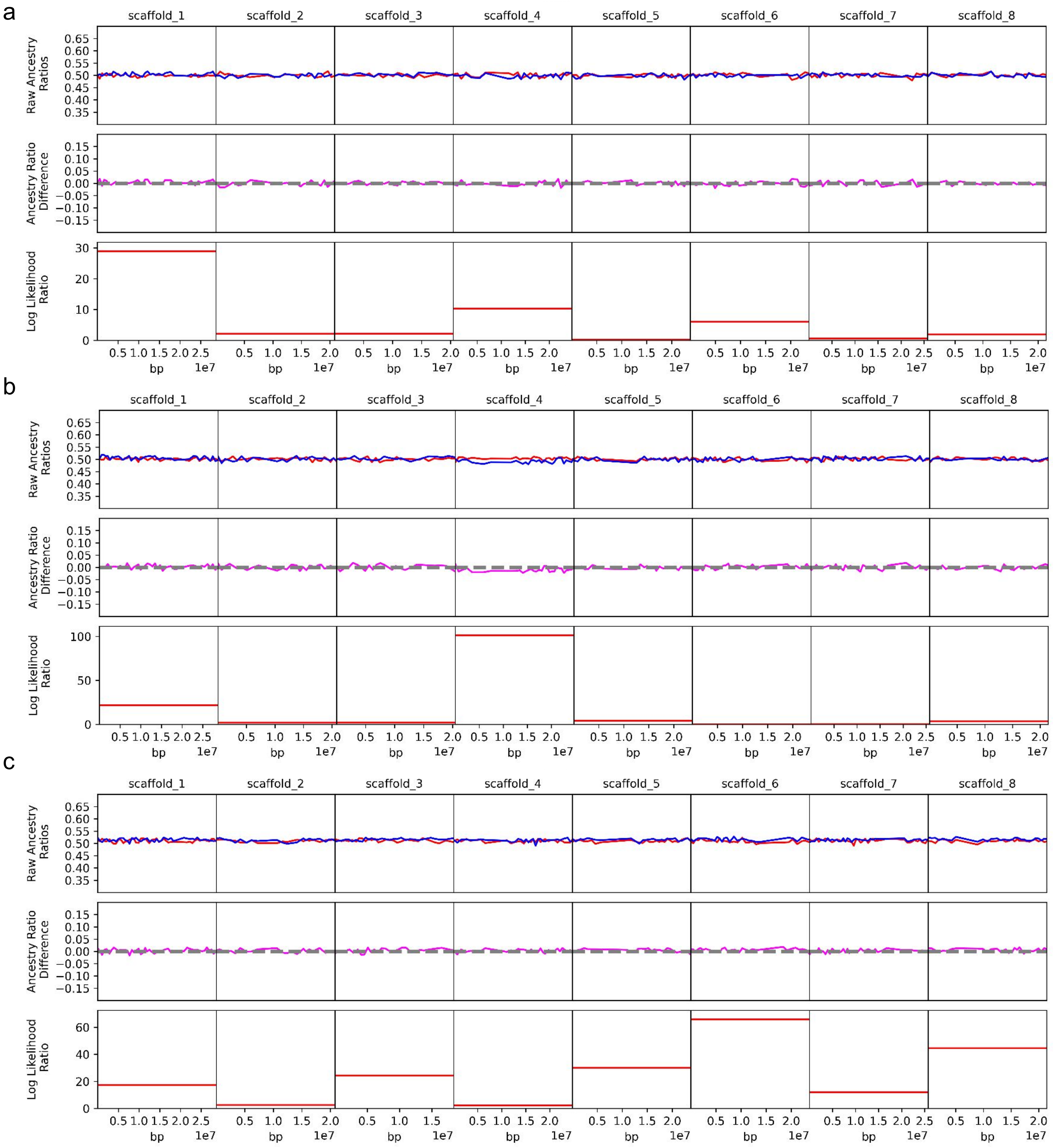


Figure S30. Ancestry ratios obtained from pollen bulk sequencing a,b) two individuals of $F_1$ cross between Central European *A. lyrata* AL62 and Siberian outcrossing *A. lyrata* TE11, c) a cross between NT8 Siberian outcrossing *A. lyrata* and NT1 Siberian selfing *A. lyrata*. No deviations from 50/50 ancestry ratio were observed.

a

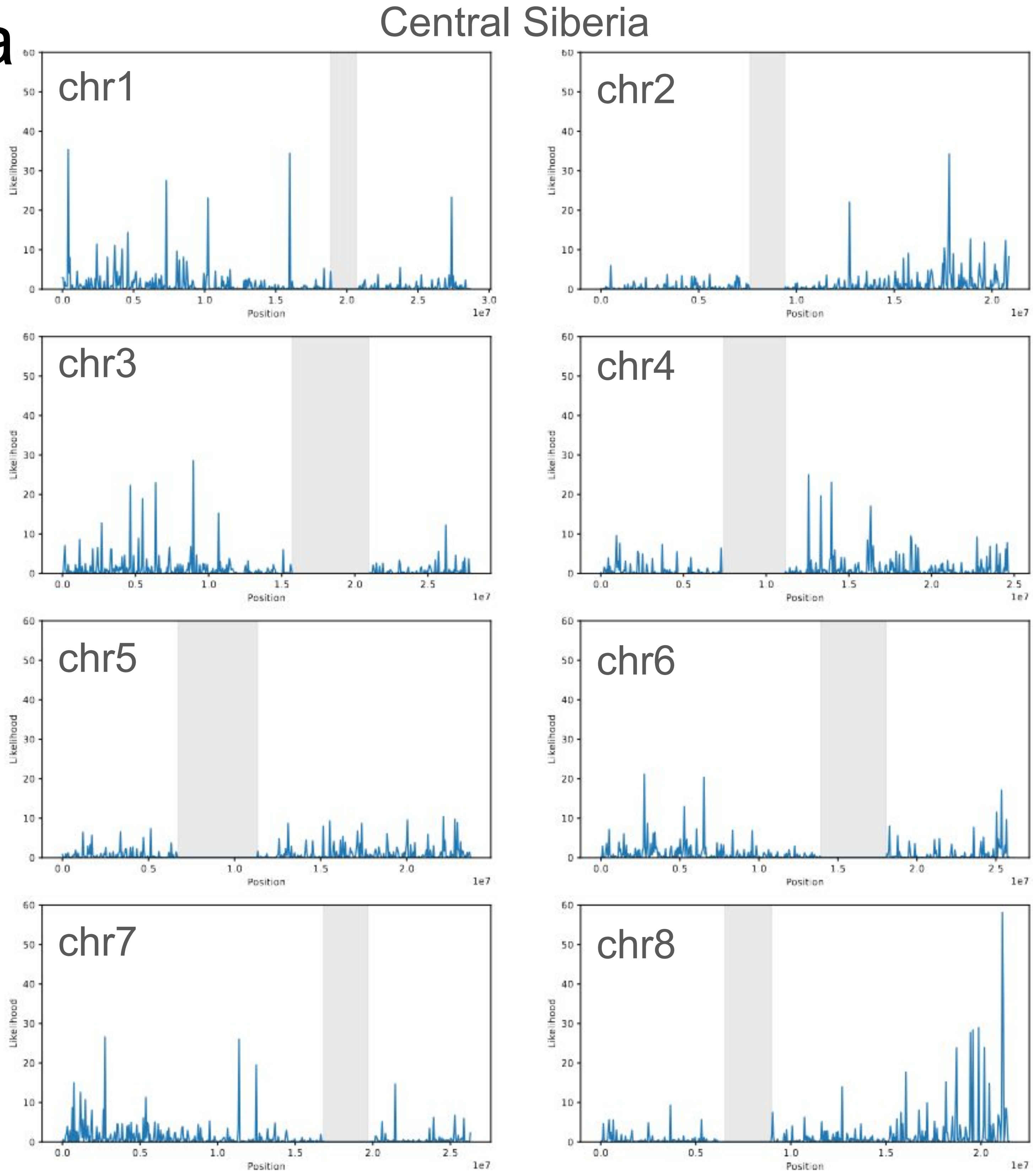

Central Siberia
chr1
chr2
chr3
chr4
chr5
chr6
chr7
chr8
Likelihood
Position
1e7

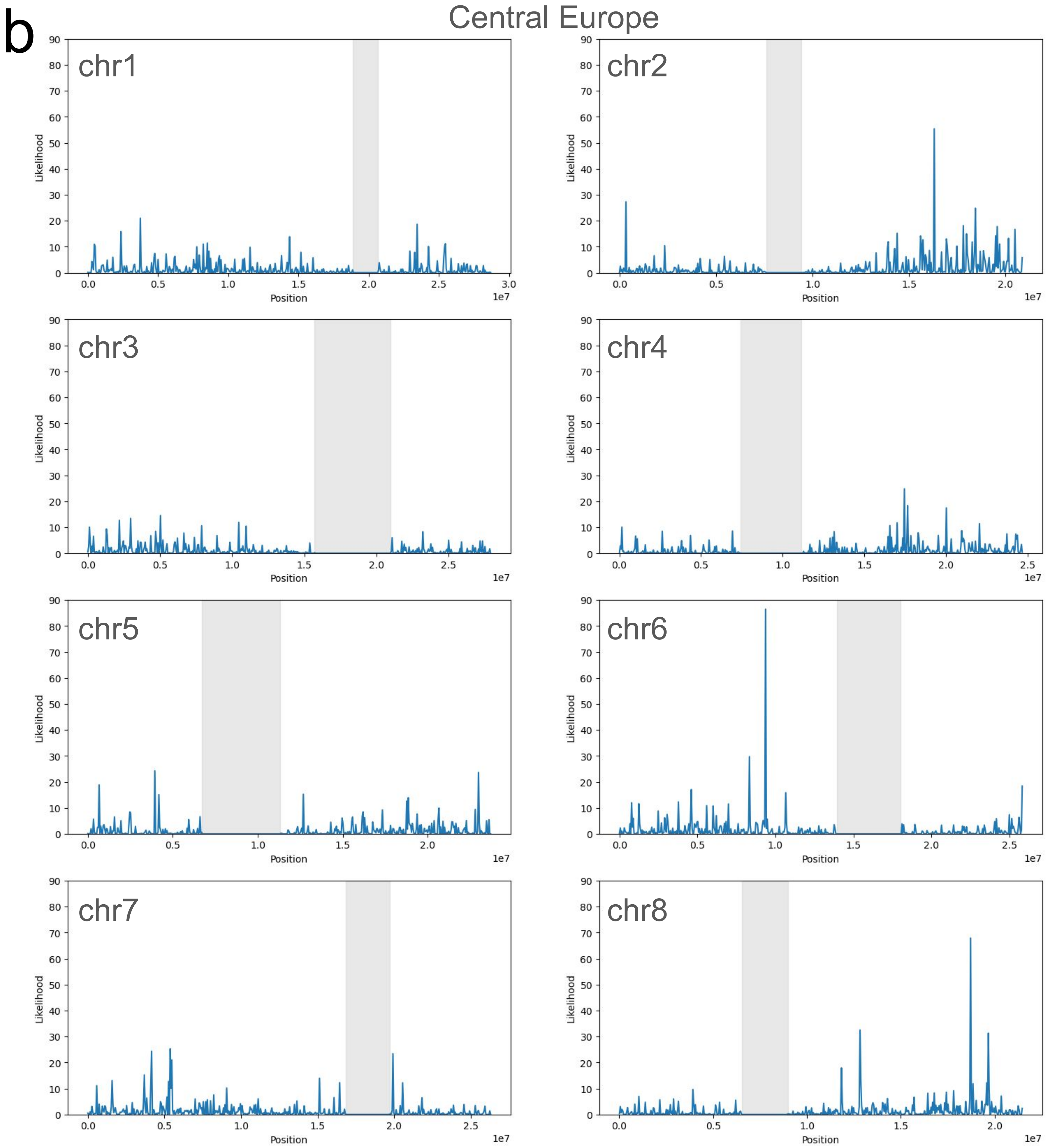

b
Central Europe
chr1
chr2
chr3
chr4
chr5
chr6
chr7
chr8
Likelihood
Position
1e7

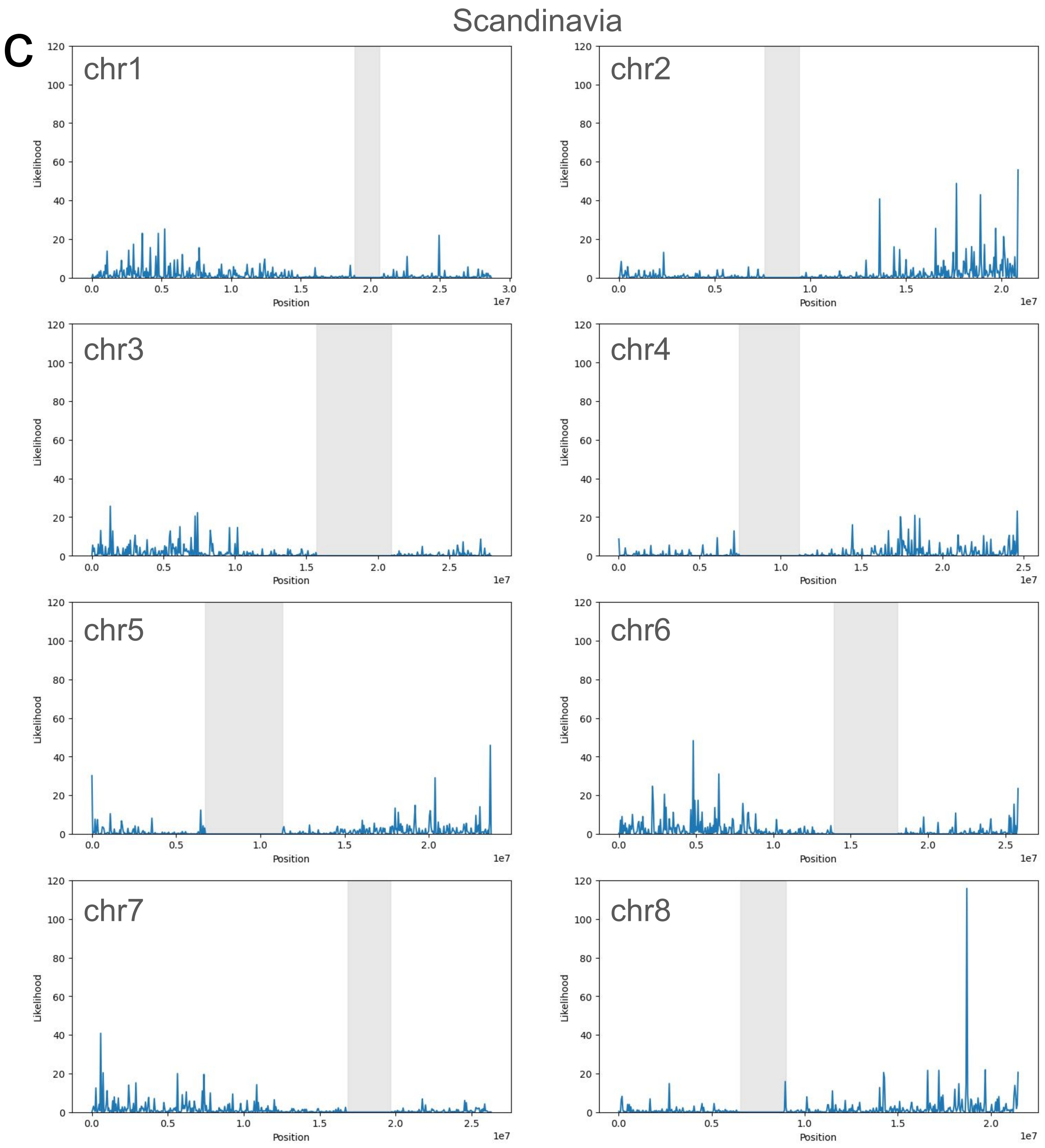
C
Scandinavia
chr1
chr2
chr3
chr4
chr5
chr6
chr7
chr8
Likelihood
Position
1e7

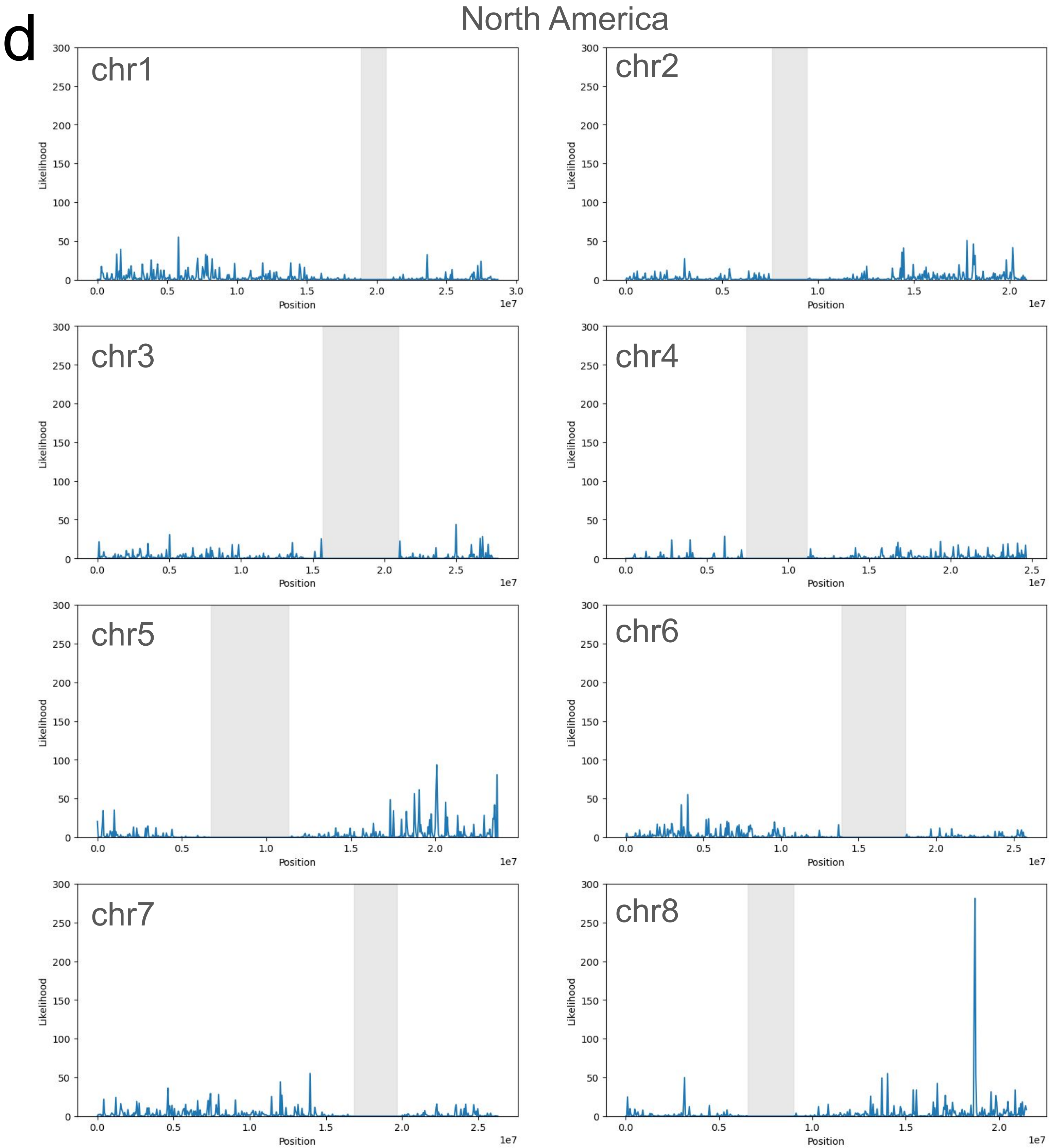


Figure S31. Likelihood values along the chromosomes obtained with SweeD scans of a) Central Siberia population, b) Central Europe population, c) Scandinavian population, d) North American population of *A. lyrata*.

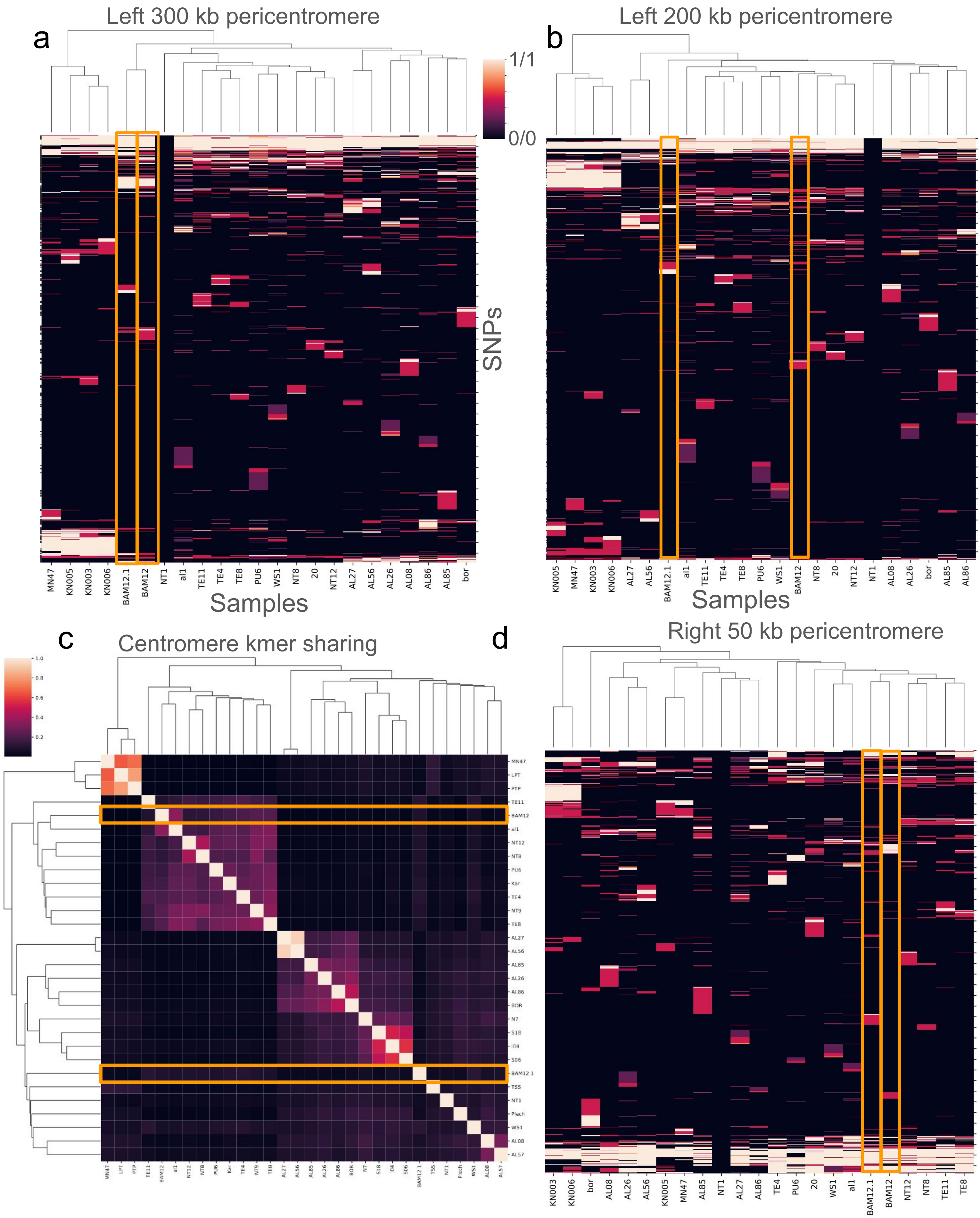


Figure S32. SNP genotype heatmaps in pericentromeres compared with centromere 60-mer sharing on chr6 of *A. lyrata*. Chr6 of BAM12.3 and BAM12.1 (marked with orange rectangles) have different repeat types, so, their centromere haplotypes are very different, however, they are samples from the same location and on chromosome arms they normally have the closest sequences. a,b) Clustermaps of SNPs from the right side of the centromere based on genotyping of long reads mapped to the reference genome. Two BAM12 samples start to cluster differently between 200 and 300 kb from the centromere array edge. c) Centromere kmer sharing on chr6 of *A. lyrata*, d) clustermap of SNPs on the right side of the centromere, BAM12 samples cluster together even in the closest 50 kb to the centromere array edge.
