## Supplementary Tables 1-3 for "The Nature of Centromeric Repeat Turnovers in the genus *Arabidopsis*"

**Supplementary Data 1. Metadata of the genomes.**
**Supplementary Data 2. Heatmap of similarity of 60-mer frequencies between the centromeric arrays.**

| P-val = 0.0007 | AlyCEN179 in centromere | Other repeat in centromere |
|---|---|---|
| AlyCEN179 outside centromere | 159 | 98 |
| No AlyCEN179 outside centromere | 55 | 72 |

| P-val = 0.0012 | AlyCEN168 in centromere | Other repeat in centromere |
|---|---|---|
| AlyCEN168 outside centromere | 96 | 110 |
| No AlyCEN168 outside centromere | 54 | 124 |

| P-val = 1.4e-18 | AlyCEN176 in centromere | Other repeat in centromere |
|---|---|---|
| AlyCEN176 outside centromere | 24 | 47 |
| No AlyCEN176 outside centromere | 1 | 312 |

Chi-squared test

| P-val = 8e-60 | AlyCEN179 outside centromere, centromere of another type | Other repeat outside centromere, centromere of another type |
|---|---|---|
| Yes | 174 | 164 |
| No | 53 | 264 |

Table S1. Repeat types present in and out of the centromeres. Fisher's exact test showed significant association for each repeat type; repeats of the same type are more probable to be present in chromosome arms when the centromere has the same repeat type. Chi-squared test showed significant difference of distribution of AlyCEN179 and other repeat types outside the centromeres. AlyCEN179 is more often present in chromosome arms when the centromere has another repeat type.

| Number of inserted repeats | Number of simulations | Maximum number of repeats of inserted type | Times inserted repeat kept for 1 mln generations |
|---|---|---|---|
| 1 | 100 | 100 | 4 |
| 20 | 100 | 510 | 14 |
| 100 | 100 | 2586 | 44 |

Table S2. The results of forward-time simulations of repeat array evolution with two repeat types.

| | **alpha, whole gene** | **p-adj (BH)** | **alpha, N-tail** | **p-adj, N-tail (BH)** |
|---|---|---|---|---|
| *A. thaliana* | 0.7091 | 1 | 0.5526 | 0.58 |
| *A. cebennensis* and *A. pedemontana*, non-syntenic copy | 0.449 | 1 | NA | |
| *A. cebennensis* and *A. pedemontana*, syntenic copy | NA | 1 | NA | |
| *A. arenosa*, non-syntenic copy | 0.5819 | 0.2 | 0.1836 | 0.58 |
| *A. arenosa*, syntenic copy | 0.173 | 1 | -0.0607 | 0.58 |
| *A. halleri*, non-syntenic copy | 0.2 | 1 | 0.5676 | 0.58 |
| *A. halleri*, syntenic copy | 0.321 | 1 | 0.3616 | 0.58 |
| *A. lyrata*, non-syntenic copy | -0.1843 | 1 | 0.2768 | 0.58 |
| *A. lyrata*, syntenic copy | 0.6806 | 0.2 | 0.625 | 0.58 |

Table S3. Results of MK test with *Capsella* as an outgroup for the whole CENH3 coding region DNA sequence and only N-terminus sequence (81 aa in *A. thaliana* protein).
